# Cryo-EM structures of the tubulin cofactors reveal the molecular basis for the biogenesis of alpha/beta-tubulin

**DOI:** 10.1101/2024.01.29.577855

**Authors:** Aryan Taheri, Zhoaqian Wang, Bharti Singal, Fei Guo, Jawdat Al-Bassam

**Affiliations:** Molecular Cellular Biology epartment, University of California, Davis, CA, USA

**Keywords:** Cryo-EM, alpha/beta-tubulin dimer, tubulin biogenesis, tubulin cofactor, TBCD, TBCE, TBCC, Arl2, microtubules

## Abstract

Microtubule polarity and dynamic polymerization originate from the self-association properties of the a-tubulin heterodimer. For decades, it has remained poorly understood how the tubulin cofactors, TBCD, TBCE, TBCC, and the Arl2 GTPase mediate a-tubulin biogenesis from α- and β-tubulins. Here, we use cryogenic electron microscopy to determine structures of tubulin cofactors bound to αβ-tubulin. These structures show that TBCD, TBCE, and Arl2 form a heterotrimeric cage-like TBC-DEG assembly around the a-tubulin heterodimer. TBCD wraps around Arl2 and almost entirely encircles -tubulin, while TBCE forms a lever arm that anchors along the other end of TBCD and rotates α-tubulin. Structures of the TBC-DEG-αβ-tubulin assemblies bound to TBCC reveal the clockwise rotation of the TBCE lever that twists a-tubulin by pulling its C-terminal tail while TBCD holds -tubulin in place. Altogether, these structures uncover transition states in αβ-tubulin biogenesis, suggesting a vise-like mechanism for the GTP-hydrolysis dependent a-tubulin biogenesis mediated by TBC-DEG and TBCC. These structures provide the first evidence of the critical functions of the tubulin cofactors as enzymes that regulate the invariant organization of αβ-tubulin, by catalyzing α- and β-tubulin assembly, disassembly, and subunit exchange which are crucial for regulating the polymerization capacities of αβ-tubulins into microtubules.

## Introduction

Microtubules (MTs) are polarized cytoskeletal polymers that generate forces through dynamic polymerization at their ends^1-3^, form stable tracks for intracellular cargo transport^4^, and compose the rigid cores of cilia^5^. MTs are assembled by protofilaments of head-to-tail polymerized αβ-tubulin heterodimers. Homotypic lateral interactions between adjacent a- and -tubulins stabilize the tube-like MT structures. The invariant organization of αβ-tubulin, with -tubulin bound on top of a-tubulin, is fundamentally conserved across eukaryotes. Polymerization of a-tubulins promotes GTP-hydrolysis at the -tubulin Exchangeable site (E-site)^1,3,6^. A second Non-exchangeable (N-site) GTP on a-tubulin, sandwiched between a and -tubulins, stabilizes the αβ-tubulin heterodimeric organization^3,7^.

A conserved family of five tu<u>b</u>ulin <u>c</u>ofactors (TBCs A, B, C, D, E) assemble newly folded α- and β-tubulins into αβ-tubulin heterodimers through a GTP-hydrolysis dependent process^8,9,13,14^. These five TBCs were shown to control the soluble a-tubulin concentration in eukaryotic model systems and are attributed to be master regulators of MT polymerization^10-15^. TBCC, TBCD, and TBCE play central roles in a-tubulin assembly or biogenesis, and a conserved GTPase, ADP-ribosylation factor-like 2 (Arl2), regulates their activity ^9,10,16-18^. In contrast, TBCA and TBCB play accessory roles in earlier steps of the biogenesis process^19,20^.

Complete inactivation of TBCs is lethal in many model organisms due to loss of dynamic MTs ^10,13,26,27^. Mutations in the human TBCD, TBCE, and Arl2 are linked to developmental and neurological disorders^21-22^. TBCD mutations are linked to infantile neurodegenerative encephalopathy and corpus callosum hypoplasia^23-26^. TBCE N-terminus mutations are linked to hypoparathyroidism facial dysmorphism (Kenny-Caffey syndrome) disorder and TBCE C-terminus mutations are linked to Giant Axonal motor neuropathy disorder ^27-29^. Mutations in Arl2 are linked to microcornea, rod-cone dystrophy, cataract, and posterior staphyloma (MCRS) syndrome^22^. Common to these disorders are impairments in MT functions during development.

An assembly of TBCD and TBCE has been suggested to promote αβ-tubulin biogenesis by forming a dynamic “super-complex”^12, 30-32^. The release of αβ-tubulin heterodimers from the complex requires TBCC binding to TBCD/TBCE-αβ-tubulin. TBCC activates GTP-hydrolysis and releases a-tubulin, and non-hydrolyzable GTP analogs prevent the a-tubulin release or TBCC dissociation^30^. The mode of TBCC binding is unknown. We previously reconstituted yeast TBCD, TBCE, and Arl2, termed TBCG, revealing their shared roles as heterotrimeric subunits in a stable catalytic platform termed TBC-DEG (Figure S1A)^9,33^. These studies demonstrated that Arl2 was a previously unknown source of GTP hydrolysis in αβ-tubulin biogenesis and a missing subunit of a larger multi-subunit TBC assembly. Arl2 nucleotide-locked mutants lead to severe defects in MT polymerization that are like defects caused by the loss of TBCC, TBCD, and TBCE^12,34^. Therefore, a multi-subunit TBC-DEG platform is the likely enzymatic site of αβ-tubulin biogenesis^9,33,35^.

There is a lack of understanding of the structural basis of TBCD, TBCE, TBCC, and Arl2 organization and αβ-tubulin biogenesis mechanisms. It remains unknown how the TBC-DEG platform assembles αβ-tubulin dimers, and what the roles of TBCC or Arl2 GTP hydrolysis are in αβ-tubulin biogenesis. Furthermore, it is unclear whether the TBC-DEG platform also promotes an alternative pathway that degrades αβ-tubulin heterodimers, and what role TBCC might play in that mechanism. Here, we present two 3.6-A cryo-EM structures for the yeast TBC-DEG-a-tubulin (binary) and two 3.6-A TBC-DEG/TBCC-a-tubulin (ternary) assemblies revealing their intricate cage-like organization around a-tubulin, extensive αβ-tubulin binding interfaces, and TBCC mediated transitions in regulating αβ-tubulin organization. Our structural studies identify molecular transitions in the TBC-DEG-a-tubulin complex driven by TBCC binding and develop a new foundation for understanding how αβ-tubulins are assembled and degraded.

## Results

### Cryo-EM Structures of TBC-DEG-αβ-tubulin

We purified recombinant yeast TBC-DEG and reconstituted it with αβ-tubulin to form 1:1 TBC-DEG-αβ-tubulin (binary) assemblies as previously described (Figure S1)^33^. Initial cryo-EM sample preparation and imaging showed these assemblies aggregate, likely due to self-assembly of αβ-tubulin (not shown). To improve sample homogeneity, we utilized a DARPin variant, ΔN-DARPin, which binds β-tubulin’s polymerization interface and prevents αβ-tubulin self-association between TBC-DEG assemblies^36^. ΔN-DARPin binding did not alter αβ-tubulin to TBC-DEG binding stoichiometry (Figure S1). 5ingle particle analysis of the TBC-DEG-αβ-tubulin-ΔN-DARPin revealed homogenous cone-shaped particles with clear secondary structure details (Figure S2). We determined cryo-EM structures for two states of the TBC-DEG-αβ-tubulin assemblies using a single particle reconstruction pipeline detailed in Material and Methods (Table 1; Figure S2-S4). The approach led to a consensus core TBC-DEG-αβ-tubulin structure at 3.6-A resolution and two unique states for TBCE that exhibit domain reorganization and differ by a 30° rotation around α-tubulin (Figure 1, Figure S3-S4, and materials and methods)^37^. We modeled the core TBC-DEG-αβ-tubulin assembly and two states of the lever arm-like extensions of TBCE (Table 1; Figure S4).

**Table 1:** Cryo-EM Data Collection, Single particle processing, refinement, model building, and validation Statistics.

| **Data collection** | TBC-DEG-$\alpha\beta$-tubulin | | TBC-DEG/TBCC-$\alpha\beta$-tubulin | |
| --- | --- | --- | --- | --- |
| Facility | UC Berkeley |  | UC Davis |  |
| Microscope | Titan Krios |  | Glacios |  |
| Detector | K2 (Gatan) |  | K3 (Gatan) |  |
| Magnification | 57,000 |  | 45,000 |  |
| Voltage (Kv) | 300 Kv |  | 200 Kv |  |
| Electron exposure (e/$\text{\AA}^2$) | 50 | | 60 | |
| Defocus Range ($\mu\text{m}$) | -0.9 to -2.1 | | -0.7 to -1.7 | |
| Pixel Size ($\text{\AA}$) | 0.662 | | 0.440 | |
| Movie stacks | 5145 |  | 21542 |  |
| **Data processing** | TBC-DEG-$\alpha\beta$-tubulin state 1 | TBC-DEG-$\alpha\beta$-tubulin state 2 | TBC-DEG/TBCC-$\alpha\beta$-tubulin state1 | TBC-DEG/TBCC-$\alpha\beta$-tubulin state 2 |
| **Map Refinement** | EMDB XXX | EMDB XXX | EMDB XXX | EMDB XXX |
| Final Particle images | 275,487 | 275,487 | 87,013 | 102,987 |
| Map Resolution ($\text{\AA}$) | 3.70 | 3.70 | 3.83 | 3.73 |
| FSC threshold | 0.143 | 0.143 | 0.143 | 0.143 |
| Map Resolution range ($\text{\AA}$) | 3.5-7 | 3.5-7 | 3-6.5 | 3-6.5 |
| **Model composition** | PDB ID XXX | PDB ID XXX | PDB ID XXX | PDB ID XXX |
| Chains | 6 | 6 | 7 | 7 |
| Atoms | 20338 | 20338 | 22934 | 23007 |
| Residues | 2681 | 2681 | 2858 | 2877 |
| Nucleotides | 3 | 3 | 3 | 3 |
| FSC threshold | 0.5 | 0.5 | 0.5 | 0.5 |
| Model resolution ($\text{\AA}$) | 4.1 | 4.1 | 4.1 | 4.2 |
| Model sharpening B factor ($\text{\AA}^2$) | Phenix<br>Autosharpen | Phenix<br>Autosharpen | Phenix<br>Autosharpen | Phenix<br>Autosharpen |
| **Validation** |  |  |  |  |
| MolProbity Score | 2.28 | XXX | 2.28 | 2.18 |
| Clash score | 19.03 | XXX | 22.46 | 26.02 |
| Rotamer Outliers ( %) | 0.05 | XXX | 0.84 | 0.63 |
| Ramachandran (favored) (%) | 91.37 | XXX | 96.52 | 97.33 |
| Ramachandran (allowed) (%) | 8.63 | XXX | 3.48 | 2.64 |
| Ramachandran (disallowed) (%) | 0 | XXX | 0 | 0 |

**Figure 1:**
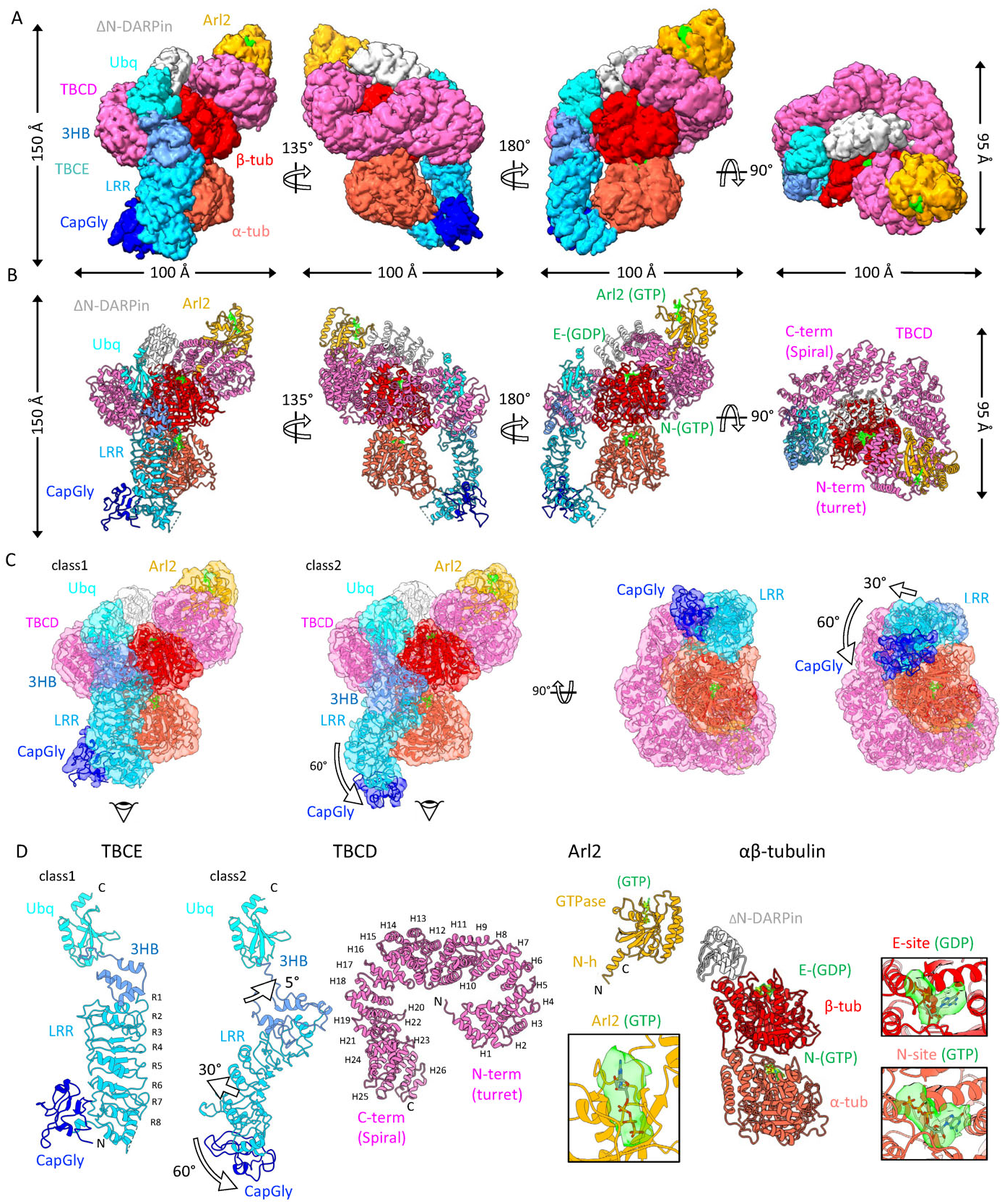
Cryo-EM structures of TBC-DEG-αβ-tubulin (binary) assembly. A) Segmentation of individual subunits of the cryo-EM reconstruction of TBC-DEG-αβ-tubulin-ΔN-DARPin following the color scheme in FigS1B and labeled. B) Display of atomic models of TBC-DEG-αβ-tubulin-ΔN-DARPin. C) Two refined cryo-EM reconstructions with fitted atomic models (ribbon). Class 1 represents a retracted state of TBCE. Class 2 represents the TBCE LRR-α-tubulin bound state. The two states differ in TBCE domain organization described in Figure 1F. D) TBC-DEG-αβ-tubulin subunit models. On the left, two TBCE states are shown with annotated rotations and translations in the 3HB (3-helix bundle), LRR (leucine-rich Repeat), and CapGly domains indicated by arrows. Second from the left, a top view of TBCD with its twenty-six HEAT repeats, N-term (TBCD turret), and C-term (TBCD Spiral) domains labeled. Third from the left, Arl2 (orange) is shown with its N-helix (N-h) and GTPase (GTPase) domains labeled. Below, a close-up view of Arl2 GTP nucleotide density (red) and its model (orange) are displayed. Fourth from left, the αβ-tubulin (tomato/red) bound to ΔN-DARPin (Gray). To its right, electron densities for N-site GTP and E-site GDP with their models are displayed.

### Organization of TBC-DEG-αβ-tubulin

The 3.6Å cryo-EM TBC-DEG-αβ-tubulin structures reveal a 100Å X 95Å X 140Å TBC-DEG cage-like assembly that encases a single αβ-tubulin (Figure 1A-B). TBCD is a 100-A ring composed of twenty-six α-helical HEAT repeats (H1-H26) that curl laterally around the Arl2 N-terminal-helix and almost encircle β-tubulin with its full length. At its C-terminal end, TBCD is bound to the C-terminal Ubq domain of the 100-Å long TBCE. We term the TBCD N-terminal region that binds the Arl2 GTPase the TBCD turret, and its C-terminal region that binds the TBCE Ubq domain the TBCD spiral (Figure 1B-D). Arl2 is placed on top of the TBCD turret above β-tubulin and TBCE is positioned under the TBCD spiral and alongside αβ-tubulin (Figure 1A-B). TBCE is composed of its N-terminal CapGly domain, a central arm-like LRR domain with eight LRR repeats (LRR1-8), a junction comprising a three-helix bundle (3HB), and a C-terminal Ubq domain anchored against the TBCD C-terminal region (Figure 1A-B, 1D). The two differing TBCE states exhibit reorganization of the CapGly and LRR domains (Figure 1C; Figure 1D; Figure S4F-G). In the first state (class 1), the TBCE 3HB and LRR are oriented vertically. TBCE is retracted from binding α-tubulin and the CapGly resides horizontally to the LRR (Figure 1E, left; Figure 1F; Figure S4F-G). In the second state (class 2), the LRR domain is rotated 30° clockwise relative to class 1 through a hinge-like rotation in the 3HB along the TBCD spiral. In this state, the CapGly is repositioned at the tip of the LRR (Figure 1C; Figure S4F-G; Figure 1D).

### TBCD is the scaffold of the TBC-DEG assembly

TBCD is a scaffolding protein that organizes the TBC-DEG assembly through extensive interactions with Arl2, TBCE, and αβ-tubulin. The TBCD turret forms a 60Å diameter solenoid through H1-13 (Figure 1F). The TBCD spiral forms a crescent around -tubulin through H14-26 (Figure 1D). The TBCD turret encircles the Arl2 N-h and binds the base of the Arl2 GTPase (Figure 2). The Arl2 N-h helix is encased through lateral packing by the B-helices of H1, H2, H7, H8, and H9 (Figure 2A-B, E-F). Conserved residues in the intra-HEAT turns of TBCD H1, H2, H3, H5, H6, H8, and H10 bind residues at the lower surface of the Arl2 GTPase through ionic and hydrophobic interactions (Figure 2A, E-F; Figure S14D-E; Figure S14D-E, G, I-J). Conserved TBCD Lys and Arg residues bind conserved Arl2 Asp, Gln, and Glu residues (Figure S11, S14 D-E, G; Figure S14D-E). Hydrophobic packing of conserved Leu, Phe, and Trp residues in TBCD and Arl2 are interspersed amongst the ionic interactions (Figure 2E-F; Figure S14D-E, G; Figure S161-J).

**Figure 2:**
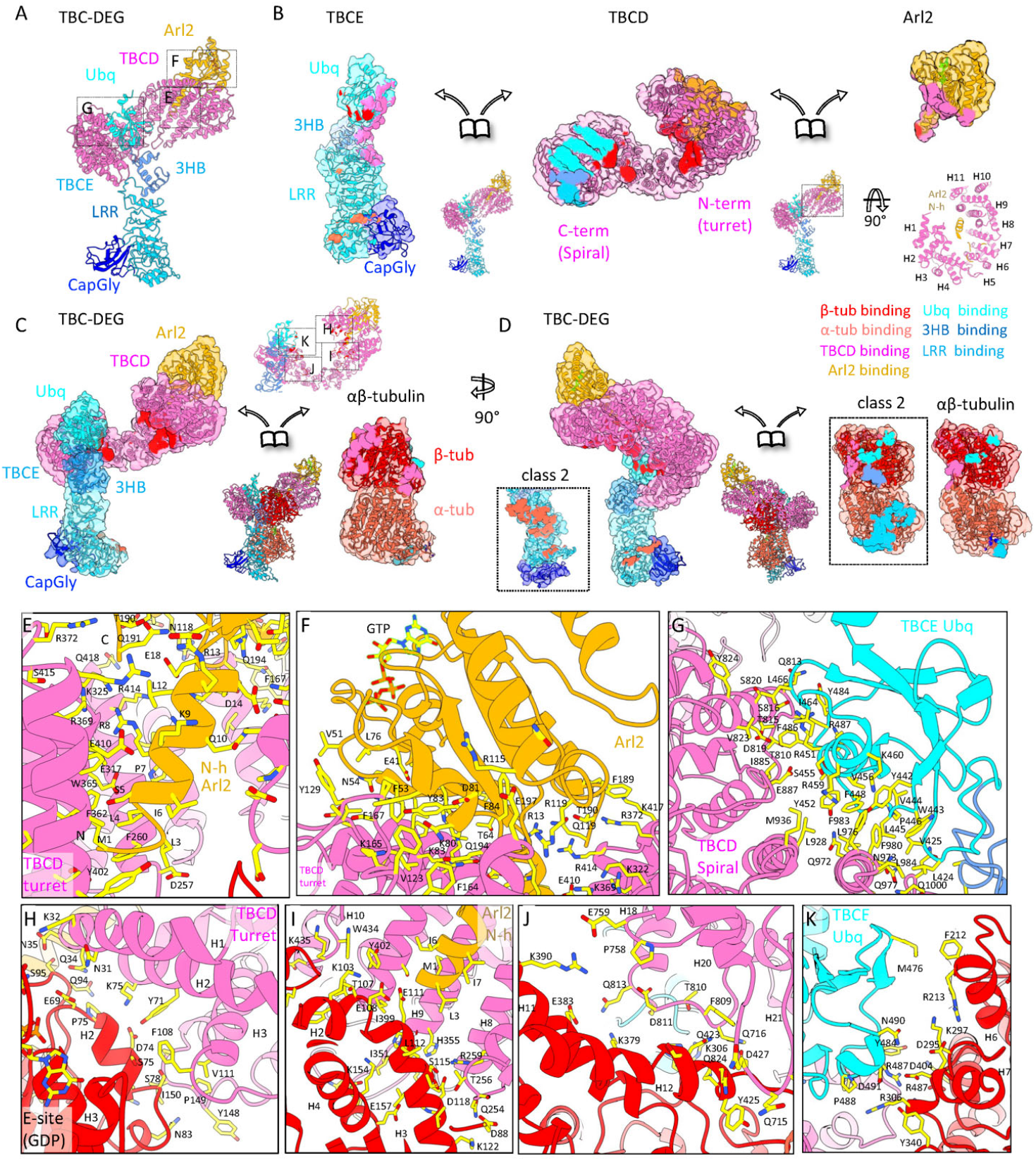
The TBC-DEG cage-like assembly organization and interfaces with αβ-tubulin. A) Model for TBC-DEG assembly without αβ-tubulin. Boxes mark the interacting interfaces shown in panels E, F, G. B) Disassembled view of the TBC-DEG. TBCD is in the center, TBCE on the left, and Arl2 on the right. Interaction interfaces are colored by the footprints of the interacting subunits (TBCD: TBCE Ubq: cyan, TBCE LRR: Sky blue, Arl2: orange, β-tubulin: red). At the bottom left, a close-up view of the Arl2 N-h interface with TBCD turret HEAT repeats is displayed. C) Disassembled view of the TBC-DEG-αβ-tubulin. The TBC-DEG-αβ-tubulin model is in the center, TBC-DEG on the left, and αβ-tubulin on the right. The interaction interfaces are colored the same as Fig2B. αβ-tubulin model shows four TBC-DEG binding zones to β-tubulin. The top atomic inset shows Boxes marking the interfaces with β-tubulin shown in panels H, I, J, and K. D) Rotated view of TBC-DEG compared to panel F. The box shows TBCE and αβ-tubulin interfaces in Class 2. To its right, class 1 TBCE interfaces on αβ-tubulin are shown. E) View of Arl2 N-h (orange) and TBCD turret (pink) interface. F) View of Arl2 GTPase (orange)-TBCD turret (pink) interface. G) View of the TBCE Ubq (cyan) -TBCD spiral (pink) interface. H) TBC-DEG interface I on β-tubulin. TBCD turret (pink) interacts with β-tubulin (red). This interface involves TBCD HEAT 1-3 helices binding β-tubulin H2 and H3. I) TBC-DEG interface 2 on β-tubulin. TBCD turret (pink) HEAT 8-10 and N-h of Arl2 (orange) interaction with β-tubulin (red) H2 and H3 is shown. J) TBC-DEG interface 3 on β-tubulin. TBCD (pink) HEAT 18-20 interacting with β-tubulin (red) H11, H12, and C-terminal tail is shown. K) TBC-DEG interface 4 on β-tubulin. TBCE Ubq (cyan) bound on the TBCD spiral (pink) interacting with β-tubulin (red) H6 and H7 is shown.

The TBCD spiral binds the TBCE Ubq at its C-terminal end. The TBCE Ubq is a ubiquitin-like fold domain that binds the B-helices of TBCD repeats H18, H20, H23, and H25 (Figure 2A-B, G). This interface is composed mostly of hydrophobic interactions between conserved TBCD and TBCE Phe, Trp, and Tyr residues (Figure 2G, Figure S11, S12, S14C-F; S16). TBCD H26 binds the TBCE 3HB and is stabilized by interactions between the TBCE Ubq-TBCD spiral interface, creating a likely pivot point for the rotation of TBCE between its two states (Figure 2A-B, 1D). Both the TBCD H26 and the TBCE 3HB are conserved across TBCD and TBCE orthologs, supporting the functional importance of this interaction (Figure S14C-D).

The TBC-DEG-αβ-tubulin structures reveal that the N and C-termini of TBCD, Arl2, and TBCE are either inaccessible or involved in forming critical interfaces (Figure S15, S16). Truncating these termini or fusing them to fluorescent protein tags would likely be deleterious to TBC-DEG assembly. Modifying the termini of TBCD, TBCE, and Arl2 with fusion proteins at their N-or C-termini may have caused destabilization of native TBC-DEG assemblies in previous studies, explaining why imaging of these proteins has failed to capture TBC-DEG assemblies^35,38^. Our recombinant reconstitution studies using a short purification tag along the TBCD N-terminus have proven to be the only way to capture intact TBC DEG assemblies^33^.

### TBC-DEG deforms αβ-tubulin heterodimers

The TBC-DEG cage forms interfaces with both α- and β-tubulin that encompasses regions known to be involved in interactions between αβ-tubulin heterodimers or interactions with microtubule-associated proteins (Figure 2C-D). Both the α- and β-tubulin C-terminal tails are bound to TBC-DEG domains and are occluded in the structures (Figure 2C-D). The TBCD turret binds β-tubulin at two sites and the TBCD spiral binds a third site (Figure 2C-D, H-J). The extreme TBCD N-terminus forms the first β-tubulin binding site (site 1; Figure 2H). This binding site is mediated through ionic and hydrophobic interactions between TBCD turret H1, H2, and H3 and the β-tubulin H2 and H3 helices (Figure 2H; Figure S14H). The TBCD and Arl2 N-h interface forms the second binding site to β-tubulin (site 11; Figure 21; Figure S141). This binding site is mediated by TBCD H8, H9, and H10 intra-repeat turns, and the extreme N-terminus of the Arl2 N-h binding to the β-tubulin microtubule luminal facing helices H3 and H4 near its GDP occupied E-site. The third TBCD β-tubulin binding site is mediated by H18 and H20 in the TBCD spiral domain binding the β-tubulin C-terminal H11 and H12 helices and C-terminal tail (site 111; Figure 2J; Figure S14J). The TBCE-Ubq is positioned by the TBCD spiral to form the fourth binding site, completing a circular set of interfaces around β-tubulin (site 1V; Figure 2K; Figure S14K). This site is near the β-tubulin microtubule plus-end oriented H6 and H7 helices and M-loop. Collectively these interfaces cover nearly all the polymerization surfaces of β-tubulin (Figure 2F-G). TBCD regions in between these binding sites block access to β-tubulin (Figure 2C-D, H-K; Figure S16A, F).

The TBCE LRR-CapGly extends along α-tubulin covering additional lateral polymerization sites of α-tubulin at its H10 helix near its MT forming interface known as the MT loop (Figure S4E-F). TBCE LRR-CapGly extends longitudinally along αβ-tubulin and swivels between two positions (Figure 1E-F). 1t either engages the α-tubulin lateral polymerizing interface through its LRR (class2), or it engages the α-tubulin C-terminus through its CapGly while the LRR retracts from engaging α-tubulin (class1) (Figure 1F, S4E-F). TBC-DEG binding to αβ-tubulin induces a 4°-clockwise rotation of α-tubulin with respect to the α-tubulin orientation in soluble αβ-tubulin. This deformation is likely caused by the TBCE LRR-CapGly binding to α-tubulin.

TBC-DEG binding fully occludes the α-tubulin and β-tubulin C-termini, β-tubulin N-terminus, but not the α-tubulin N-terminus. The structures reveal the very tight binding footprint of TBC-DEG onto these α- and β-tubulin surfaces and termini, which may explain why only N-term fusions on α-tubulin can be stable *in vivo*^12,39,40^.

### Cryo-EM Structures of TBC-DEG/TBCC-αβ-tubulin

We reconstituted the recombinant TBC-DEG with TBCC and αβ-tubulin to form TBC-DEG/TBCC-αβ-tubulin (termed ternary) assemblies for single particle cryo-EM structure determination^33^. TBCC only binds TBC-DEG when it is bound to αβ-tubulin^33^. A high-affinity TBCC interaction requires an Arl2 Q73L mutant TBC-DEG, leading the Arl2 to a GTP-locked state, or by the addition of GTPγS (Figure S5A)^33^. By incubating with S mM GTPγS, stoichiometric TBC-DEG-Arl2-Q73L, αβ-tubulin, and TBCC, ternary assemblies were reconstituted using size exclusion chromatography^33^, as described in materials and methods. Initial cryo-EM imaging revealed ternary assemblies also suffer from severe aggregation. However, when using ΔN-DARPin to block αβ-tubulin, TBCC binding stoichiometry decreased, suggesting a potential steric hindrance between ΔN-DARPin and TBCC within these assemblies (see below). So, to prevent aggregation without interfering with TBCC binding, we utilized the α-rep iHS, which binds the opposite end of αβ-tubulin compared to ΔN-DARPin (Figure S5D)^41^. In contrast to ΔN-DARPin, iHS binding did not decrease TBCC stoichiometry within ternary assemblies and led to homogenous particles (Figure S5-S6).

We determined multiple single-particle cryo-EM structures of the TBC-DEG/TBCC-αβ-tubulin as detailed in Material and Methods (Figure S6). A consensus 3.6-A core structure of TBC-DEG/TBCC-αβ-tubulin was resolved, and an arm-like extension dissociated from the core of the assembly was attributed to TBCE-LRR-CapGly (Figure S6). The dissociated conformation of the TBCE-LRR-CapGly contrasts with its bound conformation in the TBC-DEG-αβ-tubulin assemblies (Figure S6). Multiple interspersed regions of the determined structure displayed blurred features which we suspected to be averaged continuous heterogeneity (Figure S6). To improve the clarity of these regions, we performed 3D Variability Analysis (3DVA) in Cryosparc^37^ (Figure S7A-D). Comparisons between conformations of domains observed across 3DVA components allowed us to draw relationships between TBCC-N domain bindings to TBCD and TBCE positions (Figure S7C). We identified TBCC bound versus unbound variations (Figure S7A); two conformations of the TBCD C-term/TBCE-Ubq that differ by a S° shift correlating to TBCC-N binding (Figure S7B); and two variations of TBCE binding that revealed when TBCC-N binds TBC DEG, TBCE-LRR-CapGly rotates downward (Figure S7C). Two primary conformations for the TBCE arm orientation were identified (Figure S7D). Two composite maps were generated for the distinct states of the ternary assemblies based on subunit binding relationships (Figure S8).

### TBCC binds Arl2, TBCD and engages αβ-tubulin N-site

The two composite TBC-DEG/TBCC-αβ-tubulin cryo-EM structures reveal the organization of TBCC binding to TBC-DEG-αβ-tubulin (Figures 3, S8D, S9, S10). The two ternary cryo-EM structures differed by the binding of TBCC-N to the central region of TBCD. TBCC consists of a conserved N-terminal α-helical bundle (TBCC-N; residues 1-75), a partially conserved linker region (TBCC-L; residues 76-100), and a conserved C-terminal β-helix domain (TBCC-C; residues 100-270) (Figure S5B; Figure 3A-B). TBCC-N binding to the TBC-DEG-αβ-tubulin core induces conformational changes in the TBCD spiral domain, TBCE-LRR-CapGly arm, and αβ-tubulin (Figure 3C-D). The TBCC-C β-helix binds Arl2 GTPase, and that binding is homologous to the previously resolved Retinitis Pigmentosa 2 (RP2)-Arl3 structure (Figure 3C-D; Figure 4)^42^. The TBCC-C β-helix N-terminal end interacts with the TBCD turret (Figure 3C-D, Figure 4). This suggests that TBCC-C binds Arl2 when it is assembled with TBCD (Figure 4A, C). Two ordered sections of TBCC-L are in proximity to the TBCC-N and TBCC-C regions and bind along the TBCD spiral and turret regions respectively (Figure 3A-B, Figure 4). The most conserved region in the TBCC sequence is in the TBCC-L immediately C-terminal to the TBCC-N α-helical bundle (Figure S13). This region binds across and in between TBCD repeats H11 and H12, extending underneath TBCD to connect the TBCC-N α-helical bundle (Figure 4A-B). The central region of TBCC-L is poorly conserved and is not observed in these structures. The TBCC-N three-helical bundle binds underneath the central section of the TBCD between the second and third TBCD-β-tubulin binding sites (Figure 4A-B). The short end of the TBCC-N three-helical bundle binds directly to the αβ-tubulin intradimer interface, primarily contacting α-tubulin (Figure 4A-B). The TBCC-N bundle is wedged against the α-helices of TBCD H12 repeat and lies 8A away from the γ-phosphate of the N-site GTP (Figure 3, 4A-B, E). The TBC-DEG/TBCC-αβ-tubulin cryo-EM structures and models reveal the complex and intricate interactions of TBCC domains across TBC-DEG and αβ-tubulins bridging 50 A distance between the Arl2 GTPase with αβ-tubulin intradimer N-site GTP.

**Figure 3:**
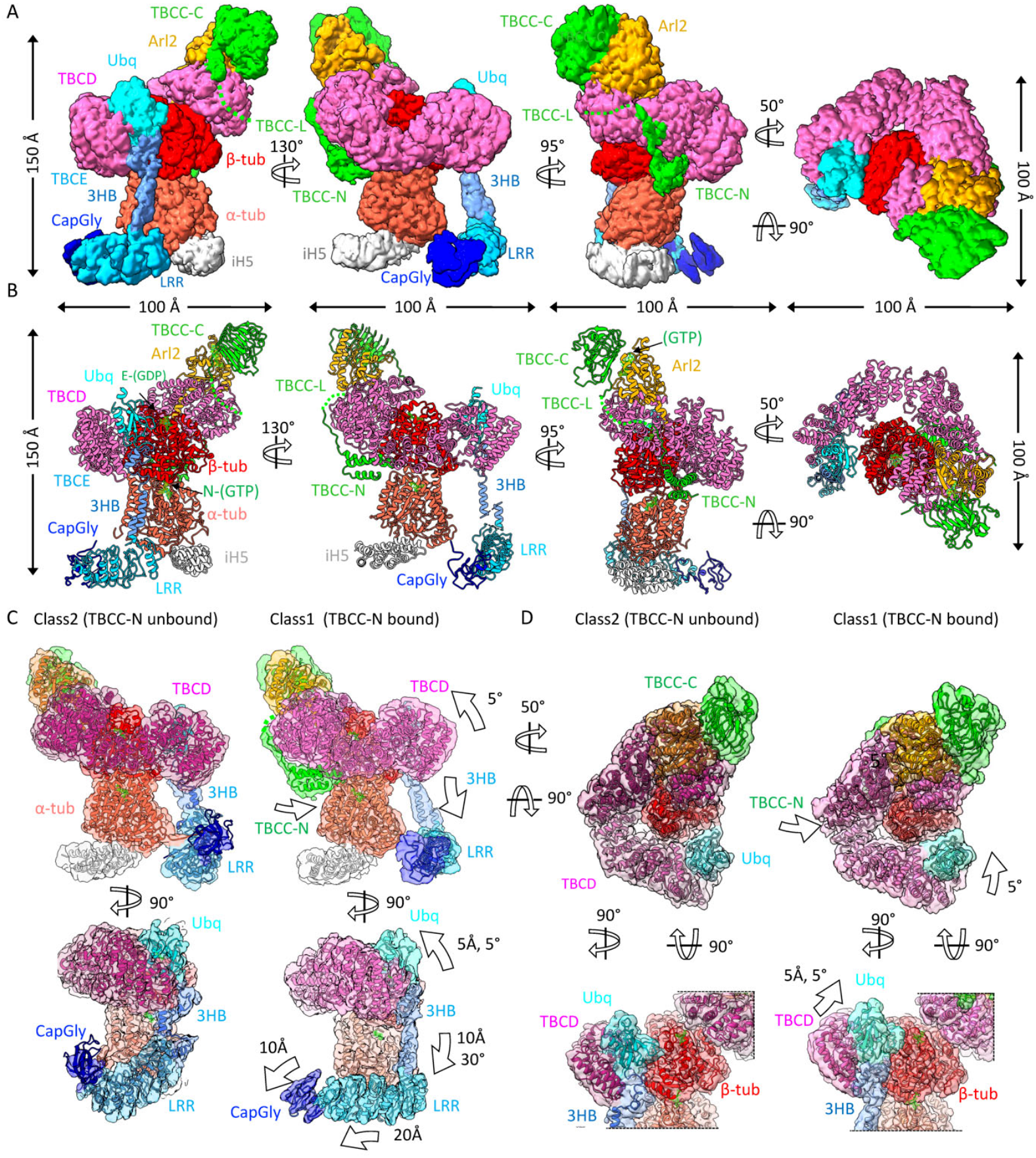
Cryo-EM structures of the TBC-DEG/TBCC-αβ-tubulin (ternary) assembly reveal the catalytic transitions induced by TBCC binding. A) 3.6 Å Cryo-EM segmented maps of TBC-DEG/TBCC-αβ-tubulin-iH5 assembly shown in different views. Different domains are colored based on the scheme shown in Figure S5B and as labeled. B) Atomic models of TBC-DEG/TBCC-αβ-tubulin-iH5 are shown in identical views to C. Subunits, domains, and elements are labeled. C) Views for refined cryo-EM maps of two classes (transparent) with atomic models (ribbon) representing two distinct TBC-DEG/TBCC-αβ-tubulin states. Top left, class 2 shows TBCC-N (green) not bound. Top middle, TBCE class 1 shows TBCC-N (green) is bound to the TBC-DEG-αβ-tubulin core. Top right, overlay of class 1 and class 2 atomic models. Bottom, views of 90Δ rotated slice views of class 2 (left) class 1(middle), and their overlay (right). D) Top, top-end views of TBC-DEG/TBCC-αβ-tubulin classes: Class 2 (top left), class 1 (top middle), and model overlay (top right). Bottom, views of TBCD-spiral and TBCE Ubq domain transitions of TBC-DEG/TBCC-αβ-tubulin classes.

**Figure 4:**
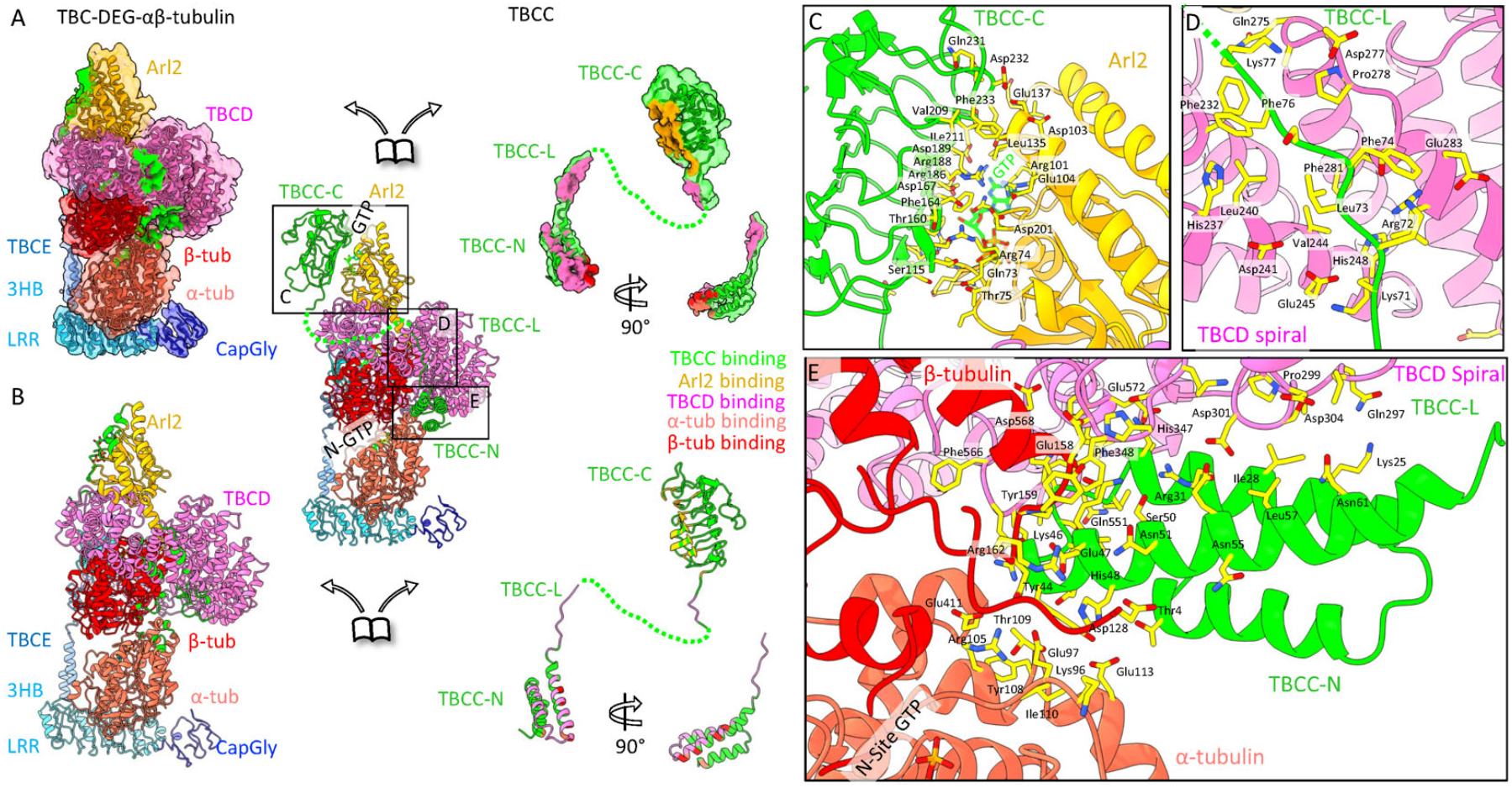
The interfaces of TBCC with TBC-DEG-αβ-tubulin. A) TBC-DEG/TBCC-αβ-tubulin map disassembled showing the TBC-DEG-αβ-tubulin core (left panel) and TBCC-C, TBCC-L, and TBCC-N domains (right panel) with their interface model (middle central panel). Interaction interfaces are colored by the footprints of the interacting subunits (TBCD: TBCE Ubq: cyan, TBCE LRR: Sky blue, Arl2: orange, β-tubulin: red). Three critical interaction interfaces are highlighted by boxes I, 2, and 3. B) Disassembled view of the TBC-DEG/TBCC-αβ-tubulin model. C) TBCC-C-Arl2 interface shown. TBCC-C β-helix and the Arl2 are shown in stick format with the GTP nucleotide. D) TBCC-L and TBCD spiral domain interface shown revealing the aromatic (Phe/Tyr) and positively charged Lys/Arg of TBCC-L interacting with acidic Glu/Asp residues on TBCD. E) TBCC-N interface with TBCD spiral and αβ-tubulin. Residues of TBCC-N three α-helix bundle shown within 10 A proximity of the αβ-tubulin N-site.

**Figure 5:**
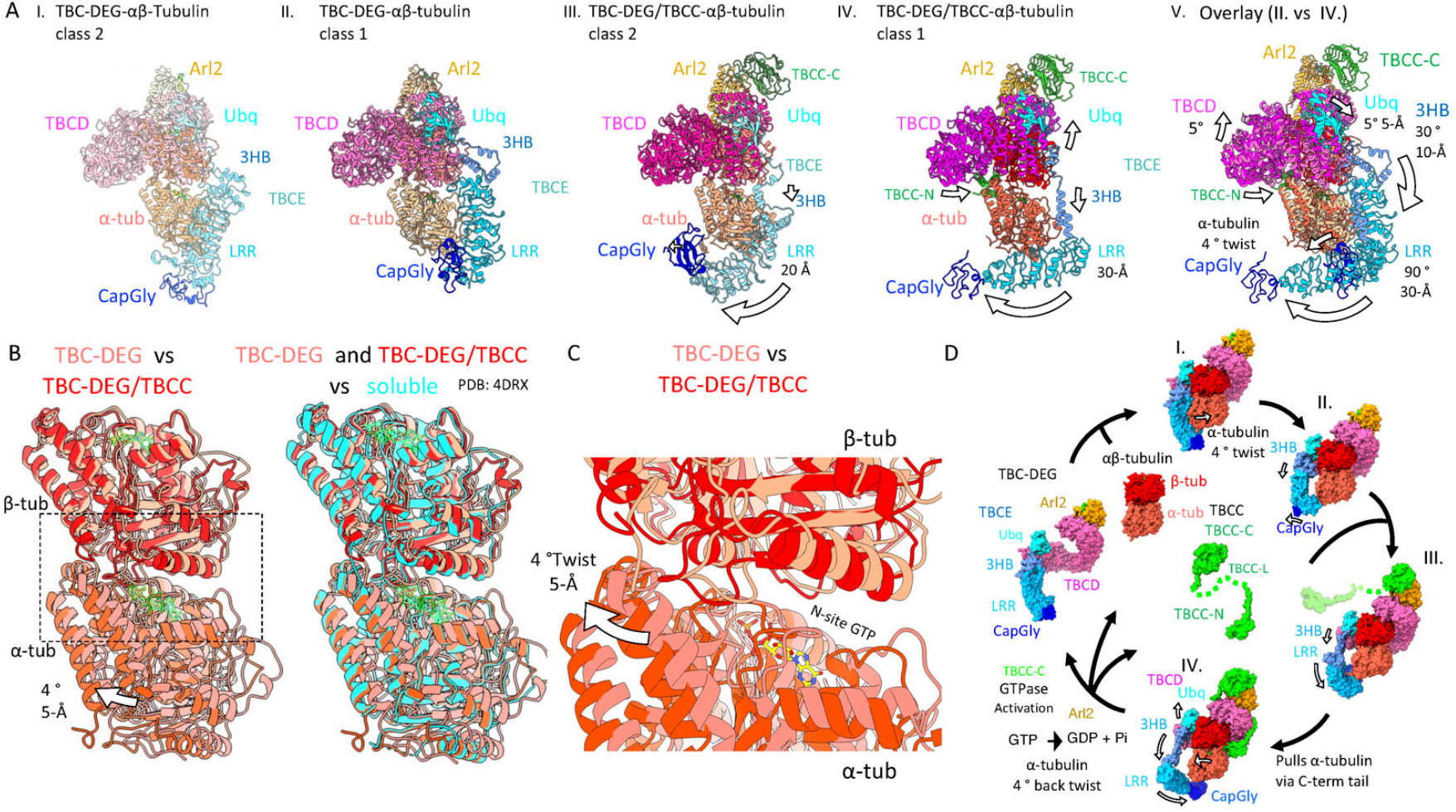
Conformational transition of TBC-DEG and their impact on αβ-tubulin organization. A) I. State I (TBC-DEG-αβ-tubulin class 2), II. State II (TBC-DEG-αβ-tubulin class 1) III. State III (TBC-DEG/TBCC-αβ-tubulin, class 2), IV. State V (TBC-DEG/TBCC-αβ-tubulin, class 1), and V. An overlay of models, shown in II. vs IV. B) Left, Comparison of αβ-tubulin models in TBC-DEG-αβ-tubulin (light red) with the TBC-DEG/TBCC-αβtubulin (dark red). β-tubulin is used as an alignment reference. Right, soluble αβ-tubulin (cyan) aligned to the two states shown on the left. C) Close-up view of the overlay shown in the left panel of C. showing the N-site and intradimer interfaces with the transitions in α-tubulin marked by the arrow. D) Model of a catalytic cycle for TBC-DEG/TBCC in the biogenesis, degradation, and recycling of αβ-tubulin. Left, TBC-DEG binds αβ-tubulin. I., TBC-DEG deforms α-tubulin by its TBCE-LRR-CapGly. II., the TBCE-LRR-CapGly retracts. III., TBCC-C binds to the Arl2-GTPase, leading to changes in TBCE 3HB/TBCD C-term interface. IV., TBCC-L guides TBCC-N to bind underneath TBCD and engages the αβ-tubulin interface, causing TBCE-LRR-CapGly to rotate. This rotation causes TBCE-CapGly to pull the α-tubulin C-terminal tail inducing a native-like configuration of α-and β-tubulin.

Close-up views of the three TBCC interfaces with TBC-DEG-αβ-tubulin suggest a mechanism to link Arl2 GTP hydrolysis activation to the gating of αβ-tubulin biogenesis (Figure 4). The TBCC β-helix, Arl2, and the TBCD-turret create an interface through conserved ionic and hydrophobic residue interactions (Figure 4A-B, C). The conserved Arg186 and Gln184 residues in TBCC bind the Arl2 GTP nucleotide (Figure 4C; Figure S14A-B, E). In the TBCC-L N-terminal region, the TBCC-L is conserved and is composed of alternating basic residues, Lys 79, Gln78, Lys77, Arg72, Lys 70, Lys68, and aromatic or hydrophobic residues Leu73, Phe74, and Phe76 (Figure 4D; Figure S13A; Figure S14A-B, Figure S16A-B, F-G). These bind TBCD conserved acidic residues Glu206, Asn205, Asp238, Glu235, Asp241, Glu245, Glu283 and hydrophobic residues Phe238, Leu240, Val244 between TBCD H11 and H12 α-helices (Figure 4A-B, D; Figure S11, S13A, S14A-B, S16A-B, F-G). TBCC-N binds tightly to a site underneath TBCD-repeats H11 and H10 through an extensive network of hydrophobic and ionic residues on the TBCD intra-HEAT turns of H11 and H12 with the second and third α-helices of TBCC-N (Figure 4A-B, E). A short end of the TBCC-N α-helical bundle at the ends of its three α-helices, and the turn between the second and third α-helices bind to β- and α-tubulin at the intradimer interface (Figure 4A-C). The TBCC-N bundle primarily contacts α-tubulin forming a wedge against it (Figure 4A-C). TBCC-N contacts acidic α-tubulin H2, H3 residues Glu97, Glu113, and Asp116, and H6 Glu411 (Figure 4C). These residues lie within an 8-Å distance of the α-tubulin N-site GTP γ-phosphate. Alphafold2 models for TBCC show that the TBCC-N first helix varies in length extensively across species. Its binding site and location in these structures suggest that the TBCC extension may facilitate more interactions with TBCD in metazoans to stabilize further the TBCC-N interface with αβ-tubulin (Figure S13B).

### TBCC catalyzes transitions in TBC DEG-αβ-tubulin

Comparisons between the two binary and two ternary assembly states reveal a series of catalytic transitions amongst TBC-DEG subunits and the impact of the binding of TBCC domains on TBC-DEG and αβ-tubulin (Figure 5A, Figure S4, Figure S11). The TBCE LRR-CapGly undergoes the greatest transitions when comparing its conformation in each of the structures. In State I (TBC-DEG-αβ-tubulin class2), the TBCE LRR tightly binds the side of α-tubulin, and the CapGly is positioned to its tip (Figure 5A, panel I). In State II (TBC-DEG-αβ-tubulin class1), TBCE retracts from α-tubulin and aligns parallel to the αβ-tubulin longitudinal axis with its CapGly repositioned along the LRR (Figure 5A, panel II). In State III (TBC-DEG/TBCC-αβ-tubulin class2), TBCC-C binding to the Arl2 GTPase induces changes in the switch I and II elements of Arl2 (Figure S9G), causing TBCD to create a tighter β-tubulin interface in comparison to the TBCD interface in the binary assembly structures (Figure 5A, panel III; Figure 3D). The TBCE-LRR-CapGly arm rotates clockwise and downward in comparison to the binary assembly due to the 3HB region extending while anchored along the TBCD C-terminus (Figure 5A, panel III). Additional conformational changes in the TBCE LRR-CapGly and the TBCD spiral are induced by TBCC-N binding to TBC-DEG-αβ-tubulin. In State IV (TBC-DEG/TBCC-αβ-tubulin class1), the TBCE LRR further rotates clockwise and translates (Figure 5A, panel IV). Additionally, the C-terminal end of the TBCD spiral rotates S° upwards, and the pivot point of this rotation is the TBCC-N docking site on TBCD (Figure S9A, S10A).

Changes in the network of interactions across the TBC-DEG states suggest a pathway through which TBCC binding drives the reorientation of α-tubulin and β-tubulin in the heterodimer. The clockwise movement of the TBCE-LRR-CapGly lever arm is depicted in Figure 5A, panel V. This movement is mediated by the unfurling of the 3HB hinge forming extended α-helices that are ordered due to the TBCD spiral S° rotation (Figure 3C-D; Figure 5A, panel IV; Figure S9B, Figure S10A). The tighter association of TBCD and TBCE-Ubq with the β-tubulin further occludes its longitudinal polymerization interface, potentially explaining the competition between ΔN-DARPIN and TBCC binding in the ternary assembly reconstitution studies. The TBCE LRR rotation impacts the CapGly interaction with α-tubulin and its C-terminus (Figure 3C-D, Figure 5, Figure S10B-C). In State II, the TBCE-LRR-CapGly dissociates from and moves laterally from α-tubulin compared to its bound state in State I (Figure 5A). Comparison of State II to State III shows a 4° rotation in α-tubulin at the intradimer interface, likely caused by the pulling of the α-tubulin C-terminal tail by the TBCE CapGly, returning the αβ-tubulin to a native conformation (Figure 5A-B, panel V). Comparing the conformations of the TBCE LRR-CapGly in all four states shows stepwise transitions of TBCE, starting from a state bound to α-tubulin, proceeding to an unbound state longitudinally aligned to α-tubulin, to successively rotated states upon binding TBCC-C and subsequent TBCC-N binding.

## Discussion

The regulation of α- and β-tubulin mRNA translation, nascent transition to folding, and their subsequent ATP-dependent folding are carried out in parallel by the SCAPERS/ TTCS, Prefoldin, and CCT chaperonins respectively^43-46^.

However, these systems do not recognize differences between α- and β-tubulins. In contrast, the conserved TBCC, TBCD, TBCE, and Arl2 GTPase system evolved to uniquely recognize α- and β-tubulins to form the αβ-tubulin heterodimer^8,9,35^. By regulating αβ-tubulin assembly, TBC activities govern MT polymerization and dynamics^8,9,35^. Here, we describe the reconstitution of the conserved TBC-DEG assemblies and their structural transitions during αβ-tubulin biogenesis regulation.

The conserved topology of αβ-tubulin requires energy for its formation. Our structural studies suggest that Arl2 GTP hydrolysis is the source of this energy and TBCC activates the catalysis of TBC-DEG as a platform for αβ-tubulin assembly and degradation. Eukaryotic cells maintain stoichiometric α- and β-tubulin expression, which is critical for MT function and homeostasis^47^. The over-expression of α-tubulin is less toxic than the overexpression of β-tubulin for eukaryotic cells^48^. β-tubulin overexpression leads to the formation of insoluble aggregates, in contrast, such aggregates do not form when α-tubulin is overexpressed^48^. In both settings, MT functions are highly disrupted^48^. The TBC-DEG binding site to β-tubulin likely evolved to prevent the formation of insoluble polymers of β-tubulin. The extensive coverage of TBC-DEG surrounding the β-tubulin ensures that β-tubulin remains isolated through the activities of the TBC-DEG until αβ-tubulin is assembled.

Our model suggests that TBC-DEG binding to αβ-tubulin deforms the α-tubulin configuration in the tubulin heterodimer, and TBCC binding then drives a series of changes that reform the native αβ-tubulin conformation (Figure 5D, I). The binding of TBCC-C to Arl2, then TBCC-L to TBCD guides TBCC-N to bind underneath TBCD and recognize the αβ-tubulin intradimer interface (Figure 5D, IV). TBCC-N binding induces the rotation of TBCE mediated by the TBCD C-term/TBCE-Ubq region’s upward movement and the 3HB downward rotation (Figure 5D, II-IV). This causes the TBCE LRR-CagGly to undergo a clockwise translation and pull α-tubulin via its CapGly interaction with the α-tubulin C-terminal tail (Figure 5D, IV). The TBCC-activated Arl2 GTP hydrolysis is a critical energy source for each αβ-tubulin assembly event. The role of Arl2 GTP hydrolysis is also consistent with the well-documented, severe defects of Arl2 GTP- and GDP-locked mutants on MT dynamics in interphase and mitosis^18^. The GTP-dependent mechanism of TBC-DEG/TBCC is likely essential to overcome the stable α- and β-tubulin interface. The model is also consistent with the idea that the TBC-DEG/TBCC cycle may reset soluble αβ-tubulin configuration to recycle αβ-tubulin for MT polymerization.

The Cryo-EM structures reveal that TBC-DEG forms extensive binding sites along αβ-tubulin (Figure 1, Figure 2C-D, H-K). Our structures explain the lack of genetic complementation of large tags fused at the α- and β-tubulin C-terminal tails and β-tubulin N-terminus ^39,40^. Across many eukaryotes, the α-tubulin N-terminus is the only site for placing fluorescent protein fusions to complement native tubulin genes^39^. This is consistent with the exposure of this region of α-tubulin in our structures. Although large tags are not tolerated, short eight to ten residue tags (such as HA, His, strpII) are tolerated at the tubulin N or C-termini, consistent with their binding sites on TBC-DEG having sufficient space available for them^39^.

## Conclusions

The reconstitution of yeast TBCD, TBCE, and Arl2 to form TBC-DEG has led to isolating the conserved TBC-DEG assembly which has remained elusive until recently. Our multiple cryo-EM structures for TBC-DEG-αβ-tubulin and TBC-DEG/TBCC-αβ-tubulin reveal a mechanism for TBC-DEG in regulating the α- and β-tubulin configuration in αβ-tubulin biogenesis, and the critical roles of TBCC and Arl2 GTP hydrolysis in forming the αβ-tubulin configuration.

## Acknowledgments

We thank the Bay Area Cryo-EM consortium led by Prof. Eva Nogales (Molecular Cell Biology, UC-Berkeley) and the UC-Davis BioEM facility for large-scale Cryo-EM data collection support. Large Scale Cryo-EM data were collected at UC-Davis and the Cryo-EM facility at UC-San Francisco with support From Dr Alexander Mysanikov (Biochemistry and Biophysics, UC-San Francisco). We thank Dr. Camille Scott and the UC-Davis High Performance Computing (HPC) facility for computational HPC infrastructure building and support. We thank Dr Stanley Nithianantham (Molecular Cellular Biology, UC-Davis) for initial studies and preliminary biochemical work. We thank Prof Jeffrey K Moore (Cell Developmental Biology, University of Colorado, Anschutz Medical Campus) for advice and suggestions and the critical reading of this manuscript. We thank Prof Richard McKenney, Prof Jonathan Scholey (Molecular Cellular Biology, UC-Davis), and Prof Ahmet Yildiz (Molecular Cell Biology, UC-Berkeley) for comments on the manuscript. JAB and JKM are supported by funding from the National Institutes of Health (NIH) GM110283.

## Author contribution

AT purified and assembled TBC-DEG/TBCC-αβ-tubulin complexes, collected cryo-EM data, determined and refined all single particle cryo-EM structures, built, refined and validated all models for assemblies, prepared figured, wrote and edited the manuscript. ZW purified and assembled TBC-DEG-αβ-tubulin complexes, collected cryo-EM data, determined initial single particle cryo-EM structures, and built initial models. BS built, refined and validated models. FG supported cryo-EM grid preparation, cryo-EM screening, and large-scale cryo-EM data collection. JAB planned and managed the project, trained scientists, obtained funding for the project, prepared assemblies, prepared figures, and wrote and edited the manuscript.

## Materials and Methods

### Protein Expression and Purification

Recombinant TBC-DEG was purified as previously described^33^: Polycistronic expression constructs were co-transformed into a SoluBL21 bacterial strain (AMSBIO) with one construct containing yeast TBCD (Cin1p) with an N-term 6Xhis tag, and the other containing TBCE (Pac2p) and either wild type or GTP-locked (Q73L) Arl2 (Cin4p). SoluBL21 bacterial expression strain were selected for by both Ampicillin and Kanamycin resistance. Transformed SoluBL21 cells were grown large scale at 37°C with antibiotic resistance and induced using 0.5 mM isothio-beta-glucopyranoside (IPTG) once an optical density (600 nm) of 0.6 Absorbance was achieved, and growth continued at 19°C overnight. Cells were pelleted and resuspended with lysis buffer (50 mM HEPES, 150 mM KCl, 1 mM MgCl2, 3 mM - mercaptoethanol, with protease inhibitors, including 1 mM PMSF, 1 µg/ml leupeptin, 20 µg/ml benzamidine, and 40 µg/ml aprotinin (Sigma Aldrich)) then lysed using a microfluidizer. Cell lysates were clarified by centrifugation at 18,000 rpm for 30 min at 4°C. Ni-IDA affinity beads (Macherey-Nagel, Bethlehem, PA, USA) were used to purify TBC-DEG assemblies. Protein was eluted through 5 column volumes of lysis buffer containing 200 mM imidazole. Eluted protein was then diluted to a final salt concentration of low salt buffer conditions (100 mM KCl, 50 mM HEPES, 1 mM MgCl2) and bound to HiTrap SP FF (GE Healthcare, USA) anion exchange column. Elution was done using 5 column volumes through a gradient with high salt buffer (500 mM KCl, 50 mM HEPES, 1 mM MgCl2). Ion exchange TBC-DEG purified fractions were then immediately concentrated using Amicon concentrators (Fisher Scientific) and then loaded on a HiLoad 16/600 Superdex-200 gel filtration column (GE Healthcare, USA). TBC-DEG was then concentrated, aliquoted, and frozen in liquid nitrogen. Recombinant yeast untagged TBCC was expressed in SoluBL21 and purified using the approach described previously^33^. Soluble αβ-tubulin was purified from porcine brains using the previously well-described method^49^. The codon-optimized capping proteins, ΔN-DARPin and α-Rep iH5, were synthesized using published sequences^41,50^. The cDNAs were assembled into pET T7 bacterial expression vectors in frame with the affinity tags 6Xhis tag or StrepII tags respectively, and then were expressed in SoluBL21 cells grown at 37°C. Expression was induced with 0.5 mM IPTG upon reaching an optical density (600 nm) of 0.6 Absorbance and cells were grown at 19 °C overnight. Cells were pelleted and then lysed using microfluidizers and a protease inhibitor cocktail was added to the lysis buffer (50 mM HEPES 300 mM KCl pH 7.0, 3 mM β-Mercaptoethanol). ΔN-DARPin and iH5 were purified using Ni-IDA or Streptactin XT columns, respectively, and eluted with lysis buffer containing the addition of either 100 mM imidazole or 50 mM Biotin. The iH5 and ΔN-DARPin were then further purified by binding to a HiTrapQ ion exchange column under low salt buffer conditions (50 mM HEPES, 100 mM KCl, 3 mM β-Mercaptoethanol) and then eluted in five-column volume elution through a gradient with high salt buffer (50 mM HEPES, 1M KCl, 3 mM β-Mercaptoethanol).

### Assembly of TBC-DEG-αβ-tubulin and TBC-DEG/TBCC-αβ-tubulin

TBC-DEG-αβ-tubulin assemblies were reconstituted as described previously^33^, and as follows: Recombinant TBC-DEG (5 µmol) was incubated with either 5 mM GTP or GTPyS and then mixed with equimolar porcine brain a-tubulin and ΔN-DARPin. TBC-DEG-αβ-tubulin assemblies were purified by size exclusion chromatography (SEC) using a Superdex 200 10/300 column (GE Healthcare, USA) in binding buffer (50 mM HEPES, 130 mM KCl, and 3 mM -mercaptoethanol at pH 7.0). Samples were collected and analyzed using SDS-PAGE (Bio-Rad, Hercules, CA, USA).

TBC-DEG/TBCC-αβ-tubulin assemblies were reconstituted as described previously^33^ and as follows: recombinant TBC-DEG with Q73L Arl2 (5 µmol) was mixed with equimolar amounts of porcine αβ-tubulin, TBCC, iH5, and 5 mM GTPγS. TBC-DEG-αβ-tubulin assemblies were purified by size exclusion chromatography (SEC) using a Superdex 200 10/300 column (GE healthcare) in binding buffer (50 mM HEPES, 130 mM KCl, and 3 mM -mercaptoethanol at pH 7.0). Samples were collected and analyzed using SDS-PAGE (Bio-Rad, Hercules, CA, USA).

### Cryo-EM sample preparation and imaging

SEC purified complexes were either obtained at 0.1 mg/ml TBC-DEG-αβ-tubulin or TBC-DEG/TBCC-αβ-tubulin, or they were diluted to that concentration. Complexes were crosslinked by incubation with 20 nM BS3 crosslinker for 1-2 hours and then treated with 100 nM Tris-HCl at pH 7 to neutralize the reaction. Copper Quantifoil R1.2/1.3 grids (Thermofisher) coated with 2nm carbon layer grid support were glow discharged, and 4µl of sample was applied to the grid surface. The sample was incubated in the humidity chamber of a Mark III vitrobot (Thermo Fisher) at 20°C and 100% humidity for 30 seconds. Grids were blotted at blot force 8 for 3-5 seconds and then plunged into liquid ethane. TBC-DEG-αβ-tubulin grids were screened, and two 5-6K movie datasets were collected using a Thermofisher Titan Krios operated using a Gatan K2 direct electron detector collecting 80 frames per 2 sec at 80 electron dose. TBC-DEG/TBCC-αβ-tubulin grids were imaged using Thermofisher Glacios operated using a Gatan K3 direct electron detector collecting 80 frames per 2 S at 80 electron dose.

### Single particle analysis pipeline

For each dataset, the movies were motion corrected through RELION^51^ suite Motioncor2 using a 5X7 patch and B-factor of 150. Images were picked by LoG Picker and were subjected to 2D classification using either RELION 3.2-4.1^5,1,52^ or Cryosparc 4.1^53^. Multiple rounds of 2D-Classification were used to remove junk particles.

For TBC-DEG-αβ-tubulin a *de novo* starting model was generated by the best 2D projections in RELION and was followed by three rounds of 3D classification to sort particles and remove junk or broken particles (Figure S2). The particles in the best classes were then combined for a 3D auto-refinement in RELION, z-flipped, and then subjected to CTF refinement and Bayesian polishing. A subsequent 3D auto-refinement led to a 3.6A structure containing a noisy region which was determined to be continuous heterogeneity. These particle images were then transferred to Cryosparc, and 3D Variability Analysis^37^ was performed on the particle pool. Three modes of variability were selected, intermediate mode was used containing 10 frames, and resolution was filtered to 6A. 3DVA separated frames of particle images in two pools with different conformations of the TBCE arm region (Figure S3). Frames were pooled together based on their position, subdomain organization, and general interaction interface with α-tubulin. Two pools of particles were identified with two distinct conformations. A 3D-auto refinement in Cryosparc was performed on each pool using the most representative frame of the pool for each refinement. DeepEMhancer^54^ was used to sharpen the resulting reconstructions using either the default sharpening model or the “wide target” model. The final processing parameters are described in Table I.

For TBC-DEG/TBCC-αβ-tubulin, three *de novo* starting models were generated in Cryosparc through *Ab-initio* reconstruction by particle stacks with clear secondary structure displayed in the 2D-class averages. All starting models were subjected to heterogeneous refinement in Cryosparc^53^. A single class containing the best pool of particles was subjected to a homogeneous refinement in Cryosparc without a mask. These data were then converted into RELION format where the following steps were performed, 3D-auto refinement, Bayesian polishing, and CTF refinement leading to a 3.6A structure. Continuous heterogeneity was displayed in a periphery region of the structure as well as part of the core. These regions were masked, and after conversion to Cryosparc file format, were subjected to 3D Variability Analysis^37^. Two separate 3DVA steps were performed using a different mask for each step. In the first 3DVA, a wide mask was used including the periphery of the structure displaying continuous heterogeneity. Three components were determined, intermediate mode was used consisting of 10 frames, and resolution was filtered to 6A. Component #1 focused on refining the TBCC-bound particles revealing two groups of particles in frames 0-3 and 7-10, which were subjected to local refinements in Cryosparc (Figure S6, Figure S7A, middle right). Component #2 focused on refining the TBC-DEG-αβ-tubulin core conformation revealing two groups of particles in frames 0-3 and 7-10, which were subjected to local refinements in Cryosparc leading to two unique states (Figure S6, bottom right; Figure S7B). Component #3 focused on refining the relationship between TBCC binding and TBCE arm conformation revealing two groups of particles in frames 0-3 and 7-10, which were subjected to local refinements in Cryosparc leading to two unique states (Figure S6, middle left; Figure S7C). Each of the three components was analyzed and comparisons were made between them to understand subunit relationships with each other. A second 3DVA step was performed with a tight mask around the core based on the outcome of one of the components of the previous 3DVA step. This 3DVA step focused on understanding the heterogeneity of the periphery region of the structure. Three components were determined, intermediate mode was used consisting of 10 frames, and resolution was filtered to 5.5A. Two pools of particles were identified and separated as unique TBCE states. The frames were pooled based on similar positioning and subjected to local refinements in Cryosparc to achieve final reconstructions. All structures were sharpened using DeepEMhancer with a “wide target” model. The final processing parameters are described in Table I.

### Model building and refinement

The two TBC-DEG-αβ-tubulin binary composite Cryo-EM maps were built using a combination of Coot^55^ and PHENIX^56^ starting with the AlphaFold2 models for yeast TBCD, TBCE, and Arl2 (PDB IDs: AF-P40987-F1, AF-P39110-F1, AF-P39937-F1) and the porcine αβ-tubulin (PDB ID:1FFX). Alphafold2 models were initially fit using FlexEM from the CCPEM program suite^57^. The TBC-DEG-αβ-tubulin was of sufficient resolution to build the full polypeptide chain of TBCD, Arl2, and TBCE Ubq domains and resolve the αβ-tubulin interactions. Three Nucleotides (Arl2-GTP, α-tubulin-N-site GTP, β-tubulin E-site GDP) were also built into the maps (Figure 1F, right panel; Figure S4). The two binary state maps for TBCE (Class1 and Class2) map densities were built based on the placement and minor changes to the LRR and CapGly, but manual placement of the 3HB helices and connection between LRR and CapGly was necessary. The models were subjected to cycles of Coot-based manual building of loops and side chain corrections and real space refinements in PHENIX. The final model validation was performed in PHENIX (table). Models and maps are deposited into the R5CB and EMDB (EMDB ID: XXXX, PDB ID: XXXX), and described in table I. Figures were generated using ChimeraX^58^.

Two Composite TBC-DEG/TBCC-αβ-tubulin ternary maps were generated by stitching mask-based regions of maps generated from 3DVA components as described in Figure S8. The models for TBC-DEG-αβ-tubulin were utilized from the binary structures, while TBCC domain models were models based on the crystal structure of TBCC-C (PDB ID: 5CVA)^33^ and AlphaFold2 model for full TBCC sequence (PDB ID: AF-P46670-F1). Models for the two states of the TBC-DEG-αβ-tubulin cores were built by placing and modifying the models for subunits based on density maps using Coot. The two ternary state maps for TBCE (Class1 and Class2) map densities were built based on the placement and minor changes to the LRR and CapGly TBCE models from AlphaFold2. Manual placement of the 3HB helices and connection between LRR and CapGly was necessary due to lower resolution. The models were subjected to cycles of PHENIX real-space refinement and Coot-based manual building of loops and side chain corrections. Models and maps are deposited into the RSCB and EMDB (EMDB ID: XXXX, PDB ID: XXXX) and described in Table I. Figures were generated using ChimeraX^58^.

## Extended Data Figures

**Figure S1:**
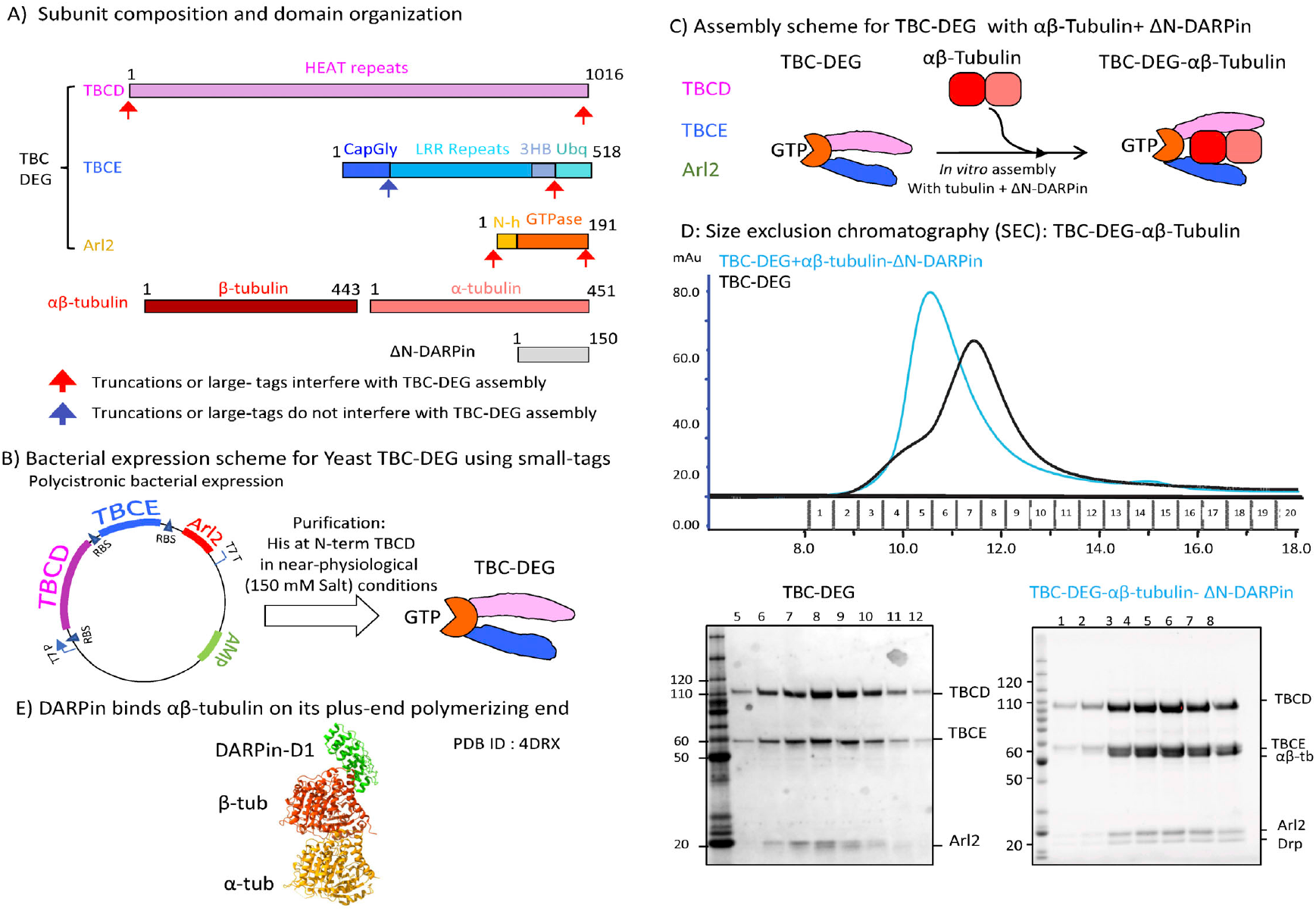
Purification of recombinant TBC-DEG and Reconstitution of the TBC-DEG-αβ-tubulin assemblies for cryo-EM. A) Subunit and domain organization, and residue length of TBCD, TBCE, Arl2, α-tubulin, β-tubulin, and ΔN-DARPin. Red arrows denote sites where subunit truncations or insertions of large protein tags led to defects in TBC-DEG solubility, Blue arrows denote sites where subunit truncations or insertion of large protein tags did not lead to defects in TBC-DEG solubility as described^33^. B) Expression scheme for TBCD, TBCE, and Arl2 purified using a short 6X his tag on the N-terminus of TBCD and purified from *E. coli* using physiological ionic conditions C) Scheme describing the reconstitution of TBC-DEG with soluble αβ-tubulin to form TBC-DEG-αβ-tubulin assemblies. Note that ΔN-DARPin was utilized to stabilize the αβ-tubulin and prevent TBC-DEG-αβ-tubulin aggregation. D) Top panel, Size exclusion chromatography (SEC) for TBC-DEG alone (black trace) and TBC-DEG-αβ-tubulin-ΔN-DARPin (blue trace) showing the elution profiles of the assemblies. Bottom panel, left, SDS PAGE showing contents of SEC fractions for TBC-DEG alone including TBCD, TBCE, and Arl2 subunits. Right, SDS PAGE showing fractions of TBC-DEG-αβ-tubulin for SEC fraction of TBC-DEG/αβ-tubulin showing the contents of TBCD, TBCE, Arl2, α-tubulin, β-tubulin, and ΔN-DARPin. F) The binding interface for ΔN-DARPin to the plus-end polymerizing interface of β-tubulin as observed in previous structures.

**Figure S2:**
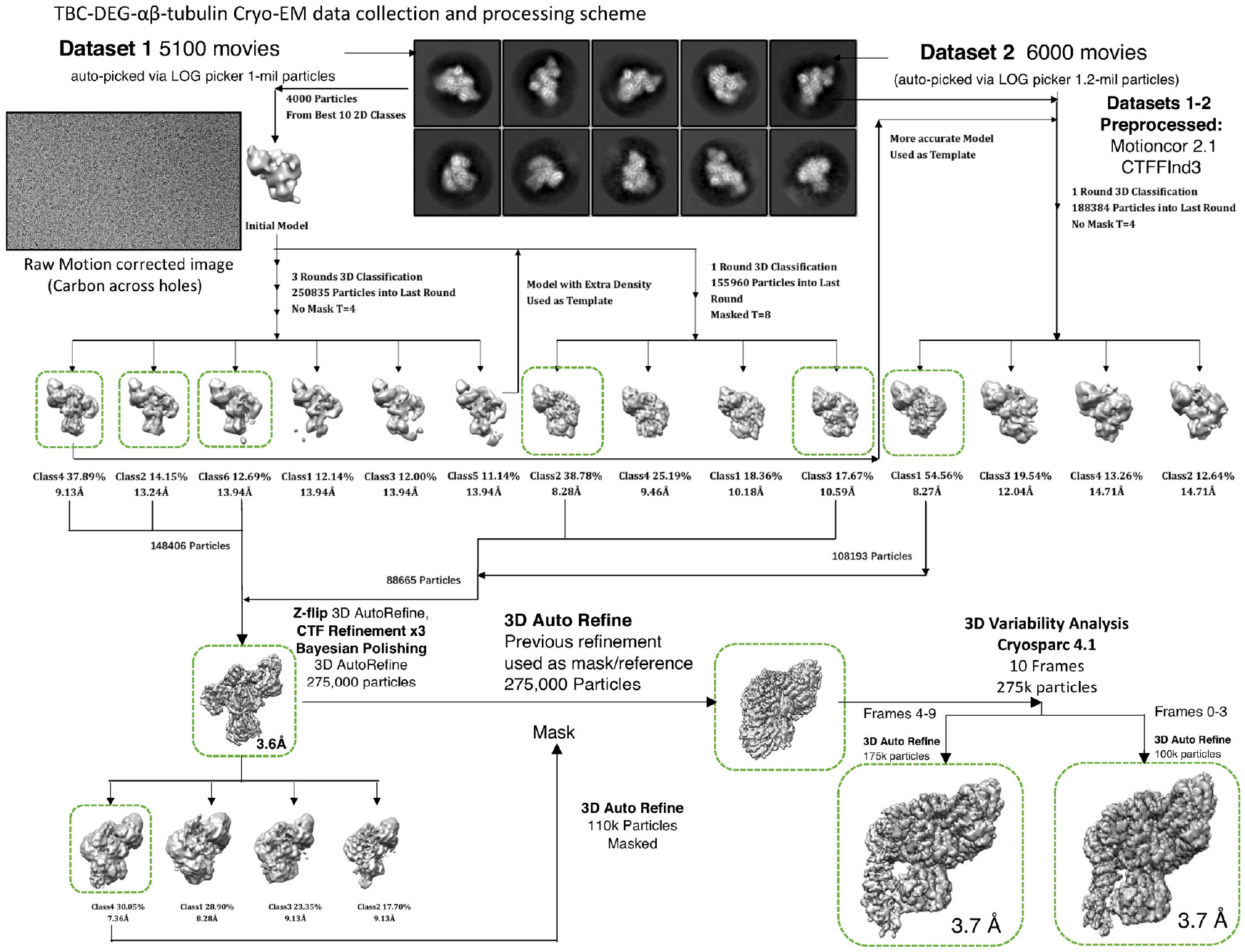
Cryo-EM image data collection and single particle image process for TBC-DEG-αβ-tubulin complexes leading to structures of two unique states (class 1 and class 2). Top to bottom, two TBC-0EG-αβ-tubulin datasets (top left and top right; example image shown in the middle) were collected and raw images were pre-processed using motioncorr2, CTFfind3 then used identify coordinates for particle images, which were processed using a combination of Relion 3, Relion 4 and cryosparc. Multiple cycles of 30 classification and 30 refinement led to 3.6-A TBC-0EG-αβ-tubulin core particle with low resolution for a mobile arm-like extension. The conformation of the arm-like extension was resolved using 30VA analysis using a mask around the arm region leading two unique classes (class1 and Class2) in which TBCE conformation was unique. Final processing statistics are described in Table 1.

**Figure S3:**
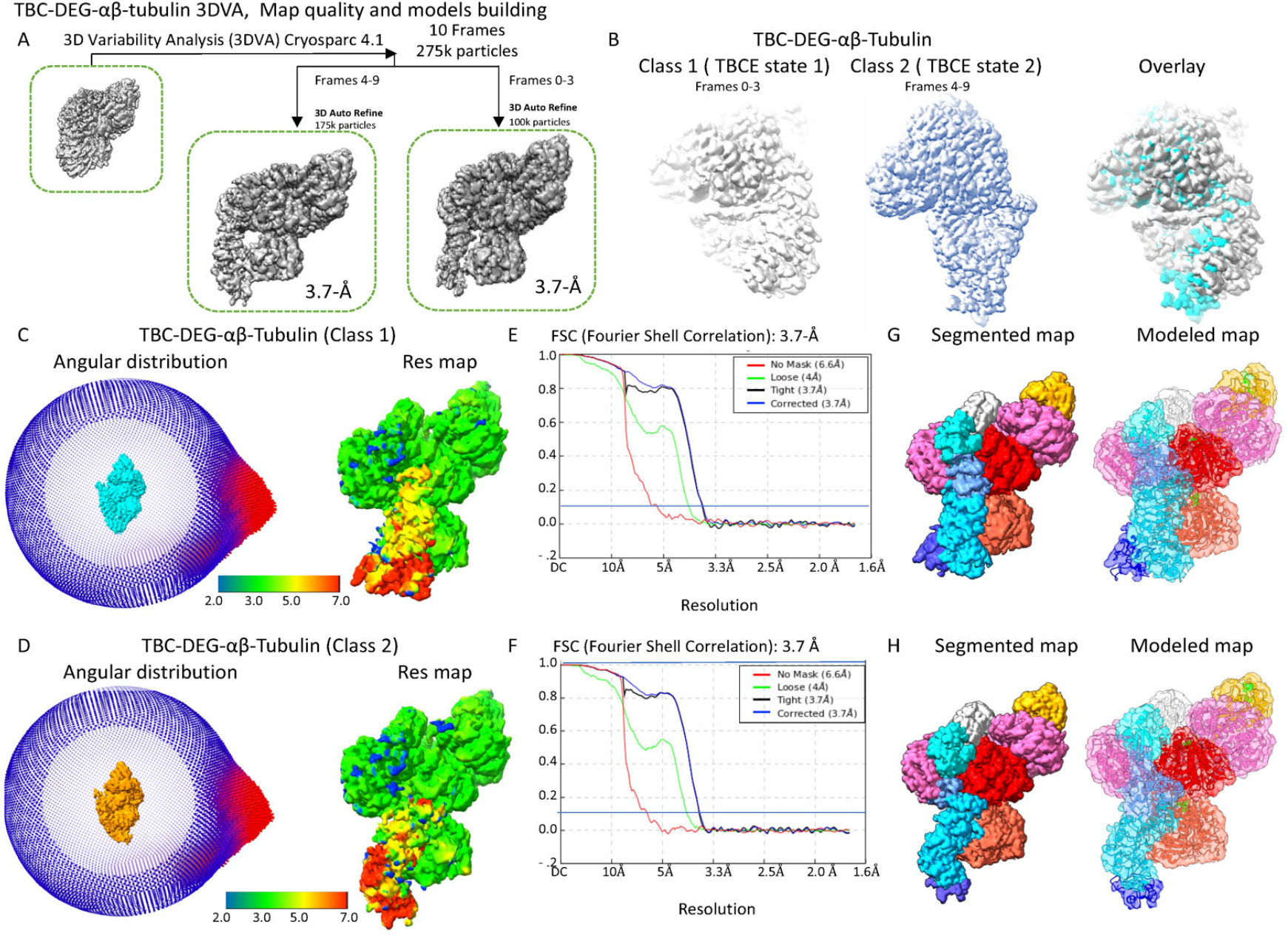
Details of the 3D variability (3DVA) analysis of TBC-DEG-αβ-tubulin cryo-EM maps leading to the two TBCE conformations bound to α-tubulin. A) Close-up view of the frame-based 3DVA analysis leading two classes which differ in the arm-like extension. 175K particle images led subsets which were refined leading to Class 1 and represented frames 0-3. 175K particle images led subsets which were refined leading to Class 1 and represented frames 7-10. B) Comparison of TBC-DEG-αβ-tubulin Class 1 and Class 2 refined cryo-EM maps. Left panel, refined Class1 TBC-DEG-αβ-tubulin cryo-EM map (frames 0-3). Middle panel, refined Class2 TBC_DEG-αβ-tubulin cryo-EM map (frames 7-10). Right panel, overlay of both maps showing the conformation change in the Arm-like extension representing TBCE. C) Left, Angular distribution for TBC-DEG-αβ-tubulin class1 particles in the final refined map. Middle panel, Resmap for TBC-DEG-αβ-tubulin in class 1 showing the resolution distribution color range onto the structure. D) Left, Angular distribution for TBC-DEG-αβ-tubulin class 2 particles in the final refined map. Middle panel, Resmap for TBC-DEG-αβ-tubulin in class 2 showing the resolution distribution color range onto the structure. E) Gold standard Fourier Shell correlation (FSC) for TBC-DEG-αβ-tubulin class 1. F) Gold standard Fourier Shell correlation (FSC) for TBC-DEG-αβ-tubulin class 2. G) Left, Model-based segmented TBC-DEG-αβ-tubulin-ΔN-DARPin class1 map (segmented map), right, atomic models for subunits placed in segments of these subunits of TBC-DEG-αβ-tubulin-ΔN-DARPin. H) Left, Model-based segmented TBC-DEG-αβ-tubulin-ΔN-DARPin class2 map (segmented map), right, atomic models for subunits placed in segments of these subunits of TBC-DEG-αβ-tubulin-ΔN-DARPin (modeled map).

**Figure S4:**
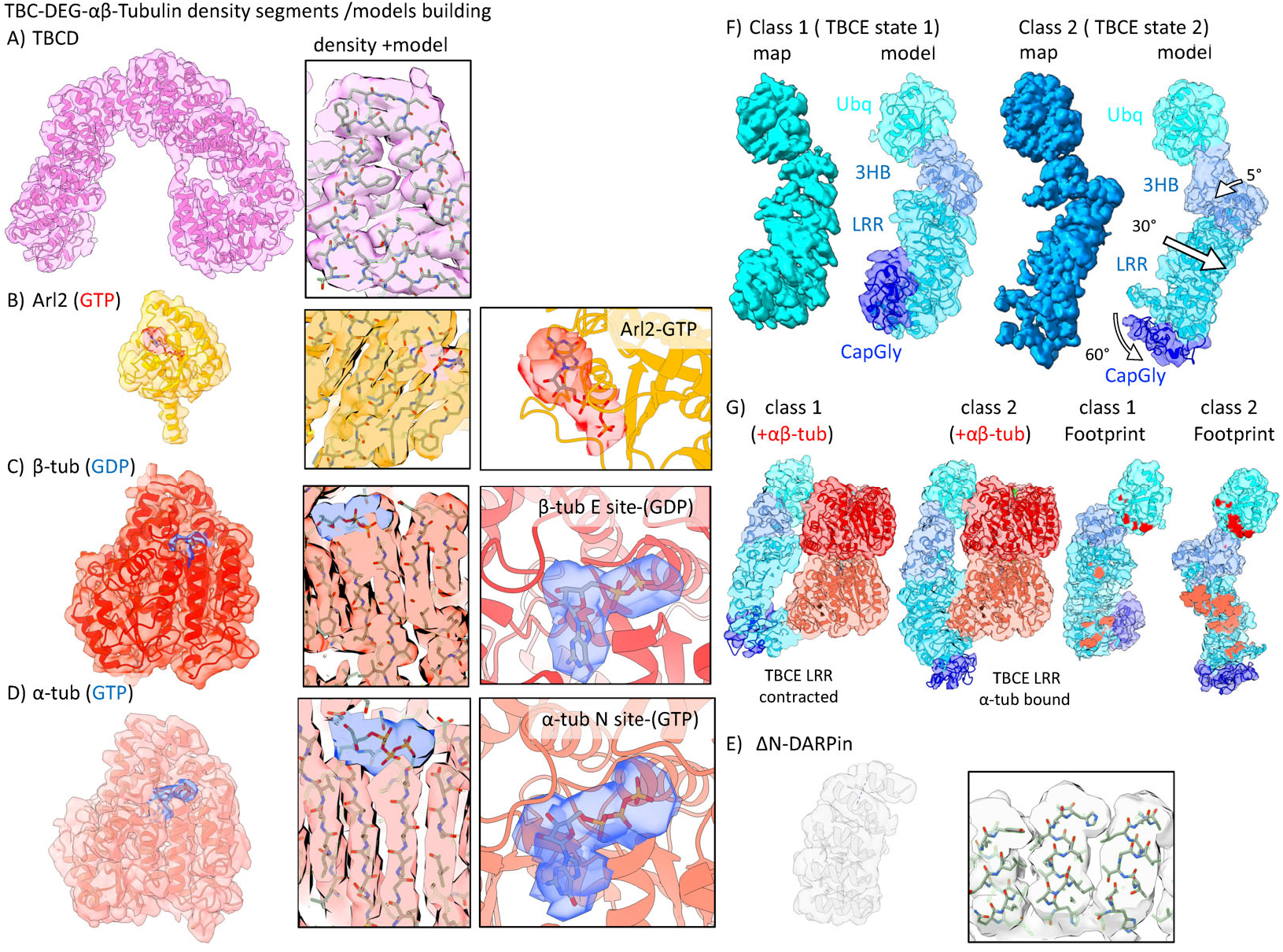
Cryo-EM map segments and atomic models built of each of the TBC-DEG-αβ-tubulin subunits including two unique TBCE models and details of the unique TBCE-αβ-tubulin interfaces in the two states. A) Left panel, Map segment, and Model built for TBCD. right panel, for example, density for an atomic model built into the TBCD electron density. B) Left panel, Map segment, and Model built for Arl2. middle panel, example density for an atomic model built into the Arl2 electron density with GTP shown in red. Right panel close-up view for GTP in Arl2 with electron density shown in Red. C) Left panel, Map segment, and Model built for β-tubulin. middle panel, example density for an atomic model built into the β-tubulin electron density with E-site GDP shown in blue. Right panel close-up view for E-site GDP in β-tubulin with electron density shown in blue. D) Left panel, Map segment, and Model built for α-tubulin. middle panel, example density for an atomic model built into the A-tubulin electron density with the N-site GTP shown in blue. Right panel close-up view for N-site GTP in α--tubulin with electron density shown in blue. E) Left panel, Map segment, and Model built for ΔN-DARPin. right panel, for example, density for an atomic model built into the ΔN-DARPin electron density. F) Left panels, Class 1 TBCE state 1 showing segmented density map (left, sky blue), model fitted in subregion segmented map (right). Right panels, Class 1 TBCE state 2 showing segmented density map (left, cyan), model fitted into subregion segmented map (right). G) Left panels, class1 vs class1 map comparison showing a slice view of TBCE-αβ-tubulin interface showing the class1 TBCE retracted state (class1) versus the TBCE-α-tubulin bound state (class 2). Right panels, class 1 (left) vs class 2 (right) density map viewed with the αβ-tubulin interaction interfaces marked by the color of α-tubulin (light red) and β-tubulin (dark red) showing changes in the TBCE LRR in interfacing with α-tubulin in Class2 compared to class 1

**Figure S5:**
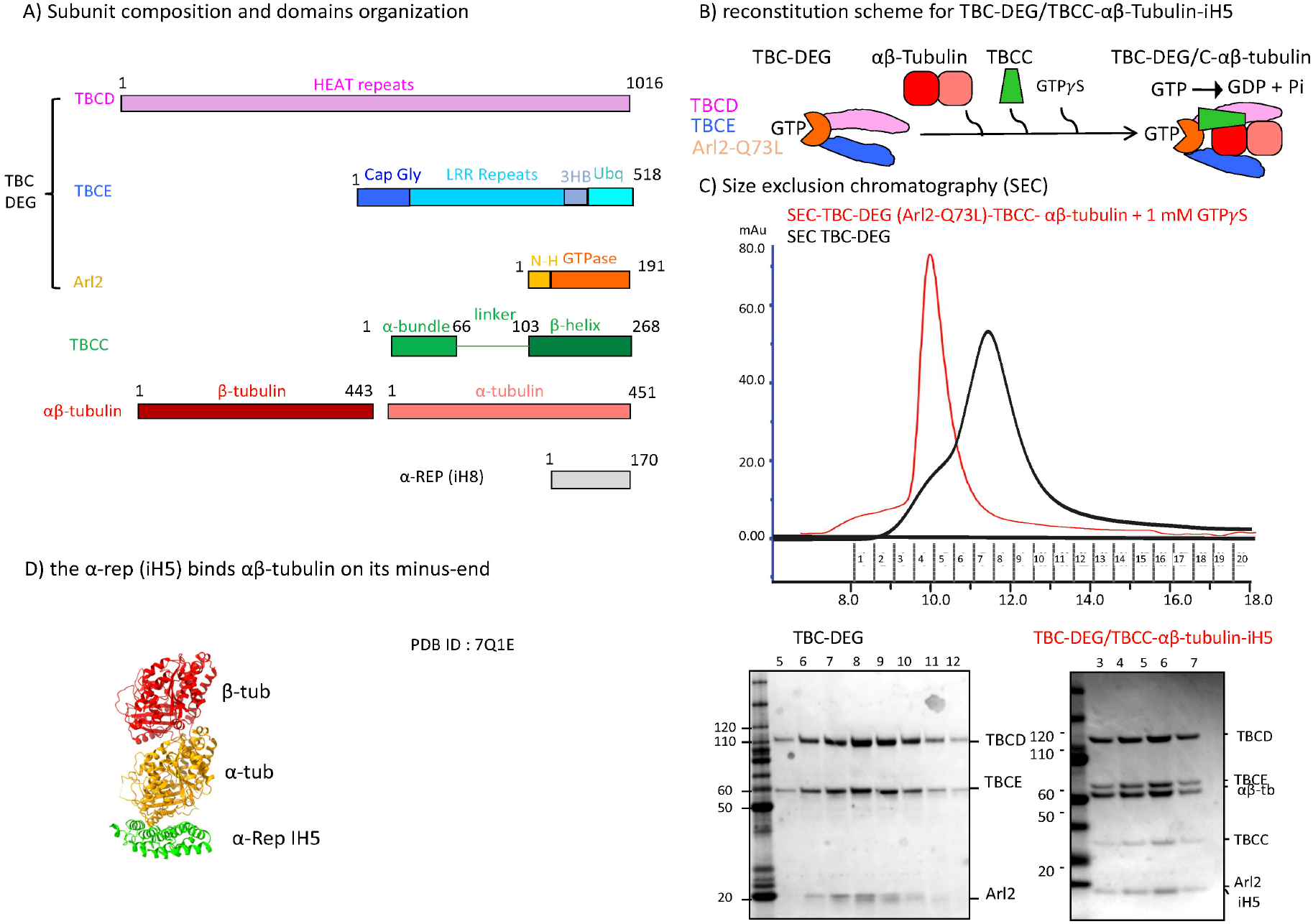
Biochemical reconstitution of TBC-DEG/TBCC-αβ-tubulin assemblies using Arl2 GTP-locked mutant and GTPγ5. A) Subunit and domain organization, and residue length of TBCD, TBCE, Arl2, TBCC, α-tubulin, β-tubulin, and α-rep (iHS). B) Scheme describing the reconstitution of TBC-DEG-Arl2 Q73L (GTP-locked mutant) with soluble αβ-tubulin and TBCC in the presence of GTPγS to form TBC-DEG/TBCC-αβ-tubulin assemblies. Note that α-rep iHS -DARPin was utilized to stabilize the αβ-tubulin in the assembly and prevent aggregation. C) Top panel, Size exclusion chromatography (SEC) for TBC-DEG alone (black trace) and TBC-DEG-Arl2 Q73L-TBCC-αβ-tubulin-GTPγS-ihS (red trace) showing the elution profiles of the assemblies. Bottom panel: left, SDS-PAGE showing SEC fraction contents for TBC-DEG alone including TBCD, TBCE, and Arl2 subunits. Right, SDS-PAGE showing fractions of TBC-DEG-αβ-tubulin for SEC fraction of TBC-DEG/αβ-tubulin showing the contents of TBCD, TBCE, Arl2, TBCC, α-tubulin, β-tubulin and iHS subunits. D) The binding interface for iHS to the minus-end polymerizing interface of α-tubulin as observed in previously determined structures. Note this is the opposite longitudinal surface of αβ-tubulin from ΔN-DARPin which did not bind the TBC-DEG/TBCC-αβ-tubulin assembly.

**Figure S6:**
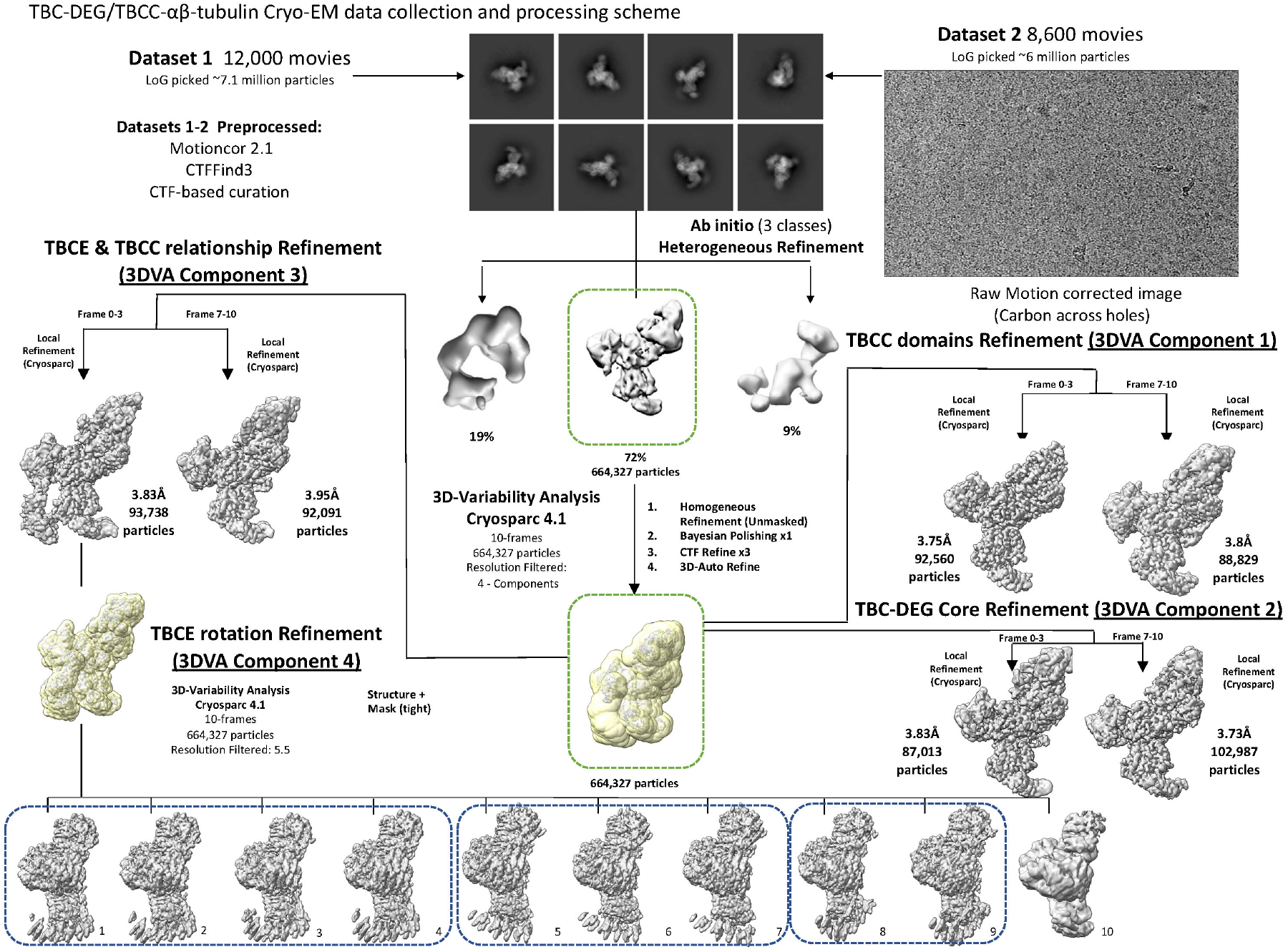
Cryo-EM image data collection and image process for TBC-DEG-αβ-tubulin complexes leading to structures leading to multiple structures of TBC-DEG/TBCC-αβ-tubulin. Top to bottom, two TBC-DEG/TBCC-αβ-tubulin-ih5 datasets (top left, top right); example image shown in the top right) were collected and raw images were pre-processed using motioncorr2, CTFfind3 then used identify coordinates for particle images were processed using a combination of Relion 3, Relion 4 and cryosparc 4.1 leading to 2D-class averages which are distinct from TBC-DEG-αβ-tubulin shown in figure S2. A cycle of ab initio model generation followed by heterogeneous refinement was carried out using a TBC-DEG-αβ-tubulin core map, leading to 72% of particles remaining in a single moderate resolution class. 3D-refinement led to moderate resolution TBC-DEG/TBCC-αβ-tubulin with multiple blurred low-resolution regions. Four components of 3DVA, using cryosparc 4.1, were carried out on the consensus refined TBC-DEG/TBCC-αβ-tubulin particle. TBCC domains were resolved using 3DVA Component 1 (middle right) leading to two classes in which TBCC domains are present or absent. The conformational changes in the TBC-DEG/TBCC-αβ-tubulin core assembly were resolved using 3DVA Component 2 (the lower right) leading two classes in which TBCC-induced conformational changes are present or absent in the TBCD and TBCE Ubq domain. The relationship between TBCE and TBCC-N density binding to the TBC-DEG/TBCC-αβ-tubulin core was resolved using 3DVA Component 3 (middle left). The rotational transition of TBCE arm-like extension was resolved using 3DVA component 4 leading to 10 frames with three distinct states for the TBCE arm

**Figure 7:**
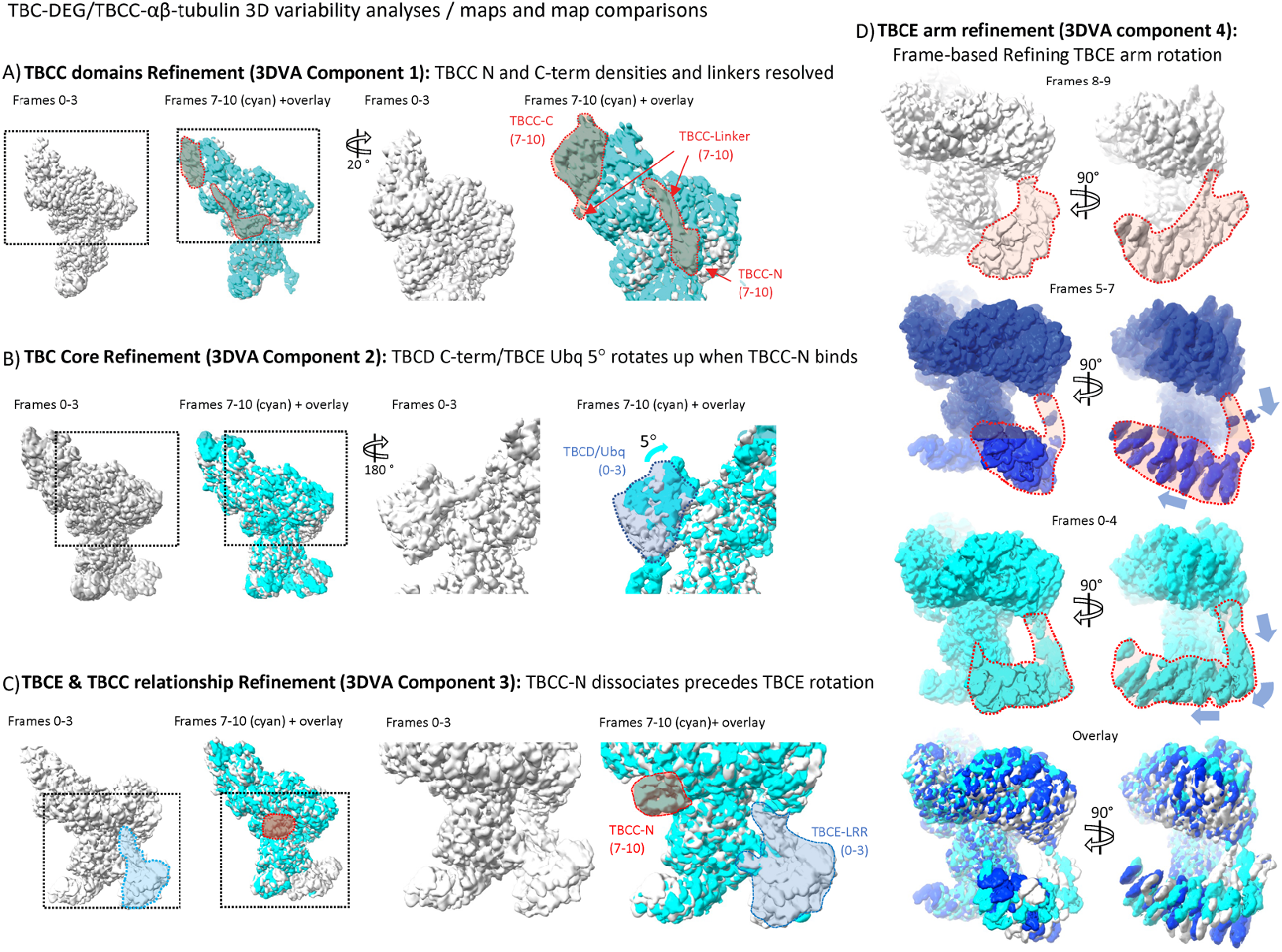
Close-up Comparison of multiple states resolved using four component 3DVA. Each 3DVA component led to two distinct states of TBC-DEG/TBCC-αβ-tubulin assemblies and allowed us to generate states with refined maps for different TBC-DEG/TBCC-αβ-tubulin subregions. A) Close-up comparison of two states resolved in 3DVA component 1: left panels, an overview of frames 0-3 without TBCC domain (light gray, left), and frames 7-10 (cyan)+ frames 0-3 overlay with TBCC domains (highlighted by dotted lines). Right panel, close-up 20° rotated views of right panels, left, showing frames 0-3 (light gray), and frames 7-10 (cyan) overlay on frames 0-3 (light gray) with TBCC-C, TBCC-L, and TBCC-N densities in frames 7-10 (highlighted with dotted red lines) B) Close-up comparison of two states resolved in 3DVA component 2 leading to a 5° rotation in the TBCD spiral/TBCE Ubq. Left panels, frames 0-3 with TBCD Spiral /TBCE Ubq in downward rotation (light gray, left) and frames 7-10 (cyan) with TBCD Spiral/TBCE Ubq in the upward a 5° overlay with frame 0-3 (light gray). Right panels, close-up view of a 180°-rotated view of frames 0-3 (light gray, left), frames 7-10 (cyan) + overlay in frames 0-3 (light gray) showing the 5° rotation upward rotation in frames 7-10 map. C) A close-up comparison of the two states revolved in 3DVA component 3 leading to a relationship between TBCC-N binding and TBCE arm conformation. Left panels, frames 0-3 (light gray, left) with TBCE being ordered (highlighted in blue) if TBCC-N density is missing and frames 7-10 (cyan) with TBCC-N density present (red) with an overlay with frame 0-3 (light gray). Right panels, close up untransformed view showing frames 0-3 (light gray, left) with TBCE density (highlighted in blue) and frames 7-10 (cyan) with TBCC-N density (highlighted in red) + overlay in frames 0-3 (light gray). These maps describe the relationship of the TBCC-N binding to the TBCE arm-like extension. D) Close-up comparison of three states resolved in 3DVA component 4 leading the three states of the TBCE arm-like extension. Top panel, two views of frames 8-9 (light gray) showing the two views of the TBCE arm. Middle panels, identical orientations of frame 5-7 (dark blue) showing two states of the TBCE arm with changes in the junction and rotation. The third set of panels frames 0-4 (cyan) of TBCE Arm-like extension showing the conformational transition and rotation of the TBCE arm. The bottom panels, show an overlay of all three states demonstrating the the tilt, and rotation of the TBCE arm around the tubulin.

**Figure S8:**
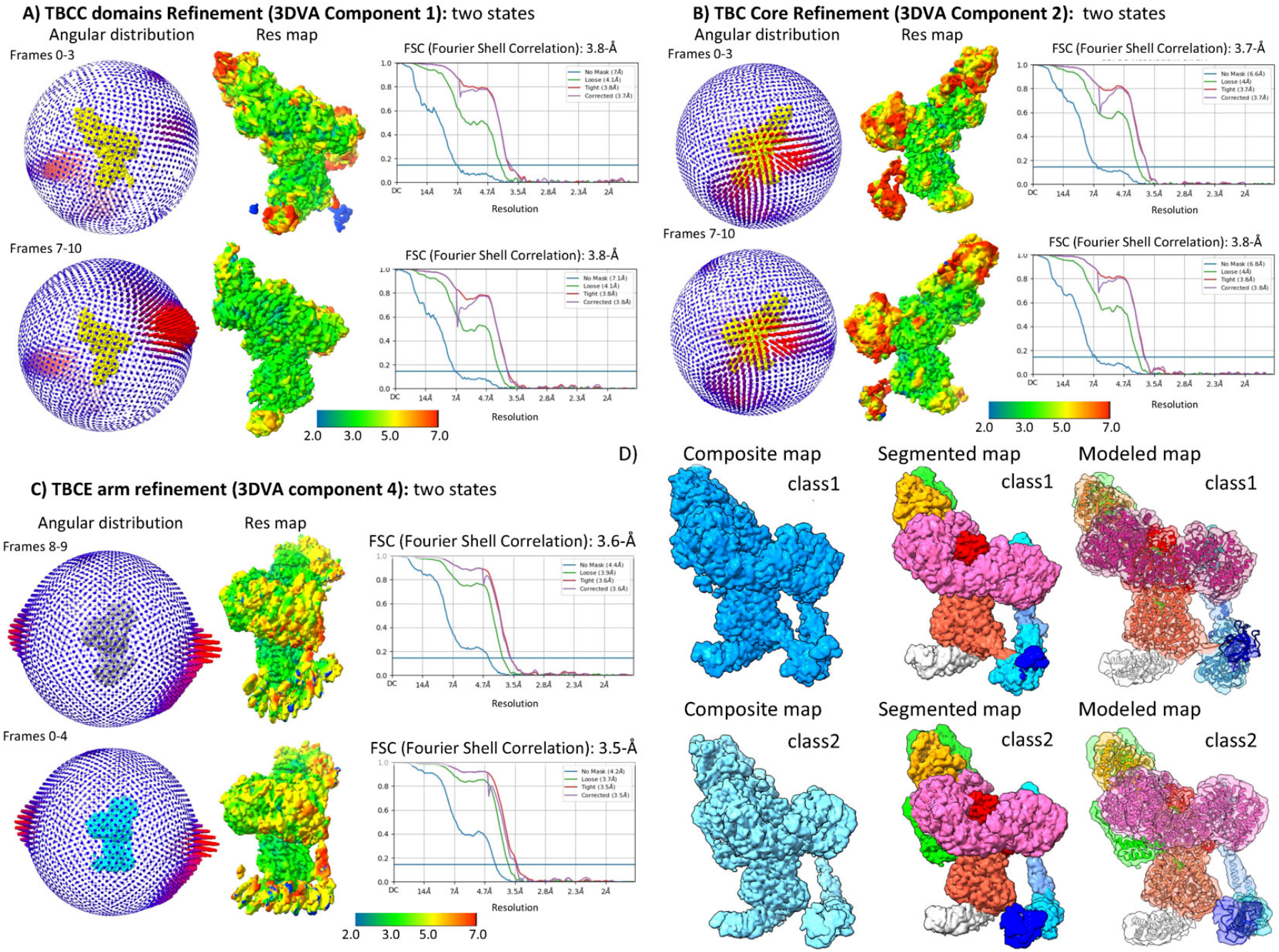
Details of the 3DVA maps used to generate two unique composite states of TBC-DEG/TBCC-αβ-tubulin states. A) Two maps from 3DVA component 1 refining TBCC-Domains: Top panels, Frames 0-3 and Bottom panels, Frames 7-10; Left panels, Angular distribution for particles in the final refined map. Middle panels, Res-map showing the resolution distribution color range onto the structure. Right panels, Fourier Shell Correlation (FSC) for each map. B) Two maps from 3DVA component 2 refining the TBC-DEG core states: Top panels, Frames 0-3 and Bottom panels, Frames 7-10; Left panels, Angular distribution for particles in the final refined map. Middle panels, Res-map showing the resolution distribution color range onto the structure. Right panels, Fourier Shell Correlation (FSC) for each map. C) Two maps from 3DVA component 4 refining the TBCE arm conformation. Top panels, Frames 8-9 and Bottom panels, Frames 0-4; Left panels, Angular distribution for particles in the final refined map. Middle panels, Res-map showing the resolution distribution color range onto the structure. Right panels, Fourier Shell Correlation (FSC) for each map. D) Composite maps for TBC-DEG/TBCC-αβ-tubulin state 1 (class 1) are shown top left, and TBC-DEG/TBCC-αβ-tubulin state 2 (class 2) are shown on the bottom left. Segmented maps for class 1 and class 2 are shown in the middle panels. Modeled maps for class 1 and class 2 are shown in the right panels.

**Figure S9:**
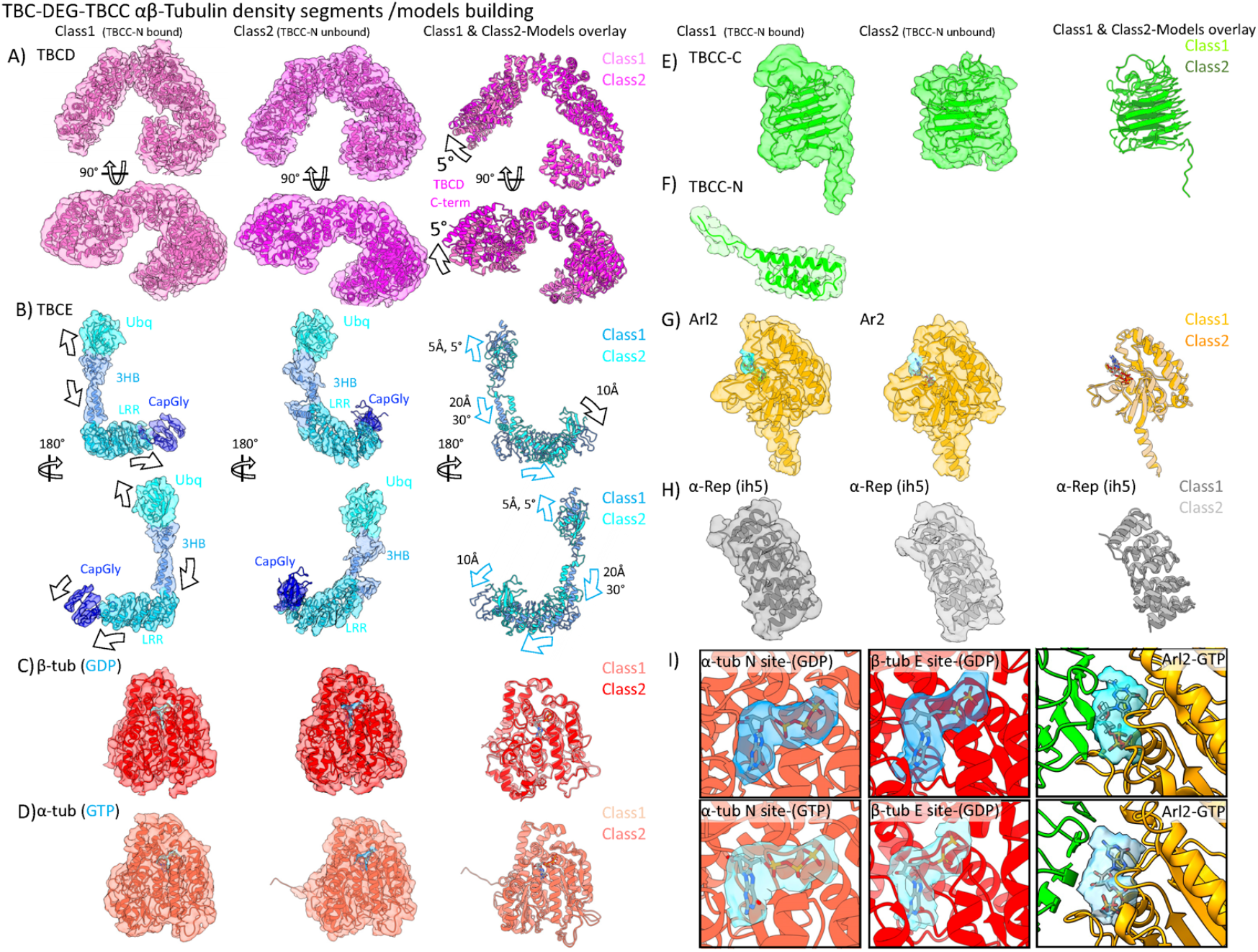
Cryo-EM map segments and atomic models built of each of the TBC-DEG/TBCC-αβ-tubulin subunits. The segments with built models of the class 1 (TBCC-N bound) (left, panel) and class 2(TBCC-N unbound) (middle panel) and overlay of the two-state models (right panel) showing the conformational changes in various subunits A) Top, Map segment and model for TBCD class 1 (pink, left panel), TBCD class 2 (magenta, middle panel), and overlay of the two models (right panel). Bottom, 90°-rotated view of the panels above. B) Map segment and model for TBCE class 1 (colored according to subdomains, left panel), TBCE class 2 (colored according to subdomains middle panel). Conformational changes are marked on various domains. The right panels show an overlay of the two models (class 1 is shown in cyan and class 2 is shown in sky blue). The bottom panels are 180° rotated views of the panels above. C) Map segment and model for β-tubulin class 1 (red, left panel), β-tubulin class 2 (red, middle panel), and overlay of the two models (right panel). E-site GDP densities are shown in cyan, and close-ups are shown in more detail in the middle panels for part I. D) Map segment and model for β-tubulin class 1 (tomato, left panel), β-tubulin class 2 (tomato, middle panel), and overlay of the two models (right panel). N-site GTP nucleotide densities are shown in blue density and shown in more detail in the right panels for part I. E) Map segment and model for TBCC-C class 1 (lime green, left panel), β-tubulin class 2 (lime green, middle panel), and overlay of the two models (right panel). F) Map segment and model for TBCC-N (lime green, left panel). These densities are missing in class 2. G) Map segment and model for Arl2 class 1 (orange, left panel), β-tubulin class 2 (orange, middle panel), and overlay of the two models (right panel). Arl2 GTP nucleotide densities are shown in cyan and close-up views are shown in more detail in the right panels for part I H) Map segment and model for α-Rep iHS in class1 (Grey, left panel), α-Rep iHS in class2 (light gray middle panel), and overlay of the two models (right panel). I) Close-up views of β-tubulin E-site (GTP), α-tubulin N-site (GDP), and Arl2 (GTP) nucleotide densities

**Figure S10:**
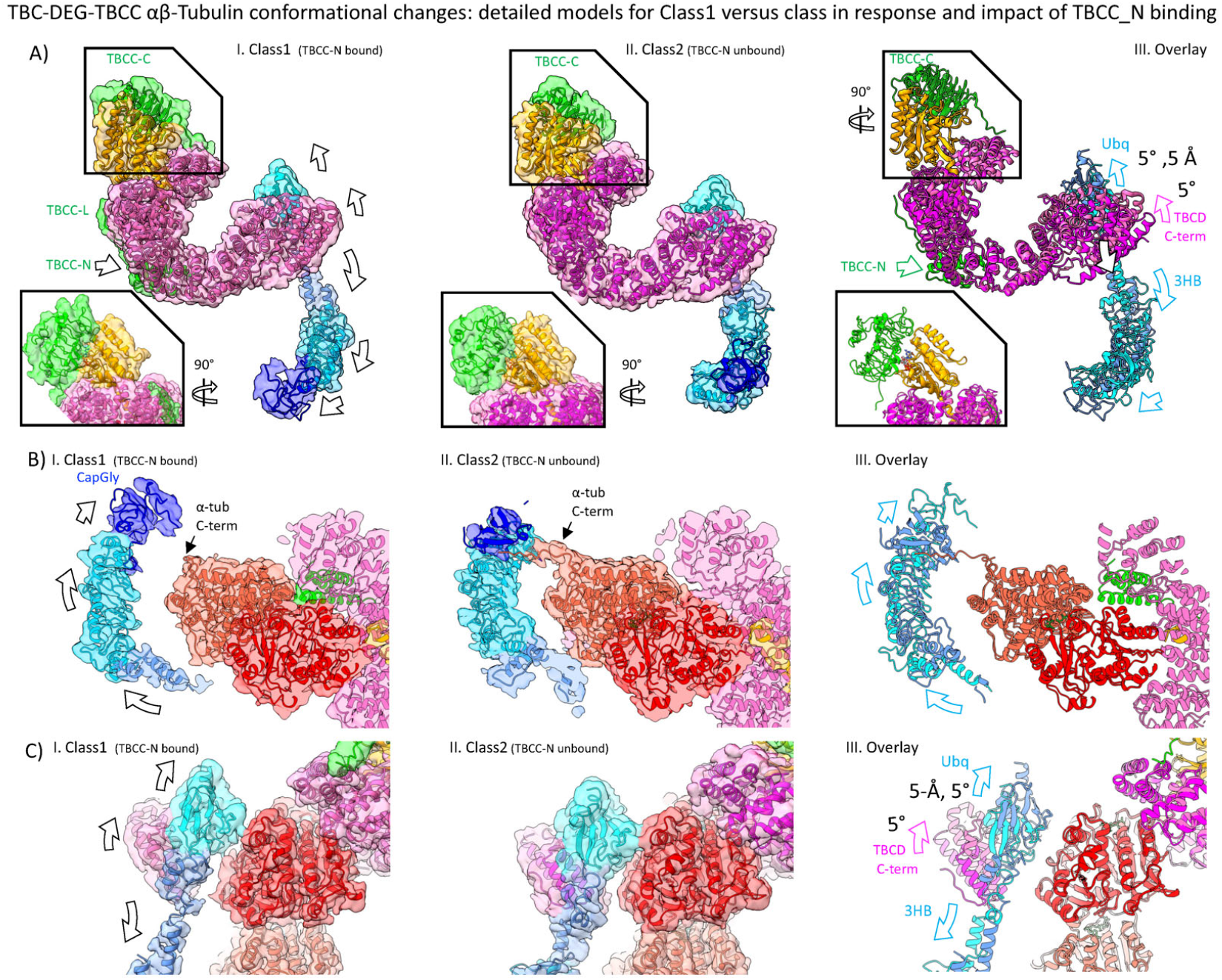
Additional views of TBC-DEG/TBCC with or without αβ-tubulin focusing on the conformational changes in TBCD, TBCE, and their interactions with αβ-tubulin. A) Side View of TBC-DEG/TBCC assembly with αβ-tubulin computationally removed; Left panel (1.) class 1 (TBCC-N bound), marking conformational changes in TBCD and TBCE; middle panel (11.), class 2 (TBCC-N unbound). Right panel (111.), overlay of model for Class 1 and Class 2, marking the conformational changes in TBCD and TBCE. 1nsets for each panel show 90° rotated views of the TBCD/Arl2/TBCC-C region of the complex. B) Slice view of the TBCE LRR-CapGly arm interface with α-tubulin. Left panel (1.), Class1 (TBCC-N bound); right panel (11.), class 2 (TBCC-N unbound) showing the TBCE CapGly α-tubulin C-terminus interface. right panel (111.), overlay of models for class 1 and class 2, marking the change in the conformation in the TBCE LRR-CapGly arm. C) Back view of the TBCD-spiral/TBCE Ubq interface with β-tubulin revealing the conformational changes in the Ubq domain and conformational change in 3HB of TBCE. Left Panel (1.), shows class 1 (TBCC-N bound) showing the TBCE Ubq in the upper position. Middle panel (11.), shows class 2 (TBCC-N unbound) showing the TBCE Ubq in the lower position. The right panel (111.), shows the overlay of Class 1 and Class 2 marking conformational changes in TBCD spiral and TBCE Ubq domains and their impacts on β-tubulin binding and 3HB changes.

**Figure S11:**
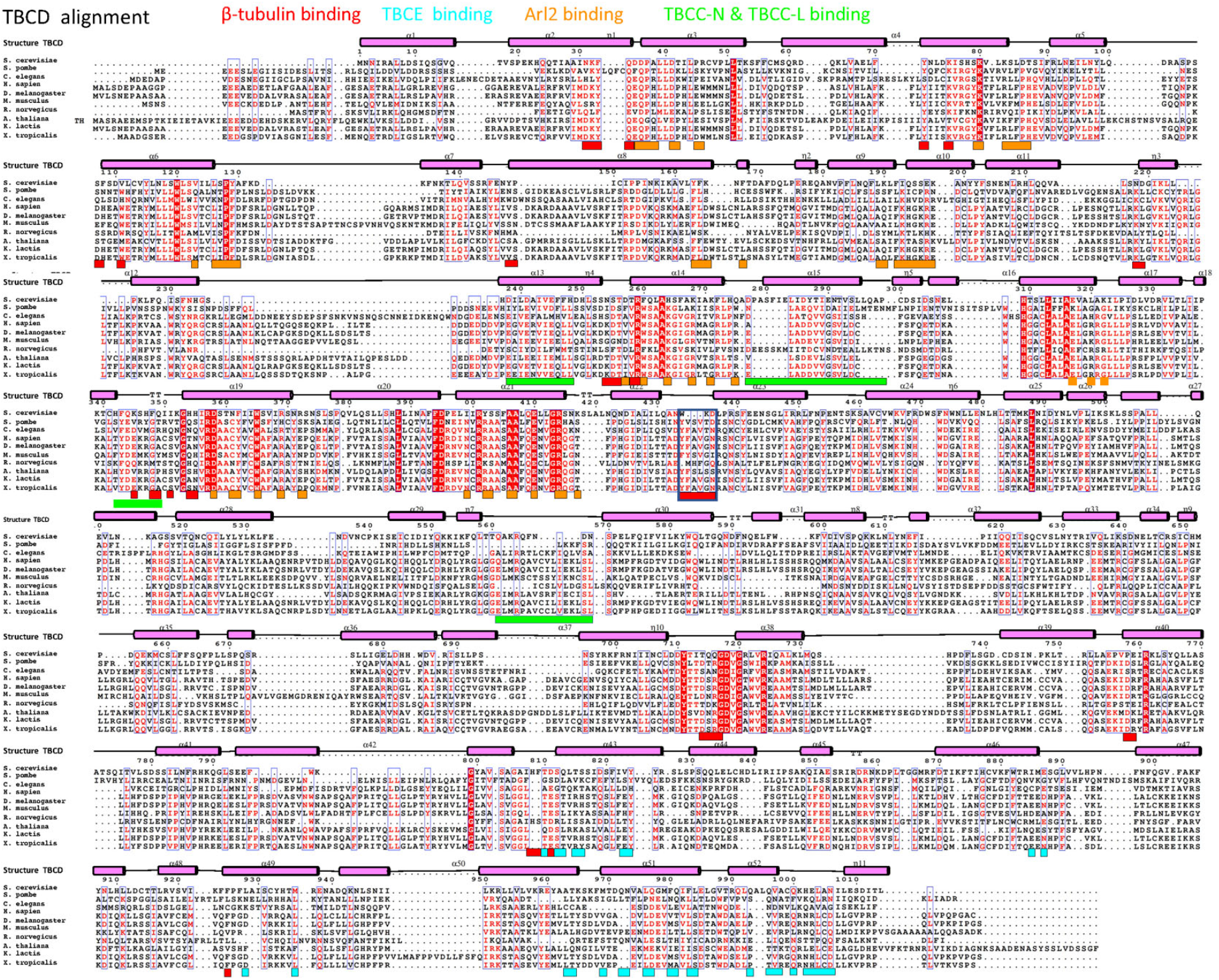
TBCD sequence alignment, secondary structure elements, and subunit interaction sites. Multiple sequence alignment of TBCD sequences with the secondary structure element boundaries marked on top of the sequences. The interacting residues for each region are marked below with the colored panels of the interacting subunits.

**Figure S12:**
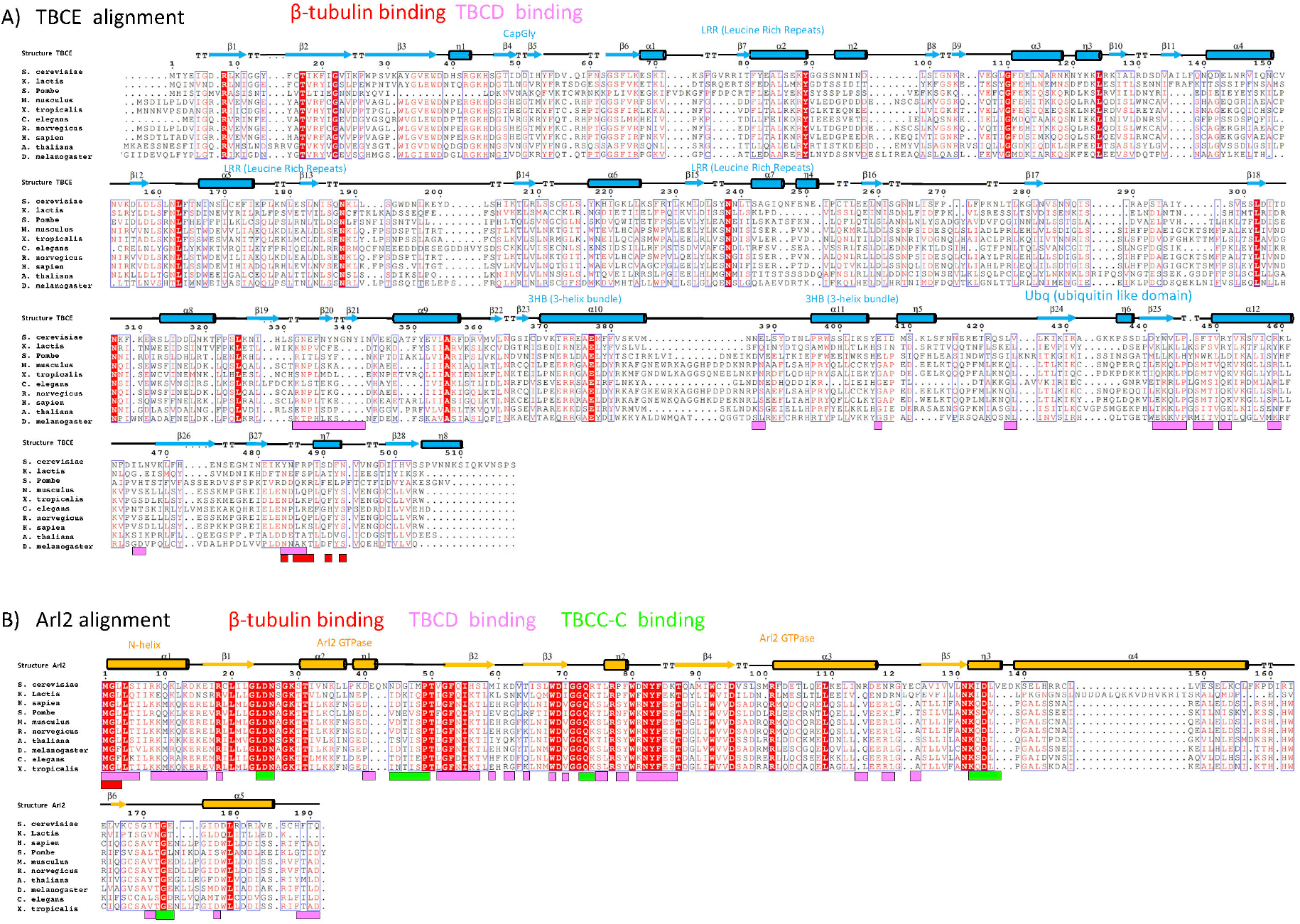
TBCE and Arl2 sequence alignments, secondary structure elements, subunit interaction sites. A) Multiple sequence alignment of TBCE orthologs with the secondary structure element boundaries marked on top of the sequences. The interacting residues for each region are marked below with the colored panels of the interacting subunits. B) Multiple sequence alignment of Arl2 orthologs with the secondary structure element boundaries marked on top of the sequences. The interacting residues for each region are marked below with the colored panels of the interacting subunits.

**Figure S13:**
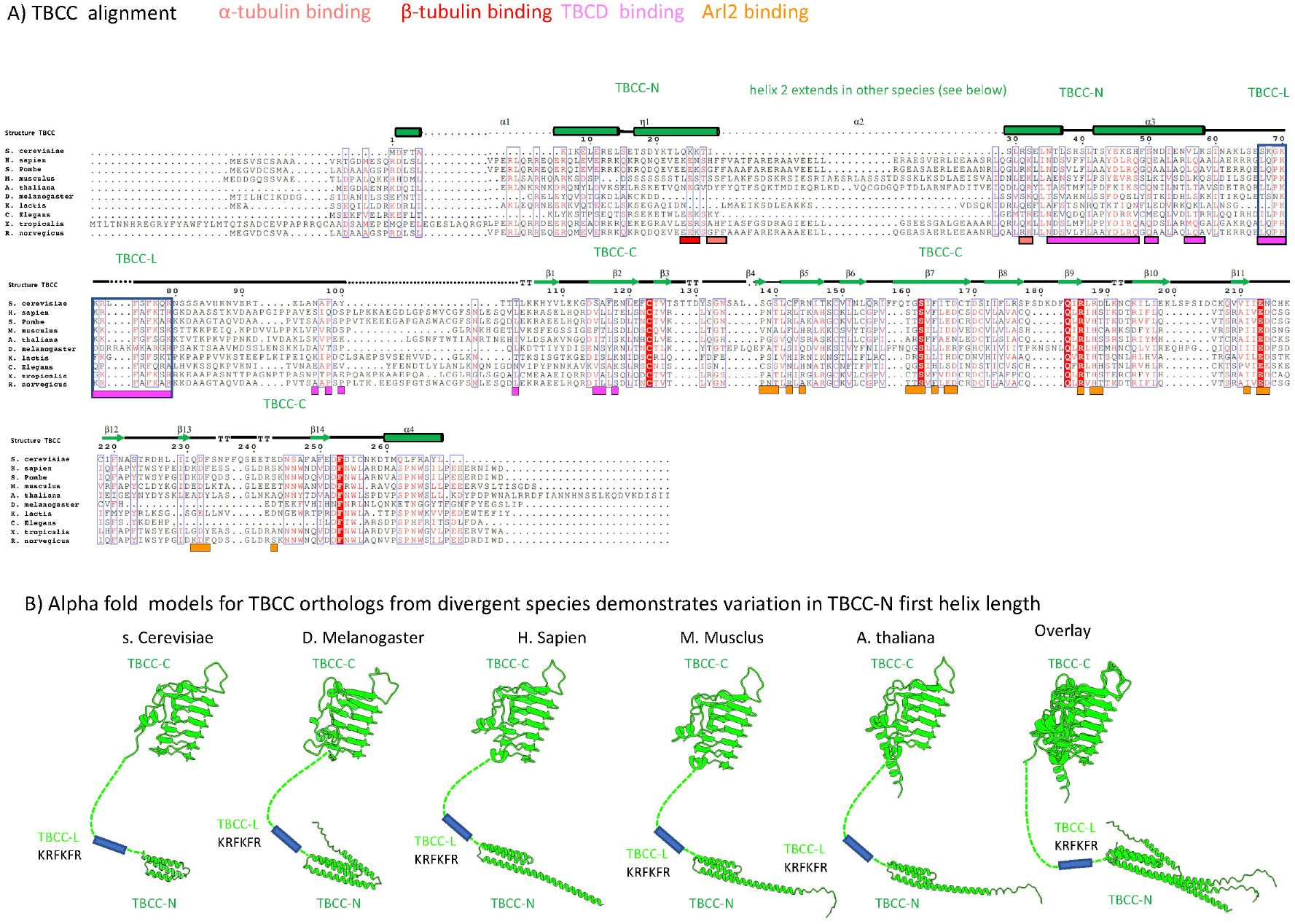
TBCC sequence alignment, structure boundaries, subunit interaction sites, and models for TBCC domains across species showing variations in TBCC-N first helical segment length. A) Multiple sequence alignment of TBCC orthologs with the secondary structure element boundaries marked on top of the sequences. The interacting residues for each region are marked below with the colored panels of the interacting subunits. B) Alphafold2 prediction models for TBCC orthologs from multiple organisms show the structural conservation in TBCC-N and TBCC-C. TBCC-N helix 1 varies dramatically in length across various species compared to its short length in S. Cerevisiae.

**Figure S14:**
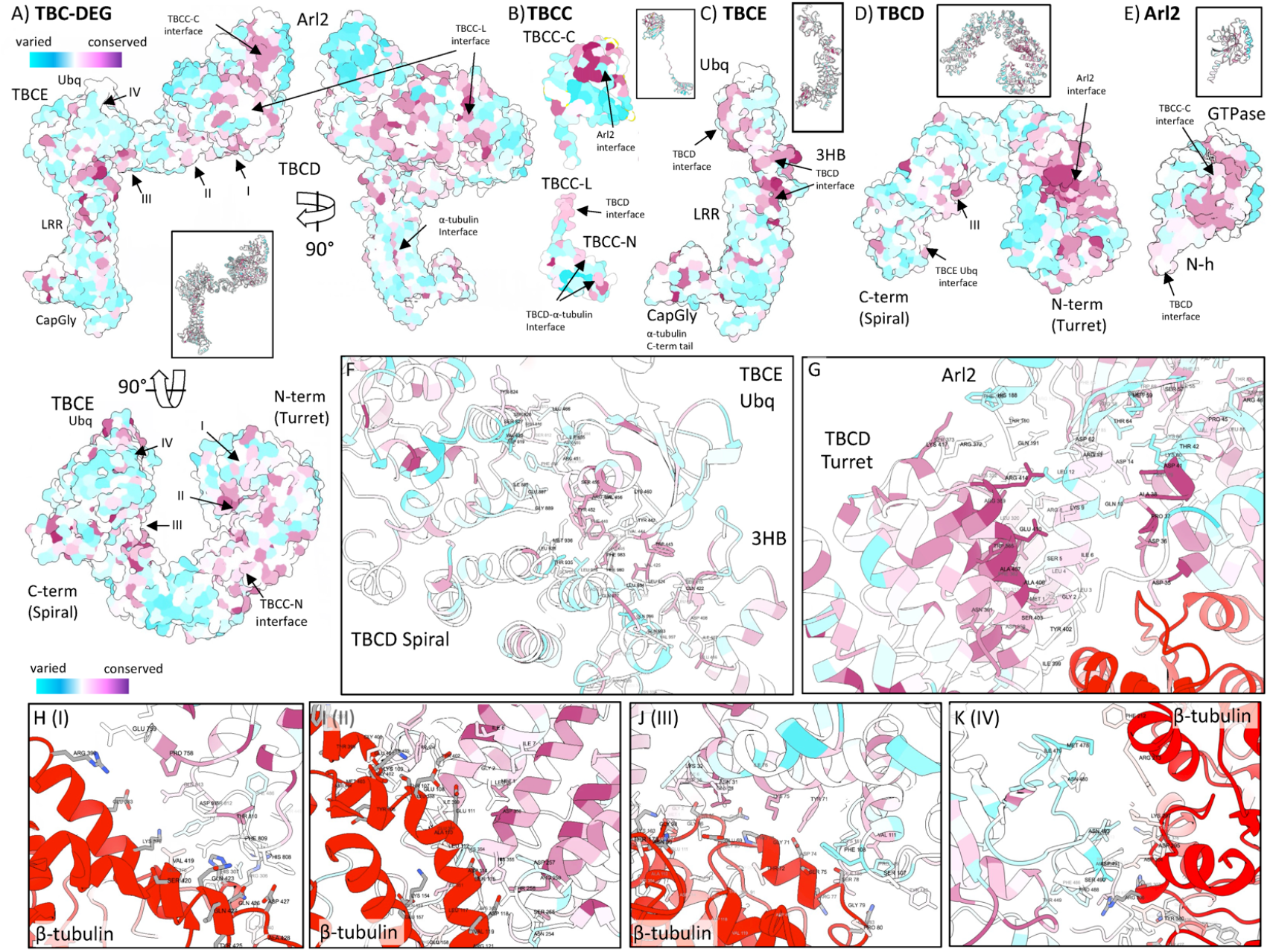
Conserved surfaces of TBC-DEG, TBCD, TBCE, Arl2, and TBCC structures: conservation in interfaces forming the TBC-DEG assembly and interactions with TBCC, and αβ-tubulin subunits. A) Three 90°-rotated views of surface representations of the TBC-DEG assembly (αβ-tubulin removed) with the conserved to varied residues colored from purple to blue, based on the scale shown on the top left. The inset shows the structure view in ribbon format. Highlighted regions include: Three TBCD binding interfaces for β-tubulin (I, II, III), TBCC binding interface on the Arl2 GTPase surface, TBCC-L and TBCC-N binding interfaces on TBCD-spiral region B) A Surface representation of TBCC domains TBCC-N and TBCC-C showing conservation in binding Arl2 interface and TBCC-L and TBCC-N binding interfaces for TBCD spiral domain C) A Surface representation of TBCE showing surface conservation. The inset shows the structure view in ribbon format. Highlighted are the Ubq domain and 3HB binding interfaces to TBCD. D) Top view of a surface representation of TBCD showing surface conservation. The inset shows the structure in ribbon format. Highlighted are TBCD surface-conserved interfaces for binding β-tubulin (I, II, III), Arl2, and TBCE. E) Two 90° views of surface representation of Arl2 showing surface conservation. The inset shows the structure view in ribbon format. Highlighted are the surface-conserved interfaces for TBCD F) Close-up view of the TBCD Spiral-TBCE Ubq interface with both TBCD and TBCE side chains colored to show conservation, shown in a similar view to Figure 2E. A) Close-up view of the TBCD Turret-Arl2 interface with both TBCD and TBCE side chains colored to show conservation, shown in a similar view to Figure 2C and 2D. H-K) Close-up view of the β-tubulin interfaces (I, II, III) TBCD and TBCE-Ubq (IV) interfaces shown in similar views to Figure 3G-H, showing the conservation with β-tubulin.

**Figure S15:**
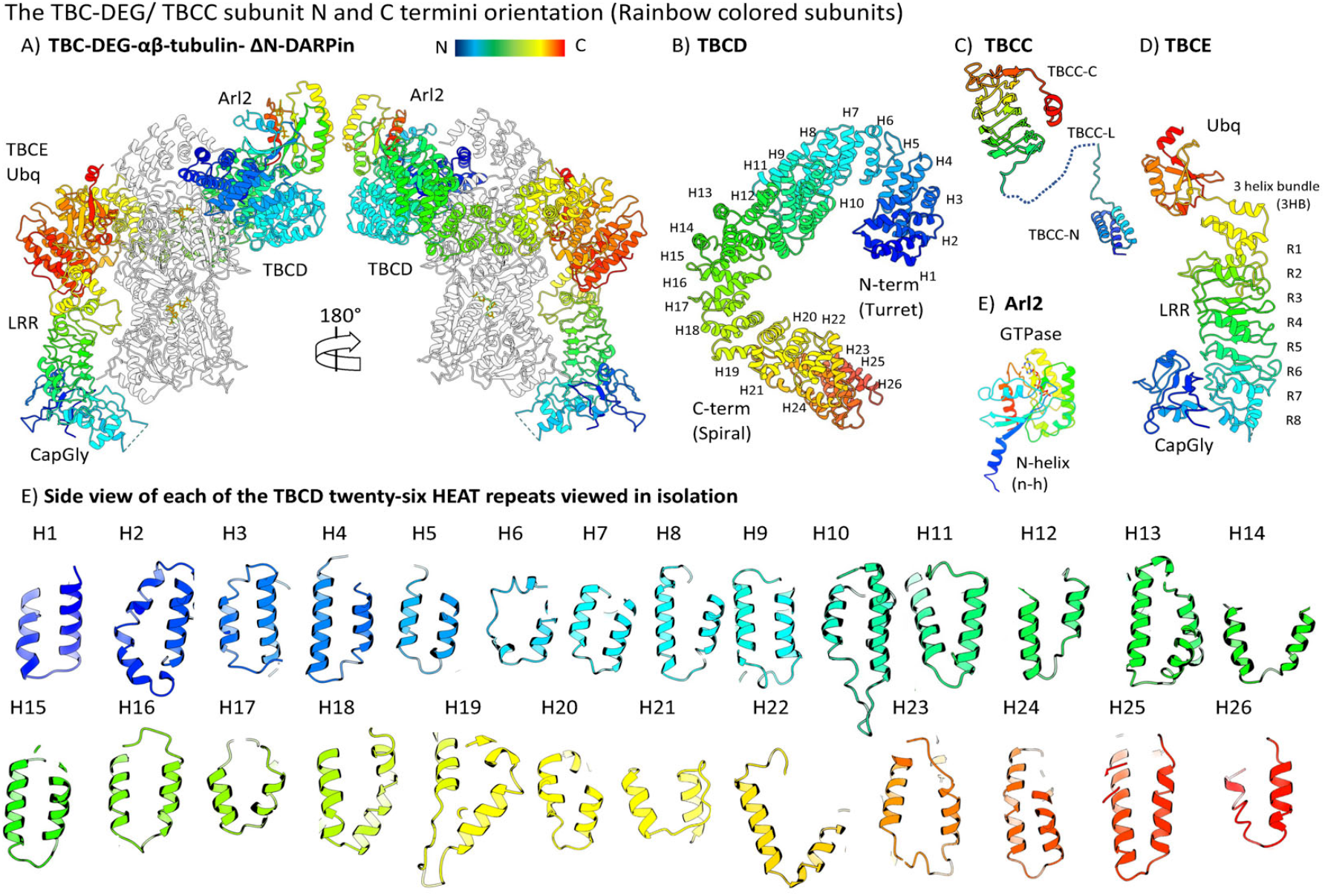
Rainbow ribbon representation of TBCD, TBCE, Arl2, and TBCC subunits and their interactions within TBC-DEG and dissociated TBCD HEAT repeats revealing their overall length and organization in TBCD. A) Two 90°-rotated views of TBC-DEG-αβ-tubulin assembly with αβ-tubulin and ΔN-DARPin are shown in white. TBCD, TBCE, and Arl2 are shown in rainbow-colored ribbon format with a palette described according to the insert. B) Top view of TBCD in ribbon rainbow color representation with its 26 HEAT repeats marked. Note E shows these HEAT repeat in isolation C) The side view of TBCC in the rainbow color representation shows its three domains. D) Side view of TBCE in ribbon rainbow color representation, showing its four domains marked. With LRR repeats marked R1-R8 E) Side view of Arl2 in ribbon rainbow color representation, showing its two domains. F) Side view of each of the TBCD twenty-six HEAT repeats shown in isolation, side by side, using colors shown in panel B.

**Figure S16:**
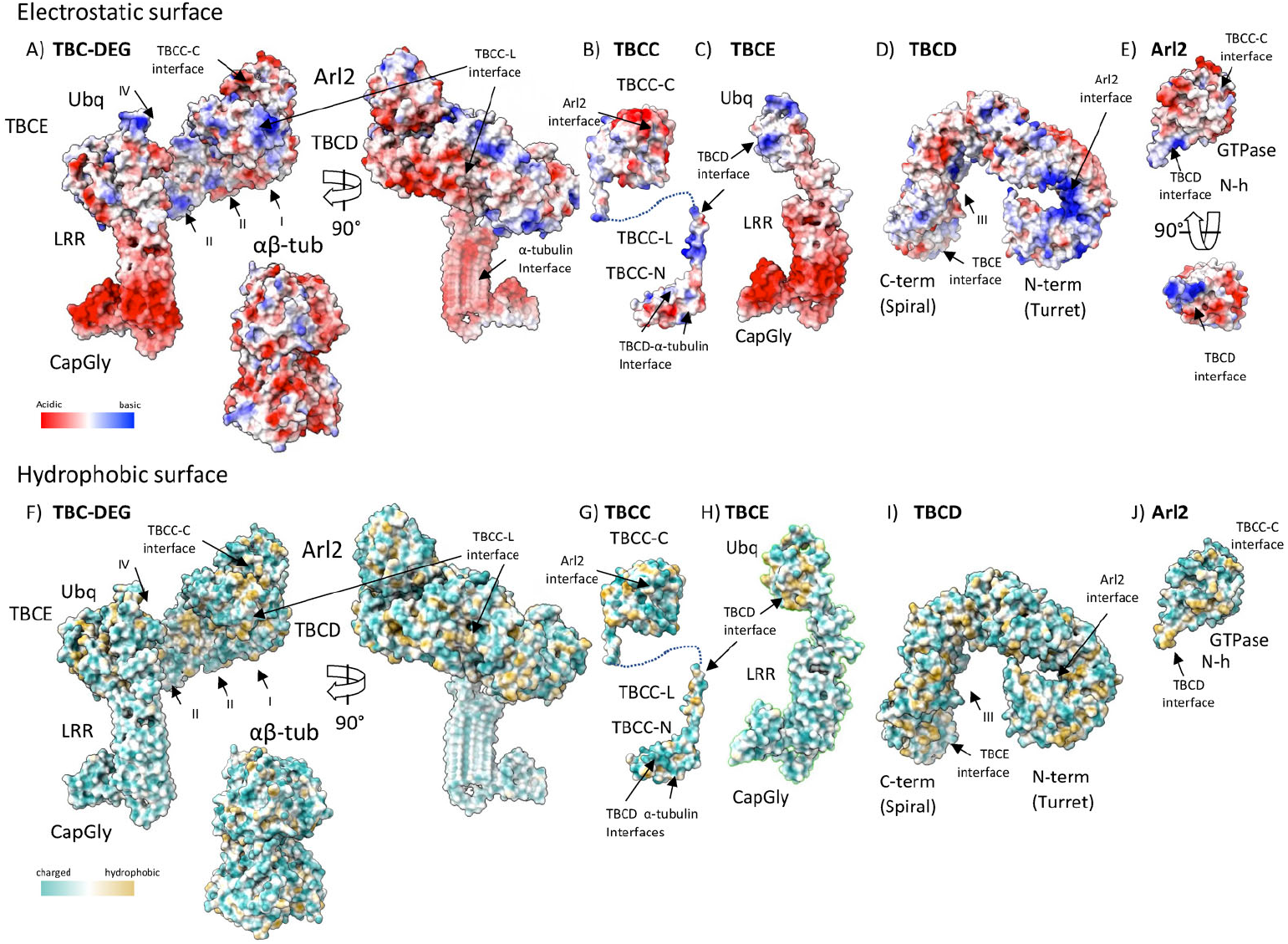
Electrostatic and hydrophobic surface representation of TBC-DEG, TBCD, TBCE, Arl2, and TBCC subunits showing the nature of the charged or hydrophobic interaction sites. A) Two 90°-rotated views of an electrostatic surface representation of the TBC-DEG assembly colored according to the insert shown on the bottom left. The β-tubulin-binding interfaces (I, II, III, IV) and the TBCC-L interface are highlighted. The isolated αβ-tubulin in a similar color representation is also shown in a similar orientation to the left panel. B) Side-view electrostatic surface representation of TBCC with its three domains shown. The Arl2, TBCD, and α-tubulin interfaces are highlighted. C) A side-view electrostatic surface representation of TBCE with its four domains is shown. The TBCD, β-tubulin, and α-tubulin interfaces are highlighted. D) The top-view electrostatic surface representation of TBCD is shown. The β-tubulin (I, II, III), Arl2, and TBCE interfaces are highlighted. E) Two 90°-rotated electrostatic surface representations of Arl2 with its two domains are shown. The TBCD and TBCC interfaces are highlighted. F) Two 90°-rotated views of the hydrophobic charged surface representation of the TBC-DEG assembly colored according to the insert shown on the bottom left. The β-tubulin-binding interfaces (I, II, III, IV) and the TBCC-L interface are highlighted. The isolated αβ-tubulin in a similar color representation is also shown in a similar orientation to the left panel. G) Side-view hydrophobic charged surface representation of TBCC with its three domains shown. The Arl2, TBCD, and α-tubulin interfaces are highlighted. H) A side-view hydrophobic charged surface representation of TBCE with its four domains is shown. The TBCD, β-tubulin, and α-tubulin interfaces are highlighted. I) The top-view hydrophobic charged surface representation of TBCD is shown. The β-tubulin (1, 11, 111), Arl2, and TBCE interfaces are highlighted. J) Two 90°-rotated hydrophobic charged surface representations of Arl2 with its two domains are shown. The TBCD and TBCC interfaces are highlighted.

