## Extended Data Figures S1-S16 for "Cryo-EM structures of the tubulin cofactors reveal the molecular basis for the biogenesis of alpha/beta-tubulin"

bioRxiv
THE PREPRINT SERVER FOR BIOLOGY

### Cryo-EM structures of the tubulin cofactors reveal the molecular basis for the biogenesis of alpha/beta-tubulin

**Aryan Taheri[1,3], Zhoaqian Wang[1,4] *, Bharti Singal[1,5] *, Fei Guo[1], Jawdat Al-Bassam[1]**

[1]Molecular Cellular Biology Department, University of California, Davis, CA, USA

* These authors contributed equally to the work.

[3] Present address: Molecular Cell Biology Department, University of California, Berkeley, CA, USA
[4] Present address: Biochemistry Department, University of Washington, Seattle, WA, USA
[5] Present address: Stanford Cryo-EM microscopy Center, Stanford University, Palo Alto, CA, USA

#### Extended Data figures

**Extended Data Figures Legends**

**Figure S1**: **Purification of recombinant TBC-DEG and Reconstitution of the TBC-DEG-αβ-tubulin assemblies for cryo-EM**

C) Left, Angular distribution for TBC-DEG-$\alpha\beta$-tubulin class1 particles in the final refined map. Middle panel, Resmap for TBC-DEG-$\alpha\beta$-tubulin in class 1 showing the resolution distribution color range onto the structure.
D) Left, Angular distribution for TBC-DEG-$\alpha\beta$-tubulin class 2 particles in the final refined map. Middle panel, Resmap for TBC-DEG-$\alpha\beta$-tubulin in class 2 showing the resolution distribution color range onto the structure.
E) Gold standard Fourier Shell correlation (FSC) for TBC-DEG-$\alpha\beta$-tubulin class 1.
F) Gold standard Fourier Shell correlation (FSC) for TBC-DEG-$\alpha\beta$-tubulin class 2.
G) Left, Model-based segmented TBC-DEG-$\alpha\beta$-tubulin-$\Delta$N-DARPin class1 map (segmented map), right, atomic models for subunits placed in segments of these subunits of TBC-DEG-$\alpha\beta$-tubulin-$\Delta$N-DARPin.
H) Left, Model-based segmented TBC-DEG-$\alpha\beta$-tubulin-$\Delta$N-DARPin class2 map (segmented map), right, atomic models for subunits placed in segments of these subunits of TBC-DEG-$\alpha\beta$-tubulin-$\Delta$N-DARPin (modeled map).

**Figure S4**: **Cryo-EM map segments and atomic models built of each of the TBC-DEG-$\alpha\beta$-tubulin subunits including two unique TBCE models and details of the unique TBCE-$\alpha\beta$-tubulin interfaces in the two states.**

A) Left panel, Map segment, and Model built for TBCD. right panel, for example, density for an atomic model built into the TBCD electron density.
B) Left panel, Map segment, and Model built for Arl2. middle panel, example density for an atomic model built into the Arl2 electron density with GTP shown in red. Right panel close-up view for GTP in Arl2 with electron density shown in Red.
C) Left panel, Map segment, and Model built for $\beta$-tubulin. middle panel, example density for an atomic model built into the $\beta$-tubulin electron density with E-site GDP shown in blue. Right panel close-up view for E-site GDP in $\beta$-tubulin with electron density shown in blue.
D) Left panel, Map segment, and Model built for $\alpha$-tubulin. middle panel, example density for an atomic model built into the A-tubulin electron density with the N-site GTP shown in blue. Right panel close-up view for N-site GTP in $\alpha$--tubulin with electron density shown in blue.
E) Left panel, Map segment, and Model built for $\Delta$N-DARPin. right panel, for example, density for an atomic model built into the $\Delta$N-DARPin electron density.
F) Left panels, Class 1 TBCE state 1 showing segmented density map (left, sky blue), model fitted in subregion segmented map (right). Right panels, Class 1 TBCE state 2 showing segmented density map (left, cyan), model fitted into subregion segmented map (right).
G) Left panels, class1 vs class1 map comparison showing a slice view of TBCE-$\alpha\beta$-tubulin interface showing the class1 TBCE retracted state (class1) versus the TBCE-$\alpha$-tubulin bound state (class 2). Right panels, class 1 (left) vs class 2 (right) density map viewed with the $\alpha\beta$-tubulin interaction interfaces marked by the color of $\alpha$-tubulin (light red) and $\beta$-tubulin (dark red) showing changes in the TBCE LRR in interfacing with $\alpha$-tubulin in Class2 compared to class 1

**Figure S10**: **Additional views of TBC-DEG/TBCC with or without αβ-tubulin focusing on the conformational changes in TBCD, TBCE, and their interactions with αβ-tubulin.**

A) Side View of TBC-DEG/TBCC assembly with αβ-tubulin computationally removed; Left panel (I.) class 1 (TBCC-N bound), marking conformational changes in TBCD and TBCE; middle panel (II.), class 2 (TBCC-N unbound). Right panel (III.), overlay of model for Class 1 and Class 2, marking the conformational changes in TBCD and TBCE. Insets for each panel show 90° rotated views of the TBCD/Arl2/TBCC-C region of the complex.
B) Slice view of the TBCE LRR-CapGly arm interface with α-tubulin. Left panel (I.), Class1 (TBCC-N bound); right panel (II.), class 2 (TBCC-N unbound) showing the TBCE CapGly α-tubulin C-terminus interface. right panel (III.), overlay of models for class 1 and class 2, marking the change in the conformation in the TBCE LRR-CapGly arm.
C) Back view of the TBCD-spiral/TBCE Ubq interface with β-tubulin revealing the conformational changes in the Ubq domain and conformational change in 3HB of TBCE. Left Panel (I.), shows class 1 (TBCC-N bound) showing the TBCE Ubq in the upper position. Middle panel (II.), shows class 2 (TBCC-N unbound) showing the TBCE Ubq in the lower position. The right panel (III.), shows the overlay of Class 1 and Class 2

marking conformational changes in TBCD spiral and TBCE Ubq domains and their impacts on β-tubulin binding and 3HB changes.

H) A side-view hydrophobic charged surface representation of TBCE with its four domains is shown. The TBCD, $\beta$-tubulin, and $\alpha$-tubulin interfaces are highlighted.

I) The top-view hydrophobic charged surface representation of TBCD is shown. The $\beta$-tubulin (I, II, III), Arl2, and TBCE interfaces are highlighted.

J) Two 90$^\circ$-rotated hydrophobic charged surface representations of Arl2 with its two domains are shown. The TBCD and TBCC interfaces are highlighted.

Figure S1

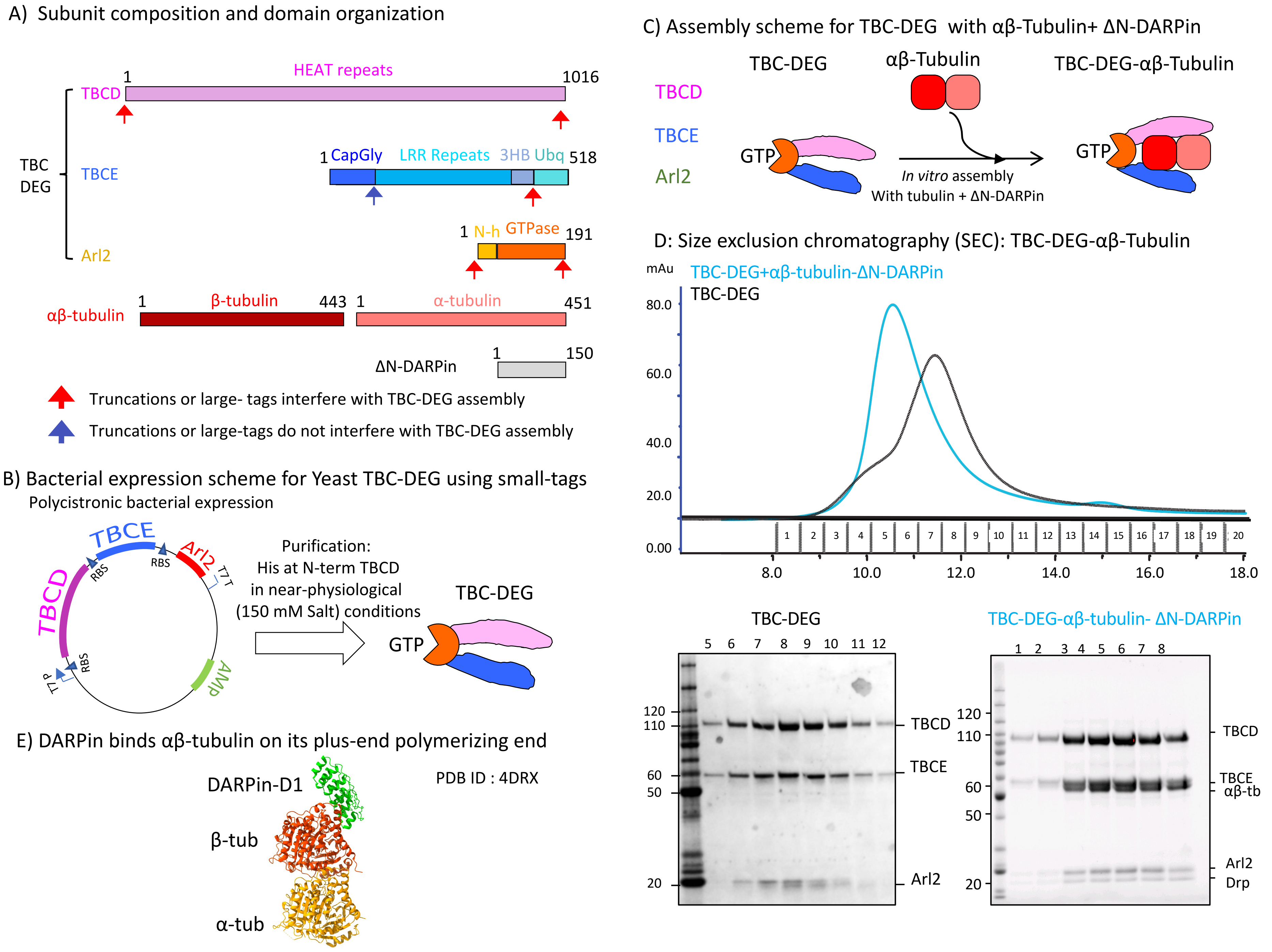

A) Subunit composition and domain organization
HEAT repeats
1
1016
TBCD
TBC
DEG
TBCE
1 CapGly LRR Repeats 3HB Ubq 518
1 N-h GTPase 191
Arl2
αβ-tubulin
1 β-tubulin 443
1 α-tubulin 451
ΔN-DARPin
1 150
Truncations or large- tags interfere with TBC-DEG assembly
Truncations or large-tags do not interfere with TBC-DEG assembly
B) Bacterial expression scheme for Yeast TBC-DEG using small-tags
Polycistronic bacterial expression
TBCD
TBCE
Arl2
RBS
RBS
RBS
T7 T
T7 P
AMP
Purification:
His at N-term TBCD
in near-physiological
(150 mM Salt) conditions
TBC-DEG
GTP
C) Assembly scheme for TBC-DEG with αβ-Tubulin+ ΔN-DARPin
TBC-DEG
αβ-Tubulin
TBC-DEG-αβ-Tubulin
TBCD
TBCE
Arl2
GTP
In vitro assembly
With tubulin + ΔN-DARPin
GTP
D: Size exclusion chromatography (SEC): TBC-DEG-αβ-Tubulin
mAu
TBC-DEG+αβ-tubulin-ΔN-DARPin
TBC-DEG
80.0
60.0
40.0
20.0
0.00
1 2 3 4 5 6 7 8 9 10 11 12 13 14 15 16 17 18 19 20
8.0
10.0
12.0
14.0
16.0
18.0
TBC-DEG
5 6 7 8 9 10 11 12
120
110
60
50
20
TBCD
TBCE
Arl2
TBC-DEG-αβ-tubulin- ΔN-DARPin
1 2 3 4 5 6 7 8
120
110
60
50
20
TBCD
TBCE
αβ-tb
Arl2
Drp
E) DARPin binds αβ-tubulin on its plus-end polymerizing end
DARPin-D1
PDB ID : 4DRX
β-tub
α-tub

TBC-DEG-αβ-tubulin Cryo-EM data collection and processing scheme

Figure S2

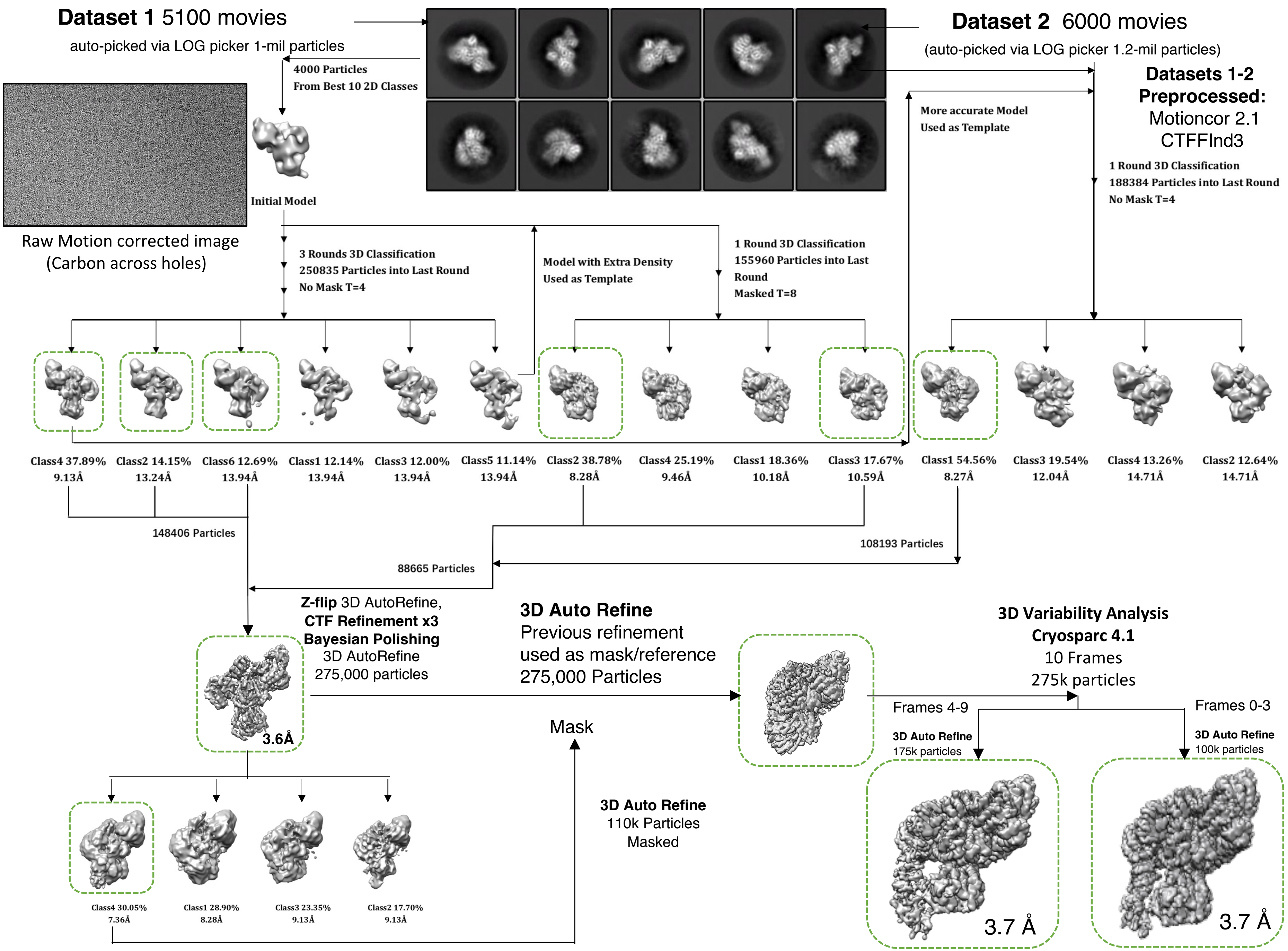

Figure S3

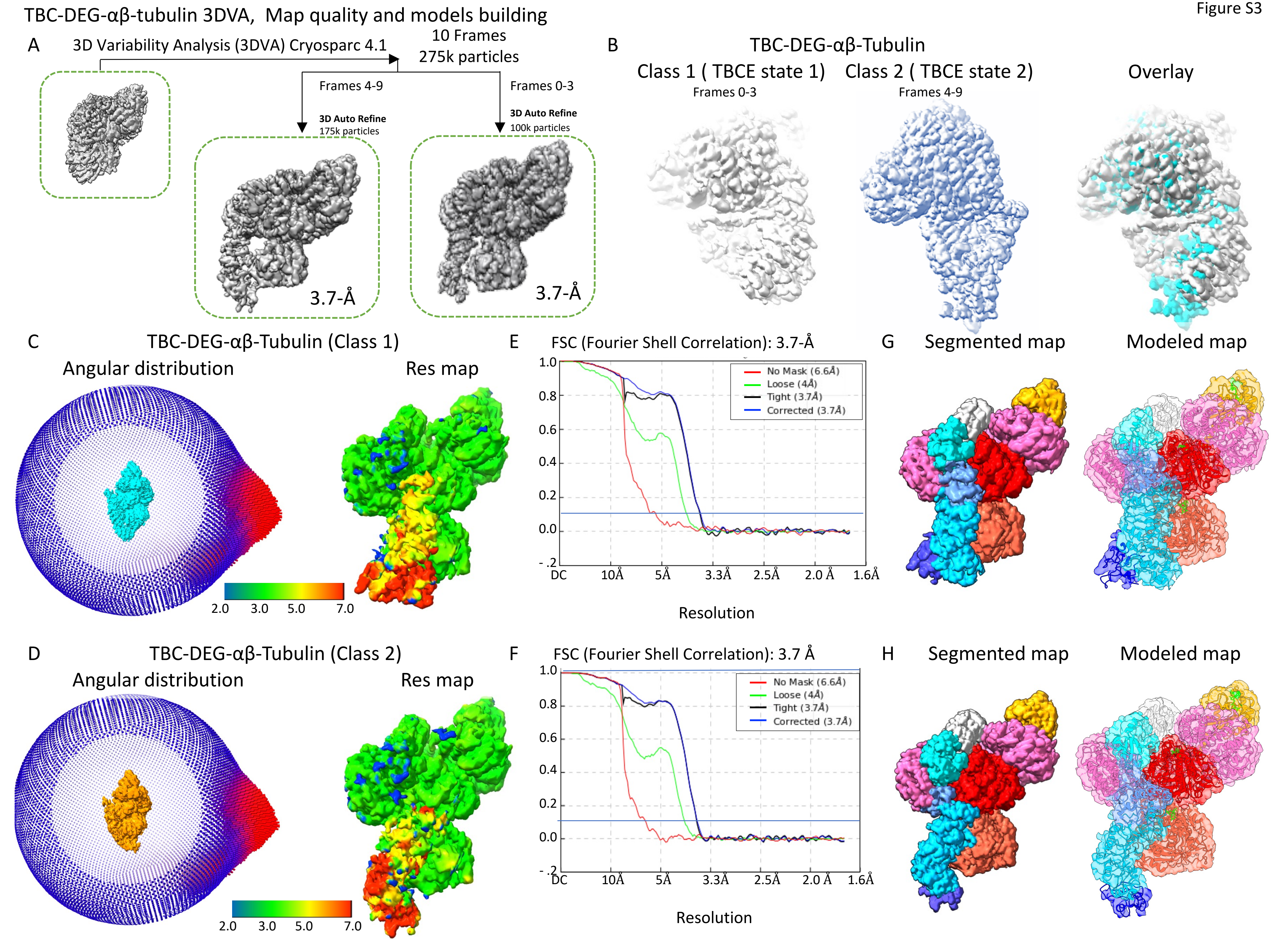

TBC-DEG-αβ-tubulin 3DVA, Map quality and models building
A
3D Variability Analysis (3DVA) Cryosparc 4.1
10 Frames
275k particles
Frames 4-9
Frames 0-3
3D Auto Refine
175k particles
3D Auto Refine
100k particles
3.7-Å
3.7-Å
B
TBC-DEG-αβ-Tubulin
Class 1 ( TBCE state 1)
Frames 0-3
Class 2 ( TBCE state 2)
Frames 4-9
Overlay
C
TBC-DEG-αβ-Tubulin (Class 1)
Angular distribution
Res map
2.0 3.0 5.0 7.0
E
FSC (Fourier Shell Correlation): 3.7-Å
No Mask (6.6Å)
Loose (4Å)
Tight (3.7Å)
Corrected (3.7Å)
1.0 0.8 0.6 0.4 0.2 0.0 -.2
DC 10Å 5Å 3.3Å 2.5Å 2.0 Å 1.6Å
Resolution
G
Segmented map
Modeled map
D
TBC-DEG-αβ-Tubulin (Class 2)
Angular distribution
Res map
2.0 3.0 5.0 7.0
F
FSC (Fourier Shell Correlation): 3.7 Å
No Mask (6.6Å)
Loose (4Å)
Tight (3.7Å)
Corrected (3.7Å)
1.0 0.8 0.6 0.4 0.2 0.0 -.2
DC 10Å 5Å 3.3Å 2.5Å 2.0 Å 1.6Å
Resolution
H
Segmented map
Modeled map

Figure S4

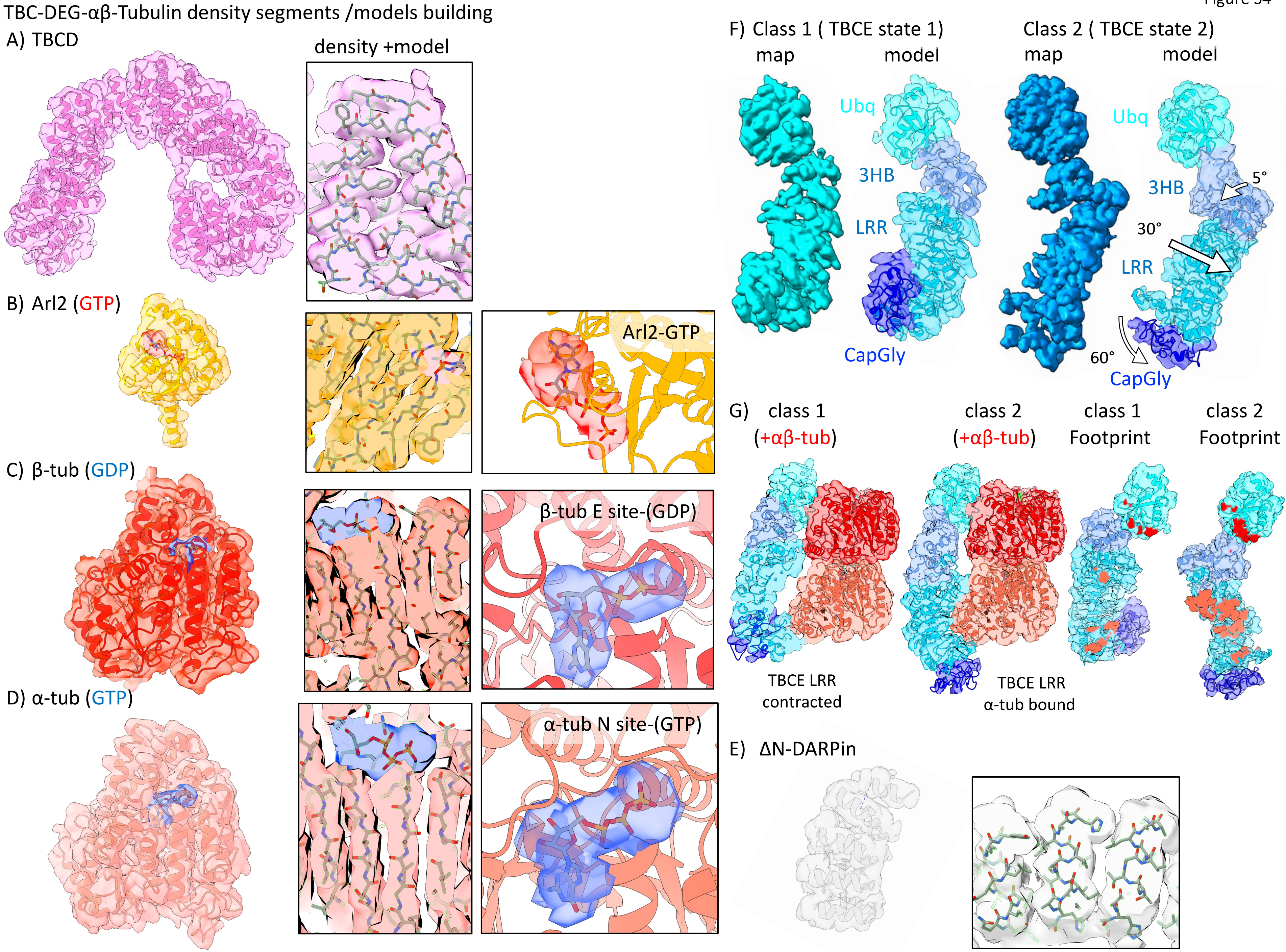
TBC-DEG-αβ-Tubulin density segments /models building
A) TBCD
density +model
B) Arl2 (GTP)
Arl2-GTP
C) β-tub (GDP)
β-tub E site-(GDP)
D) α-tub (GTP)
α-tub N site-(GTP)
F) Class 1 ( TBCE state 1)
map
model
Class 2 ( TBCE state 2)
map
model
Ubq
3HB
LRR
CapGly
Ubq
3HB
5°
30°
LRR
60°
CapGly
G) class 1 (+αβ-tub)
class 2 (+αβ-tub)
class 1 Footprint
class 2 Footprint
TBCE LRR contracted
TBCE LRR α-tub bound
E) ΔN-DARPin

Figure S5

A) Subunit composition and domains organization

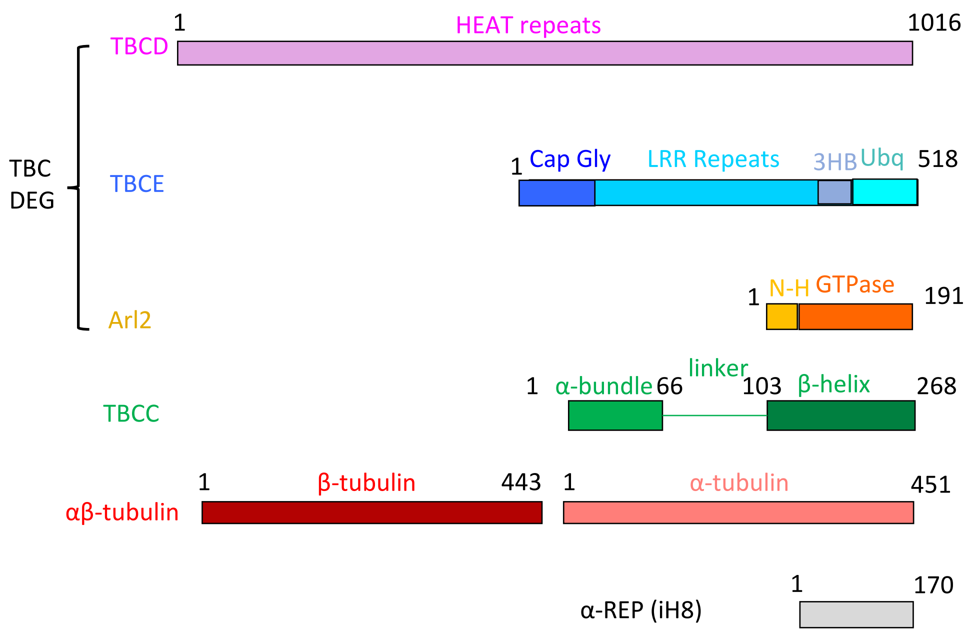


B) reconstitution scheme for TBC-DEG/TBCC-αβ-Tubulin-iH5

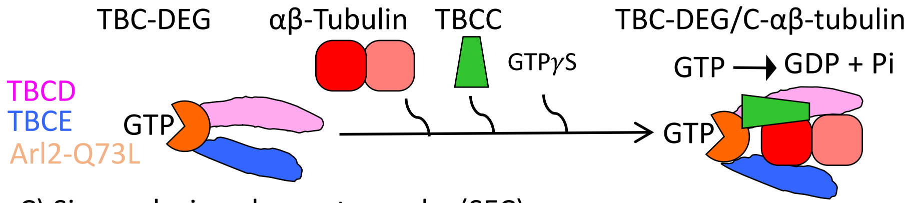


C) Size exclusion chromatography (SEC)

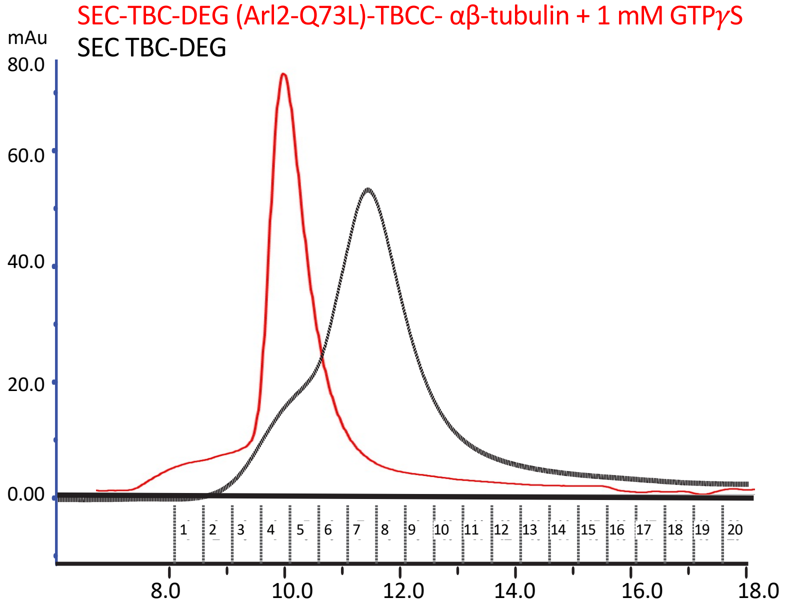


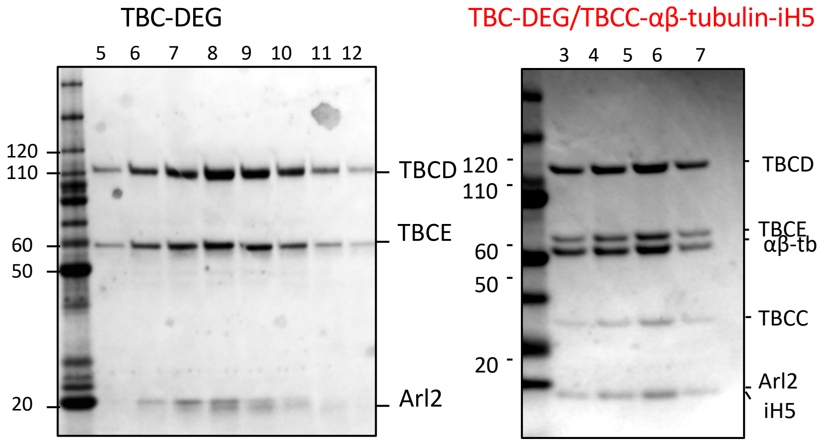


D) the α-rep (iH5) binds αβ-tubulin on its minus-end

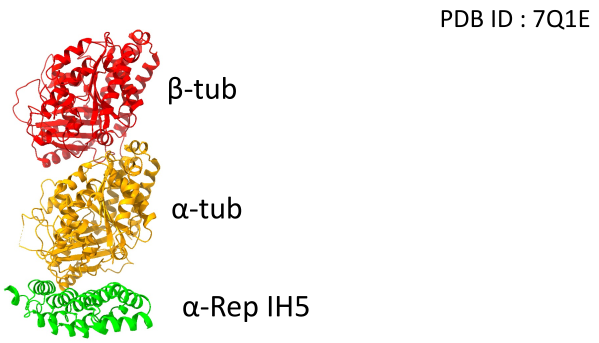

TBC-DEG/TBCC-αβ-tubulin Cryo-EM data collection and processing scheme

Figure S6

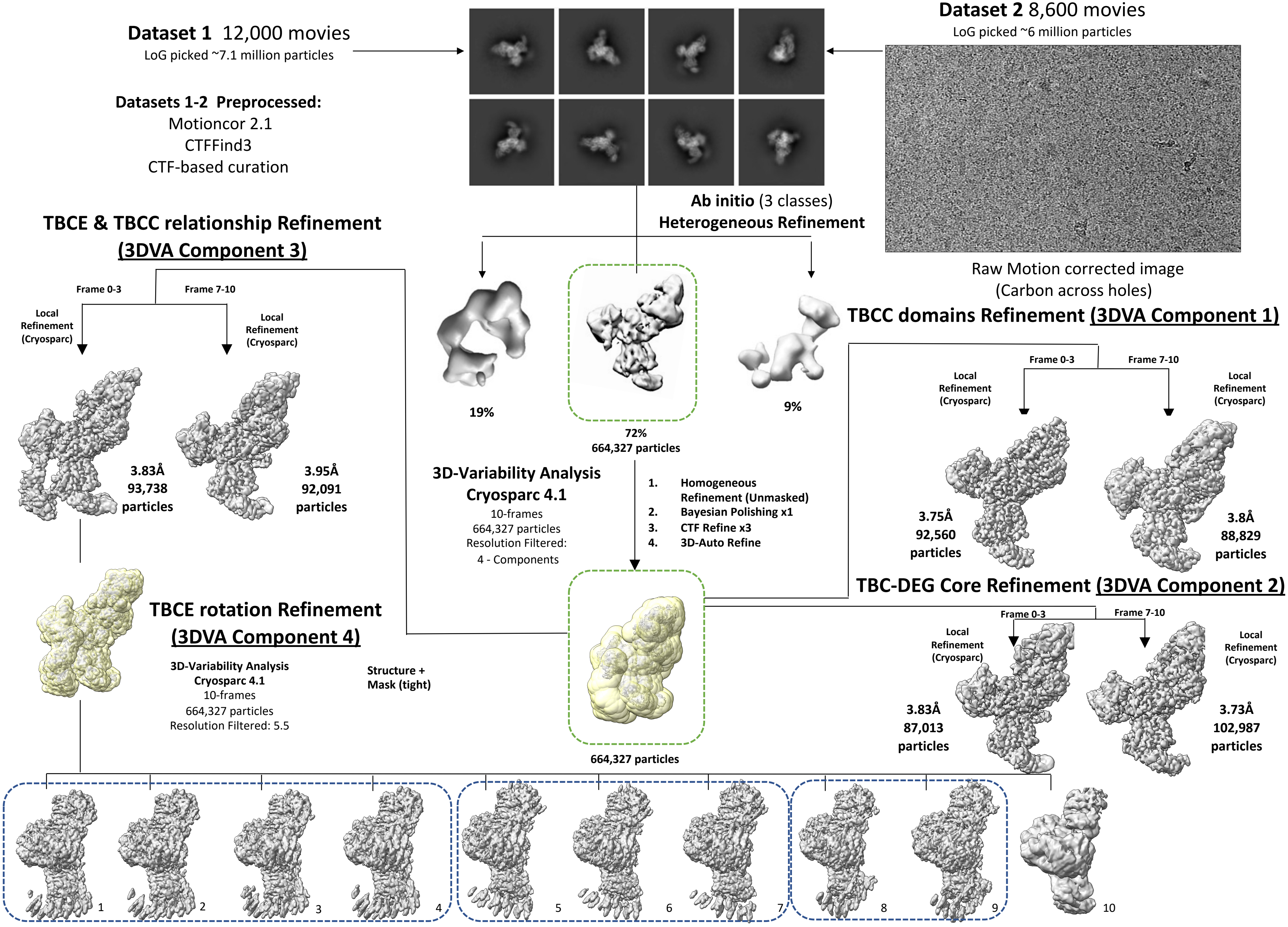

Figure S7

TBC-DEG/TBCC-αβ-tubulin 3D variability analyses / maps and map comparisons

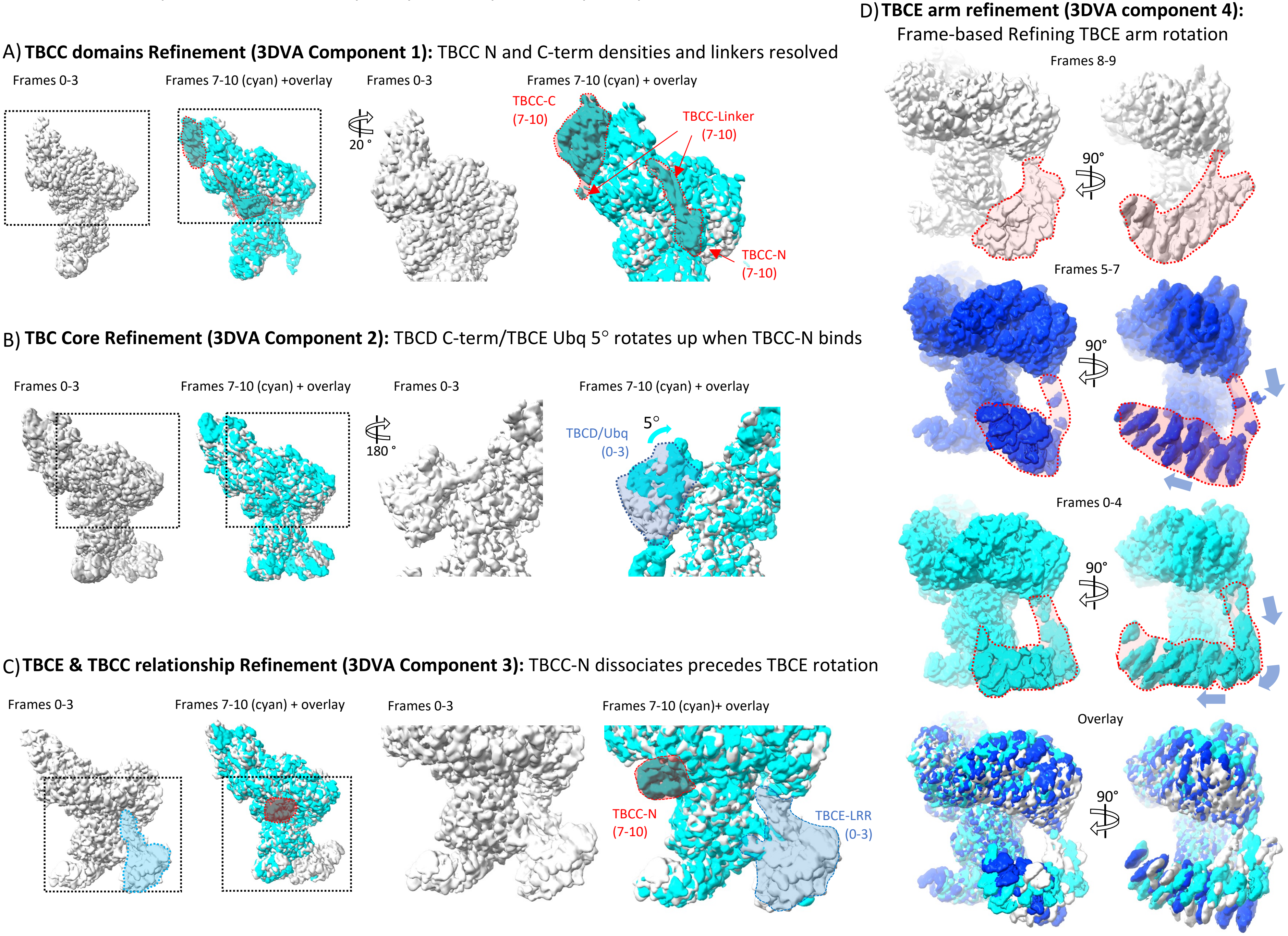

Figure S8

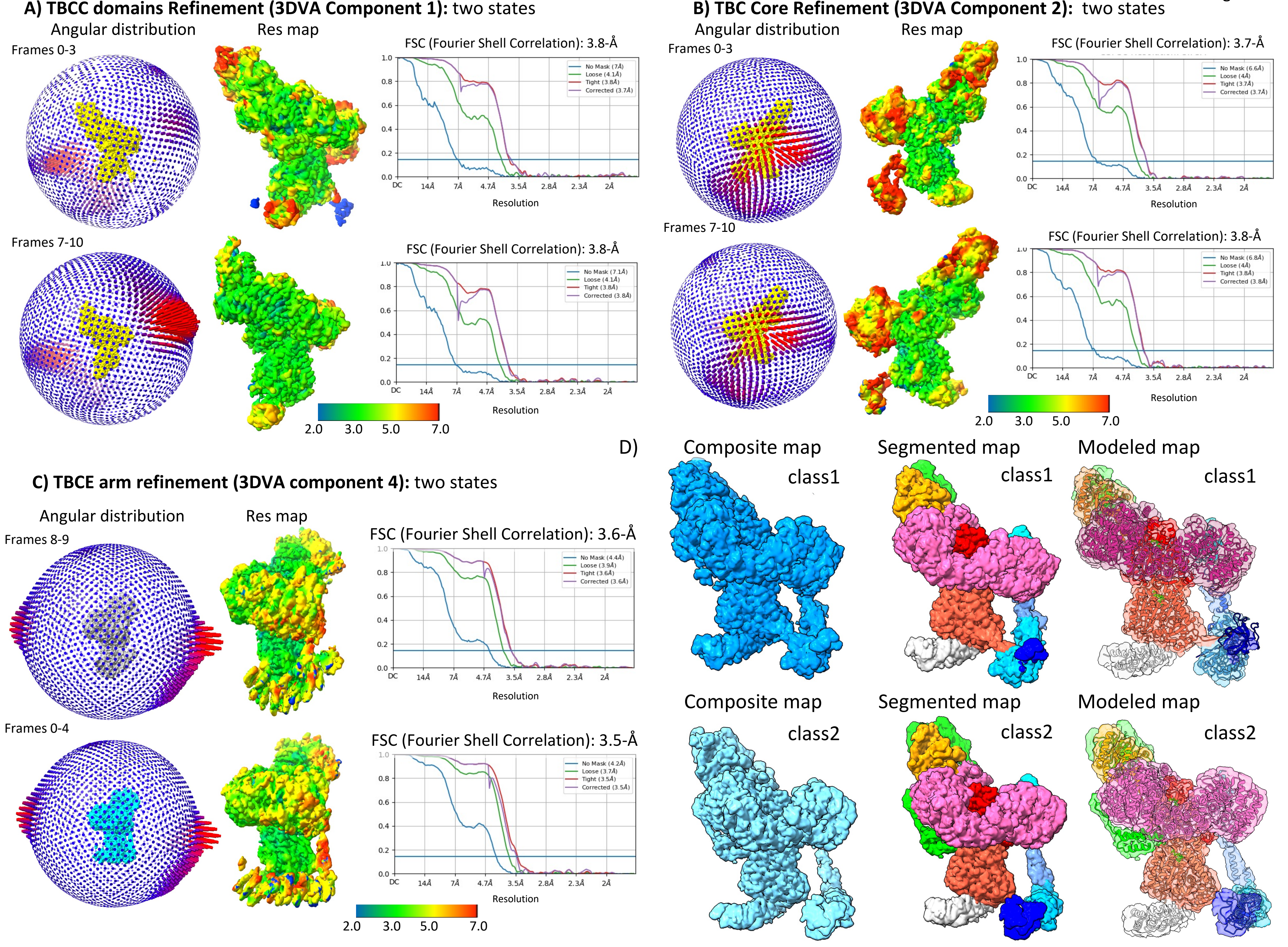

A) TBCC domains Refinement (3DVA Component 1): two states
Angular distribution
Res map
Frames 0-3
FSC (Fourier Shell Correlation): 3.8-Å
Resolution
Frames 7-10
FSC (Fourier Shell Correlation): 3.8-Å
Resolution
2.0 3.0 5.0 7.0
B) TBC Core Refinement (3DVA Component 2): two states
Angular distribution
Res map
Frames 0-3
FSC (Fourier Shell Correlation): 3.7-Å
Resolution
Frames 7-10
FSC (Fourier Shell Correlation): 3.8-Å
Resolution
2.0 3.0 5.0 7.0
C) TBCE arm refinement (3DVA component 4): two states
Angular distribution
Res map
Frames 8-9
FSC (Fourier Shell Correlation): 3.6-Å
Resolution
Frames 0-4
FSC (Fourier Shell Correlation): 3.5-Å
Resolution
2.0 3.0 5.0 7.0
D)
Composite map
class1
Segmented map
class1
Modeled map
class1
Composite map
class2
Segmented map
class2
Modeled map
class2

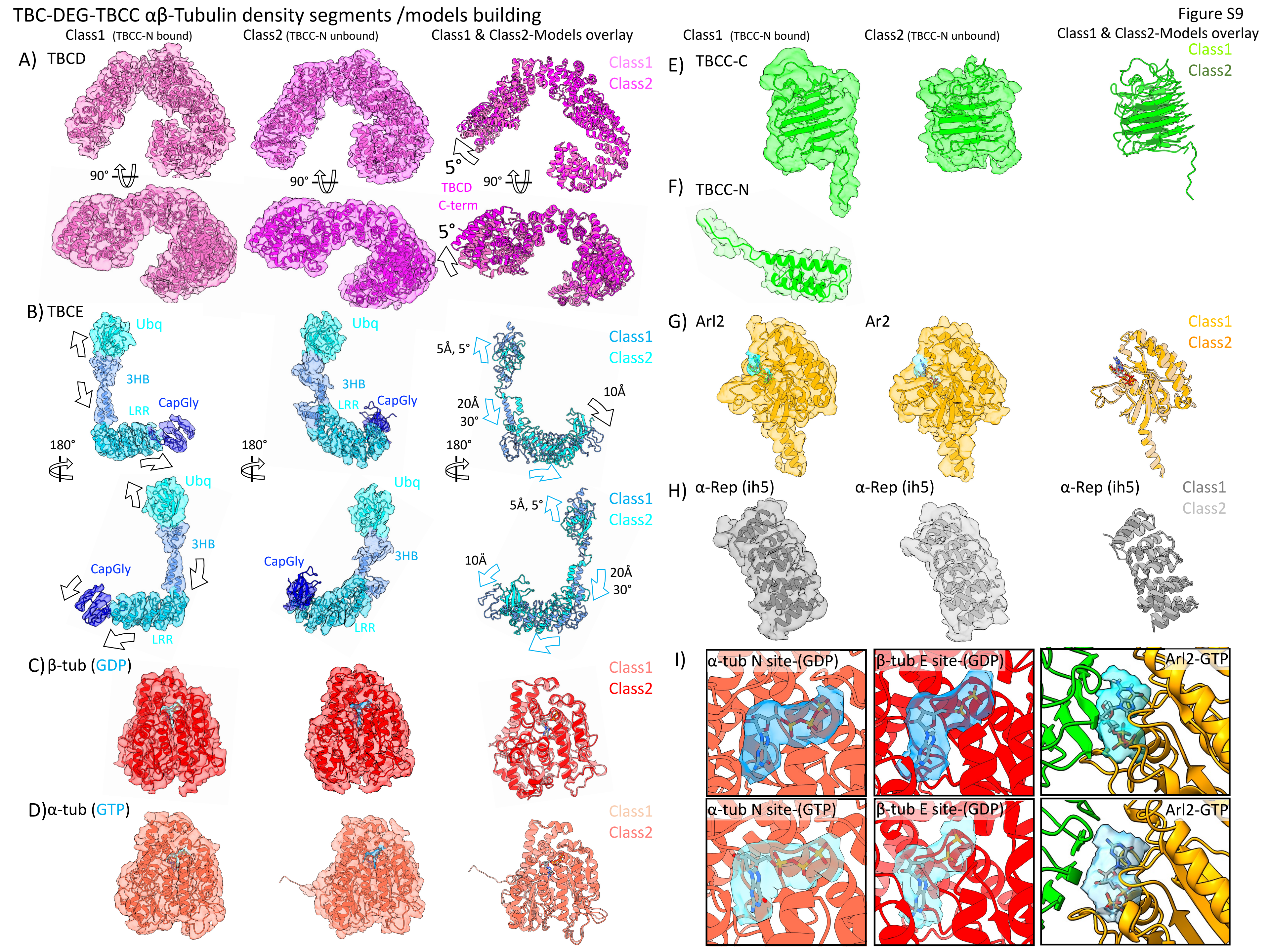

TBC-DEG-TBCC αβ-Tubulin density segments /models building
Figure S9
Class1 (TBCC-N bound)
Class2 (TBCC-N unbound)
Class1 & Class2-Models overlay
A) TBCD
Class1
Class2
90°
5°
TBCD
C-term
B) TBCE
Ubq
3HB
LRR
CapGly
180°
5Å, 5°
20Å
30°
10Å
C) β-tub (GDP)
D) α-tub (GTP)
E) TBCC-C
F) TBCC-N
G) Arl2
Ar2
H) α-Rep (ih5)
I)
α-tub N site-(GDP)
β-tub E site-(GDP)
Arl2-GTP
α-tub N site-(GTP)
β-tub E site-(GDP)
Arl2-GTP

TBC-DEG-TBCC αβ-Tubulin conformational changes: detailed models for Class1 versus class in response and impact of TBCC_N binding

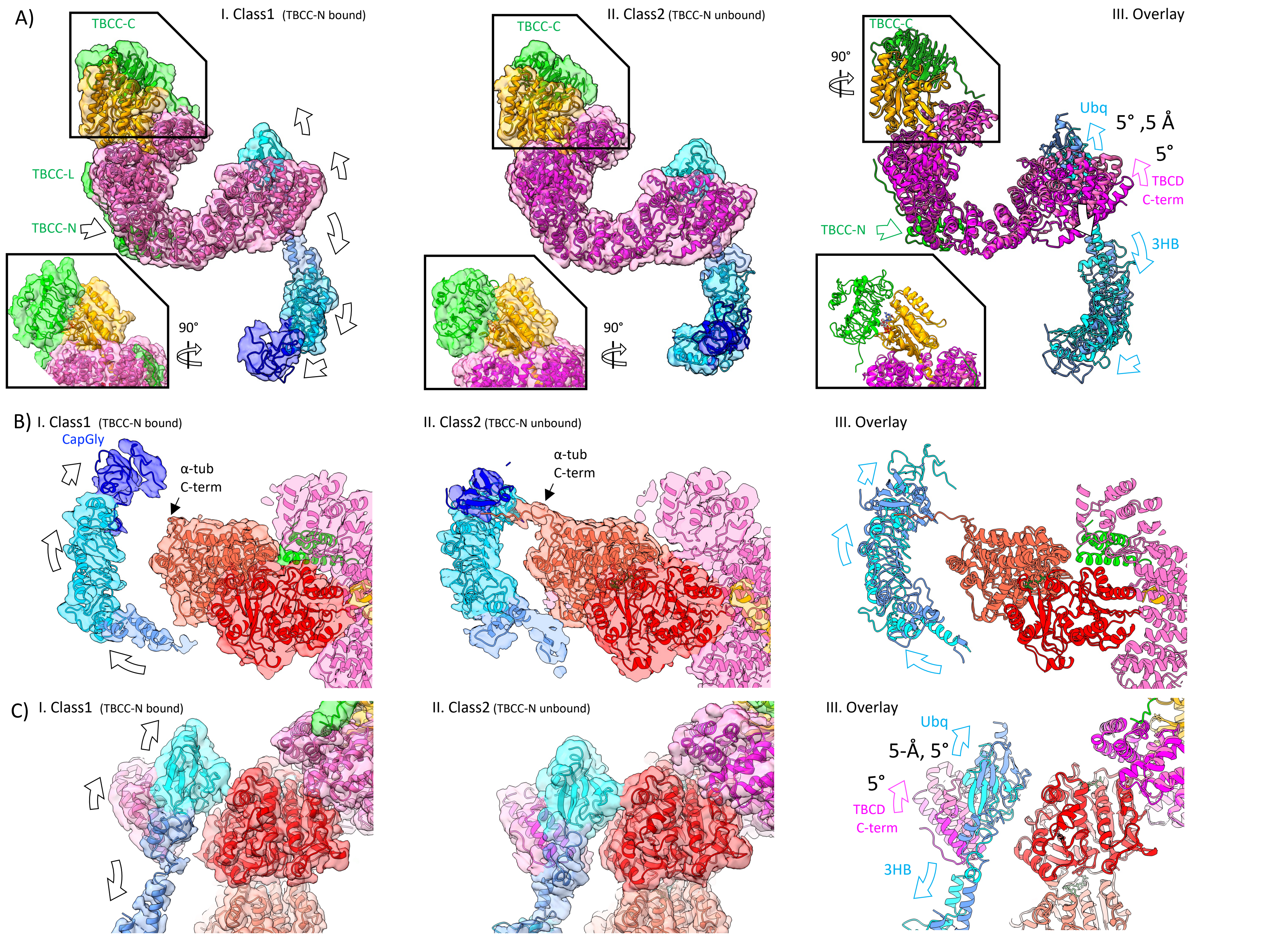

TBCD alignment    Figure S11

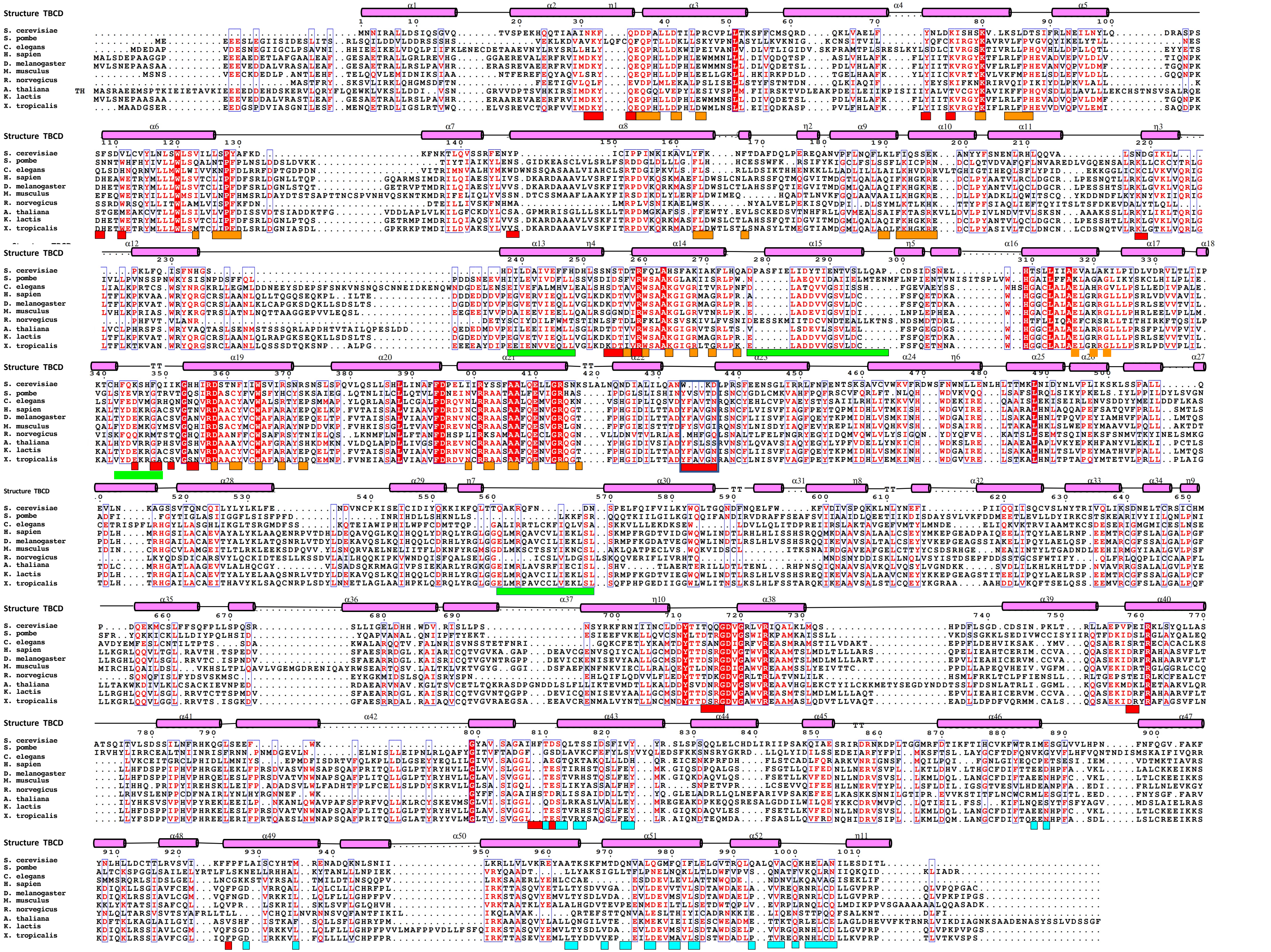

Figure S12

A) TBCE alignment β-tubulin binding TBCD binding

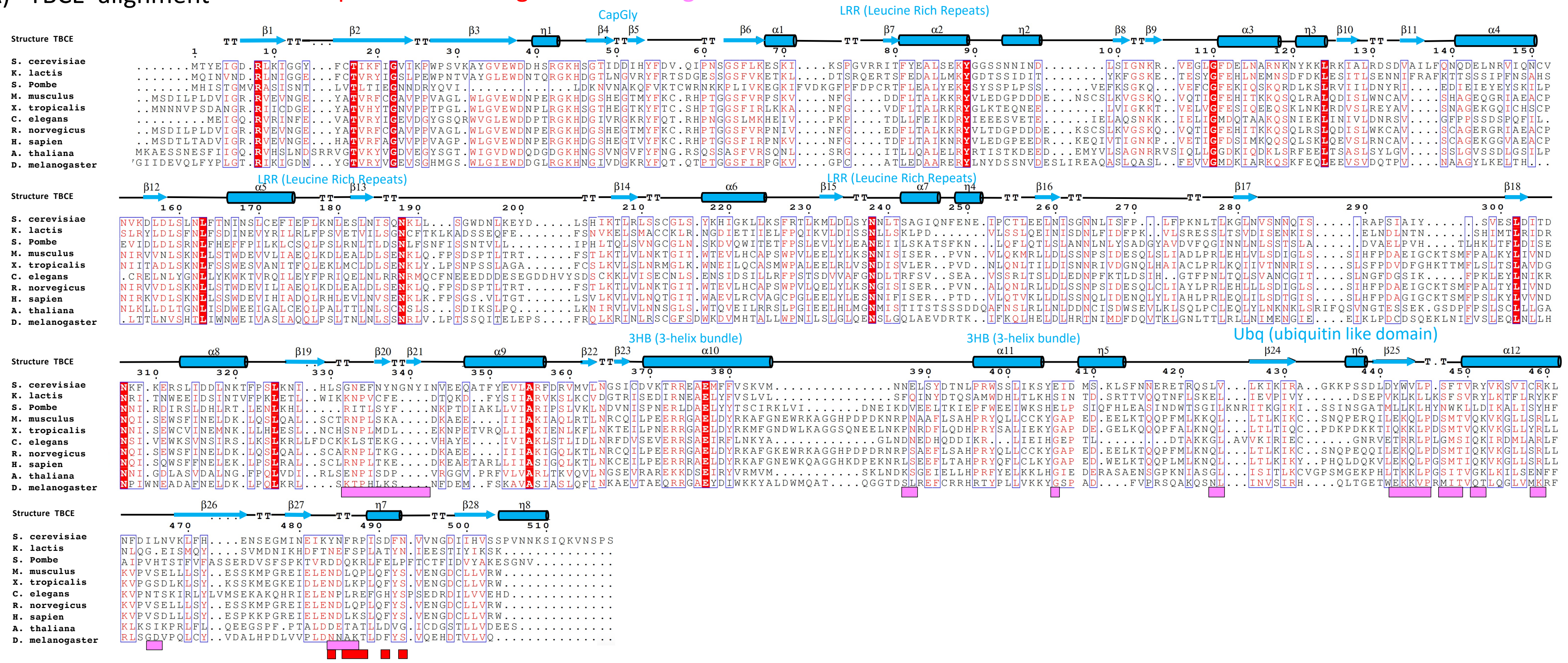


B) Arl2 alignment β-tubulin binding TBCD binding TBCC-C binding

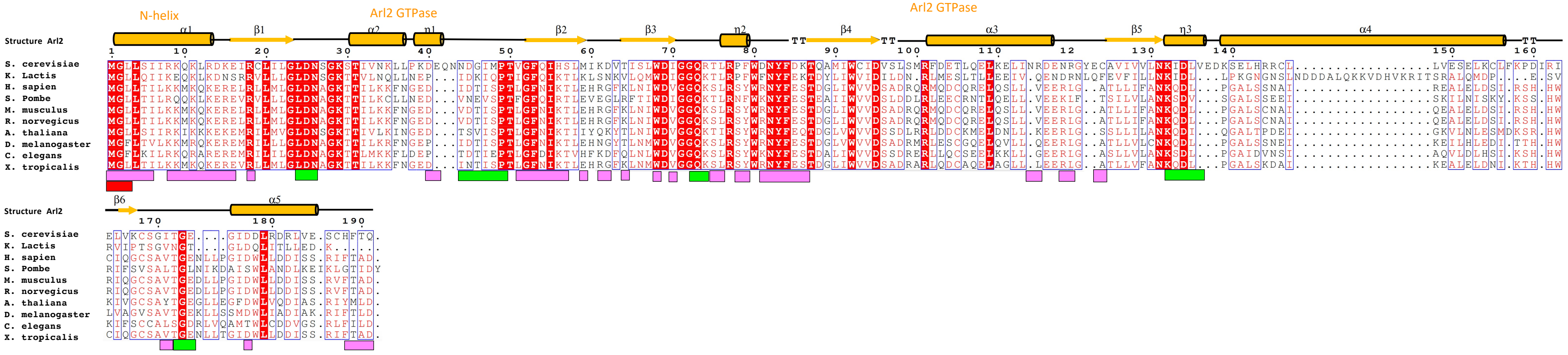

Figure S13

A) TBCC alignment α-tubulin binding β-tubulin binding TBCD binding Arl2 binding

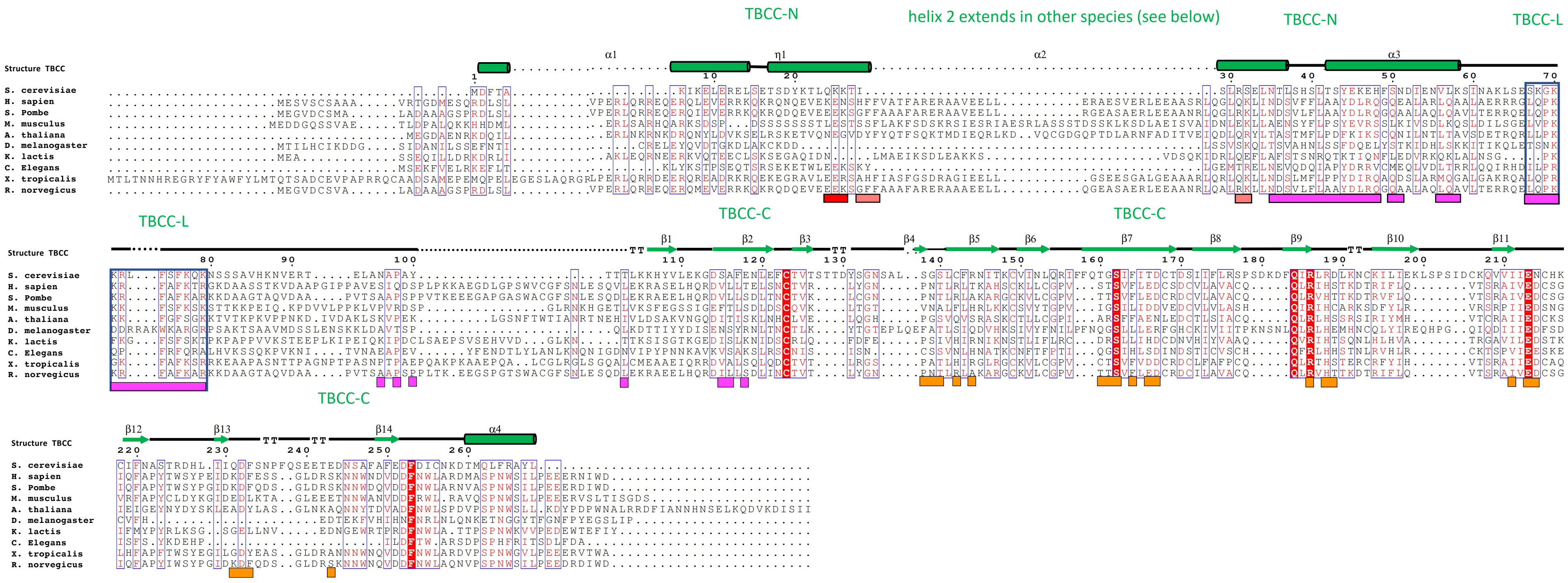


B) Alpha fold models for TBCC orthologs from divergent species demonstrates variation in TBCC-N first helix length

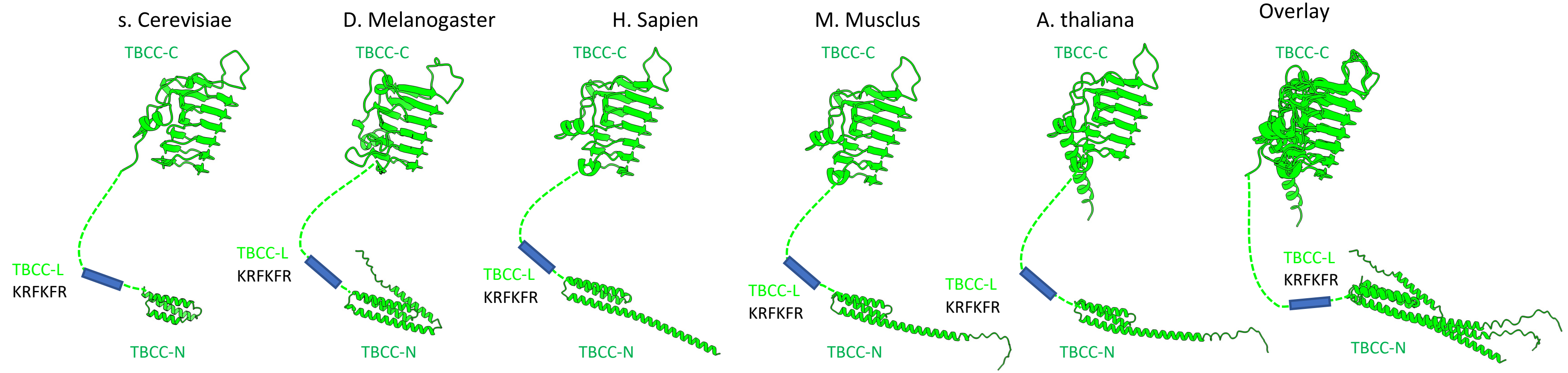

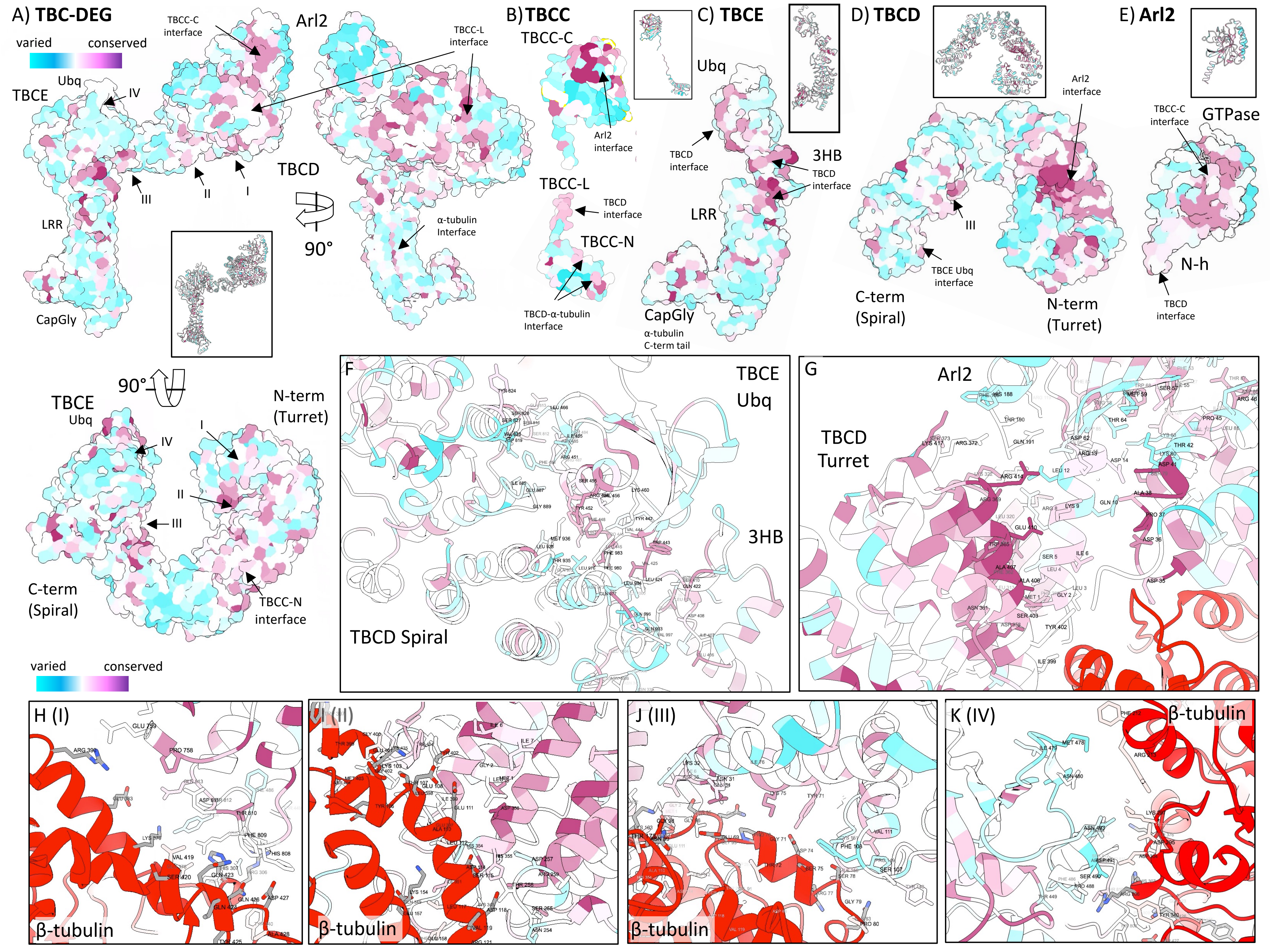
A) TBC-DEG
varied
conserved
TBCC-C interface
Arl2
TBCC-L interface
TBCE
Ubq
IV
III
II
I
TBCD
90°
LRR
α-tubulin Interface
CapGly
B) TBCC
TBCC-C
Arl2 interface
TBCC-L
TBCD interface
TBCC-N
TBCD-α-tubulin Interface
C) TBCE
Ubq
TBCD interface
3HB
TBCD interface
LRR
CapGly
α-tubulin C-term tail
D) TBCD
Arl2 interface
III
TBCE Ubq interface
C-term (Spiral)
N-term (Turret)
E) Arl2
TBCC-C interface
GTPase
N-h
TBCD interface
90°
TBCE
Ubq
IV
N-term (Turret)
I
II
III
C-term (Spiral)
TBCC-N interface
varied
conserved
F
TBCE Ubq
3HB
TBCD Spiral
G
Arl2
TBCD Turret
H (I)
β-tubulin
I (II)
β-tubulin
J (III)
β-tubulin
K (IV)
β-tubulin

#### The TBC-DEG/ TBCC subunit N and C termini orientation (Rainbow colored subunits)

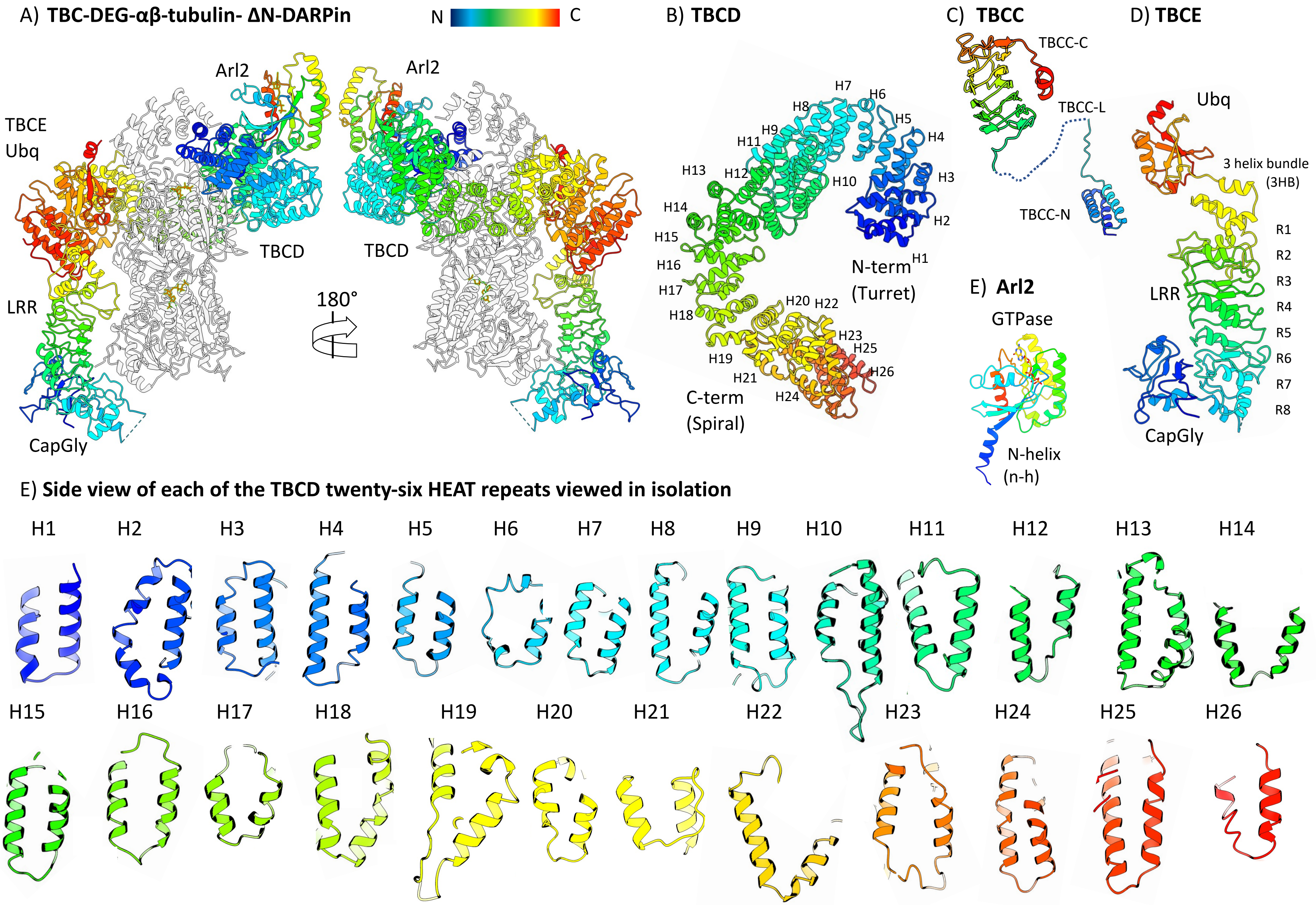

Figure S16

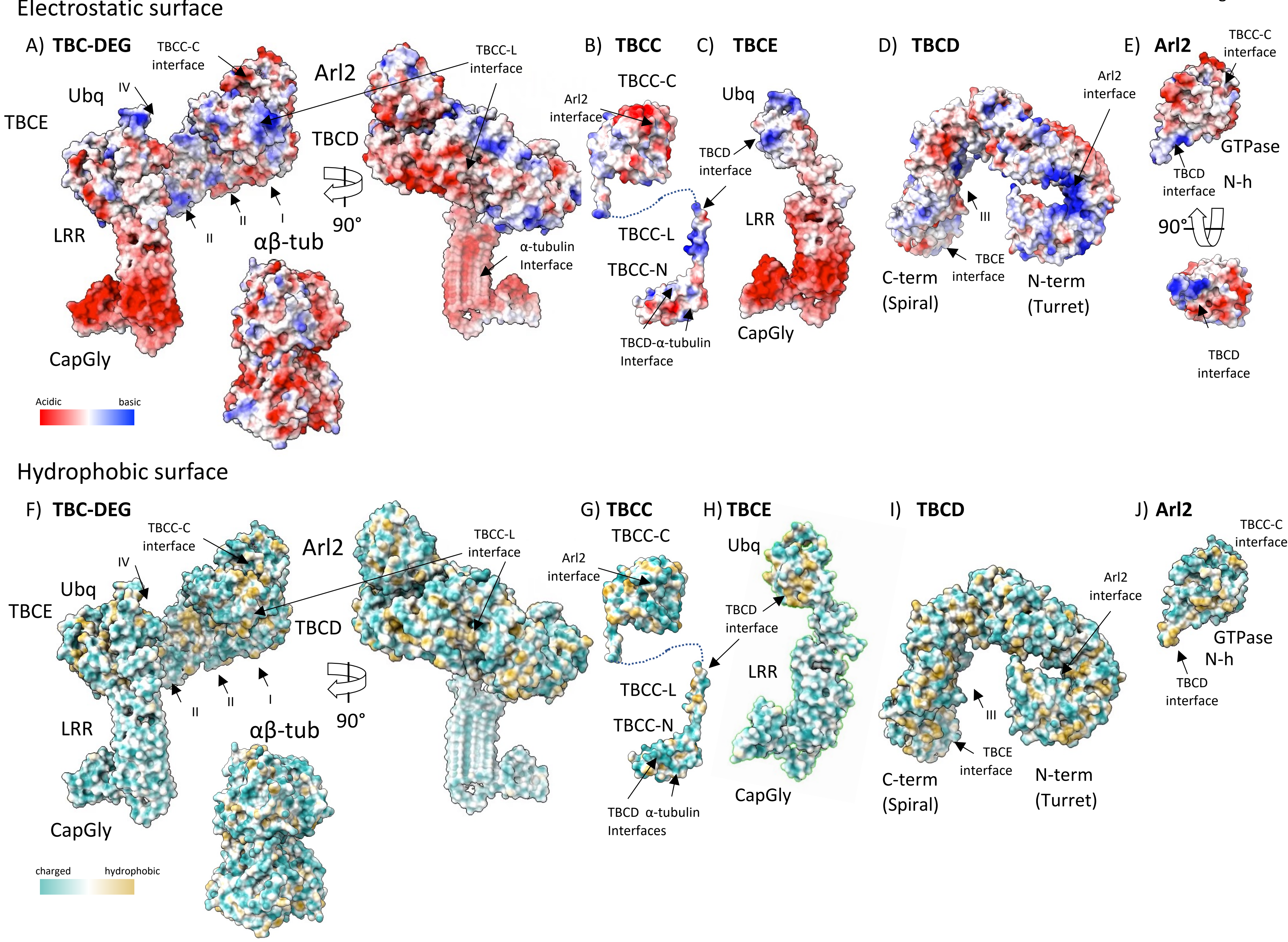

Electrostatic surface
A) TBC-DEG
TBCC-C interface
Arl2
TBCC-L interface
Ubq
IV
TBCE
TBCD
LRR
II
II
I
90°
αβ-tub
α-tubulin Interface
CapGly
Acidic
basic
B) TBCC
TBCC-C
Arl2 interface
TBCC-L
TBCC-N
TBCD-α-tubulin Interface
C) TBCE
Ubq
TBCD interface
LRR
CapGly
D) TBCD
Arl2 interface
III
TBCE interface
C-term (Spiral)
N-term (Turret)
E) Arl2
TBCC-C interface
GTPase
TBCD interface
N-h
90°
TBCD interface
Hydrophobic surface
F) TBC-DEG
TBCC-C interface
Arl2
TBCC-L interface
IV
Ubq
TBCE
TBCD
II
II
I
90°
LRR
αβ-tub
CapGly
charged
hydrophobic
G) TBCC
TBCC-C
Arl2 interface
TBCC-L
TBCC-N
TBCD α-tubulin Interfaces
H) TBCE
Ubq
TBCD interface
LRR
CapGly
I) TBCD
Arl2 interface
III
TBCE interface
C-term (Spiral)
N-term (Turret)
J) Arl2
TBCC-C interface
GTPase
N-h
TBCD interface
